# Structural Diversity of Photoswitchable Sphingolipids for Optodynamic Control of Lipid Raft Microdomains

**DOI:** 10.1101/2021.10.11.463883

**Authors:** Nina Hartrampf, Samuel M. Leitao, Nils Winter, Henry Toombs-Ruane, James A. Frank, Petra Schwille, Dirk Trauner, Henri G. Franquelim

**Affiliations:** Department of Chemistry, Ludwig Maximilian University of Munich, Butenandtstraße 5-13, Munich 81377, Germany; Department of Chemistry, University of Zurich, Winterthurerstrasse 190, Zurich 8057, Switzerland; Cellular and Molecular Biophysics Department, Max Planck Institute of Biochemistry, Am Klopferspitz 18, Martinsried 82152, Germany; Institute for Bioengineering, École Polytechnique Fédérale de Lausanne, Bâtiment BM 3109 Station 17, Lausanne 1015, Switzerland; Vollum Institute, Oregon Health & Science University, 3181 SW Sam Jackson Park Road, Portland, Oregon 97239, United States; Department of Chemistry, New York University, 100 Washington Square East, New York City, New York 10003, United States

**Author notes:** Co-authors with equal contribution (Nina Hartrampf, Samuel M. Leitao). Corresponding authors: HGF & DT.

## Abstract

Sphingolipids are a structurally diverse class of lipids predominantly found in the plasma membrane of eukaryotic cells. These lipids can laterally segregate with other saturated lipids and cholesterol into lipid rafts; liquid-ordered (L_o_) microdomains that act as organizing centers within biomembranes. Owing the vital role of sphingolipids for lipid segregation, controlling their lateral localization is of utmost significance. Hence, we made use of the light-induced *trans-cis* isomerization of azobenzene-modified acyl chains, to develop a set of photoswitchable sphingolipids, with different headgroups (hydroxyl, galactosyl, phosphocholine) and backbones (sphingosine, phytosphingosine, tetrahydropyran (THP)-blocked sphingosine), able to shuttle between liquid-ordered (L_o_) and liquid-disordered (L_d_) regions of model membranes upon irradiation with UV-A (λ = 365 nm) and blue (λ = 470 nm) light, respectively. Using combined high-speed atomic force microscopy, fluorescence microscopy, and force spectroscopy, we investigated how these active sphingolipids laterally remodel supported bilayers upon photo-isomerization, notably in terms of domain area changes, height mismatch, line tension, and membrane piercing. Hereby, we show that all sphingosine-(Azo-β-Gal-Cer, Azo-SM, Azo-Cer) and phytosphingosine-based (Azo-α-Gal-PhCer, Azo-PhCer) photolipids behave similarly, promoting a reduction in L_o_ domain area when in the UV-adapted *cis*-isoform. In contrast, azo-sphingolipids having THP groups that block H-bonding at the sphingosine backbone (Azo-THP-SM, Azo-THP-Cer) induce an increase in the L_o_ domain area when in *cis*, accompanied by a major rise in height mismatch and line tension. These changes were fully reversible upon blue light-triggered isomerization of the various lipids back to *trans*, pinpointing the role of interfacial interactions for the formation of stable L_o_ lipid raft domains.

## 1. Introduction

Sphingolipids are a major component of eukaryotic (notably mammalian) membranes and play a crucial role in signaling inside cells^[1–3]^. Members of this lipid class, like ceramide (Cer) or sphingomyelin (SM), have a backbone formed from sphingosine, and have been widely studied in terms of their biophysical properties, behavior on membrane models, and affinity to other lipids^[4–11]^. Another important, but less studied, class of sphingolipids are phytosphingolipids, which are abundant in plants and fungi^[12–14]^. These lipids have phytosphingosine as sphingoid base^[2]^, a backbone with increased polarity in comparison to sphingosine. Sphingolipids are also largely localized in the plasma membranes of eukaryotic cells, when compared to inner organelle membranes^[3]^. Here, they are usually linked to so-called lipid rafts^[15]^, liquid ordered (L_o_) phase nano- or microdomains composed of saturated lipid species and sterols, thought to be a means by which cells organize or segregate important proteins within the membrane^[16–18]^.

From a molecular point of view, sphingolipids can form stable hydrogen-bond and hydrophobic interactions with other sphingolipids (e.g. SM) and cholesterol (Chol)^[2,19]^. This can, e.g., be observed *in vitro* on supported membrane model systems, as segregated rigid L_o_ phase microdomains embedded in a fluid bulk liquid disordered (L_d_) membrane phase^[20–24]^. The presence of hydroxyl (-OH) groups on the backbone of sphingolipids are particularly relevant for the formation of L_o_ domains, as interfacial H-bonding markedly stabilize the interactions among sphingolipids and Chol^[19,25]^. In fact, the central role of the 3-OH moiety on the sphingosine backbone of sphingolipids, like Cer or SM, has been thoroughly scrutinized^[13,26–32]^. Likewise, the presence of a second 4-OH hydroxyl group on phytosphingosine-based lipids, like PhCer, further strengthens H-bonding and domain thermostability^[13]^. In contrast, hindering H-bonding by adding a methyl, ethyl or tetrahydropyranyl (THP) group at the 3-OH hydroxyl, severely affected the molecular packing and ability of those blocked sphingolipids to interact with Chol^[26,33]^. Indeed, functionalization of the 3-OH of SM by THP greatly decreased gel-phase stability (lowering the melting temperature (T_m_) by 10°C), impede tight contacts with Chol, and increase the rate of sterol desorption from vesicles containing this blocked SM analog^[26]^.

As lipid segregation plays a crucial physiological role in biomembranes, new nano-tools to investigate and control membrane phase properties are urgently needed. In that regard, strategies based on photopharmacology^[34,35]^, which take advantage of the high spatiotemporal precision of light, are particularly appealing. In 2016, we reported that photoswitchable ceramides (ACes), which have an azobenzene photoswitch incorporated in the lipid fatty acid chain, enable optical control of lipid rafts within synthetic membranes.^[36]^ While recent advancements already demonstrated the potential of azobenzene-modified photoswitchable lipids for altering membrane properties ^[36–43]^, the structural diversity of these photo-active molecules is still fairly limited. This stands in stark contrast with the impressive diversity of sphingolipids found in nature. Indeed, diverse sphingolipids can serve as docking site for various toxins (e.g. Shiga toxin binds to Gb3^[67]^ or Cholera toxin B binds to GM1^[44]^), or even modulate the uptake of viruses by the cells (e.g. SV40 requires GM1 as receptor^[45]^ or HIV-1 gp120 surface protein binds to GalCer on epithelial cells^[46]^). Hence, controlling their lateral localization within membranes is of vital importance. An expanded palette of photoswitchable sphingolipids could therefore offer new photo-responsive *N*-acyl azobenzene sphingolipids with more complex headgroup functionalities and other types of sphingoid backbones.

In our present work we then aimed to develop a new set of photoswitchable sphingolipids with increased functionalization and incorporate these various photolipids into L_d_--L_o_ phase-separated supported model membranes. Our main goal was to investigate the influence of these modifications on the reversible remodeling of membranes microdomains upon photoswitching, as well as on fundamental mechanical properties of lipid bilayers. To this end, we performed atomic force microscopy (AFM) combined with fluorescence confocal microscopy, following the generated changes in domain area, height and line tension.

## 2. Results and Discussion

### 2.1. Synthesis of photoswitchable sphingolipids

Inspired by the structural design of our simpler azo-ceramides (ACes)^[36]^ and more complex α-galactosyl-phytoceramides (α-GalACers)^[37]^, we introduced five new azobenzene-modified sphingolipids, namely **Azo-PhCer**, **Azo-THP-Cer**, **Azo-β-Gal-Cer**, **Azo-SM** and **Azo-THP-SM**. These photolipids featured (Fig. 1A): 1) a FAAzo-4 fatty acid^[34]^ at the *N*-acyl chain (equivalent to a Δ9 unsaturation when in the *cis*-isoform), 2) sphingoid backbones based on naturally-occurring sphingosine and phytosphingosine, or hydroxyl-blocked sphingosine with a THP protecting group, as well as 3) lipid headgroups presenting either a free -OH, galactosyl or phosphocholine moiety. For our comparative studies, **Azo-Cer** (previously named ACe-1)^[36]^ and **Azo-α-Gal-PhCer**^[37]^ (previously named GalACer-4), having the same FAAzo-4 moiety able to undergo *trans-cis* photoisomerization (Fig. 1B), were also assessed.

**Fig. 1.**
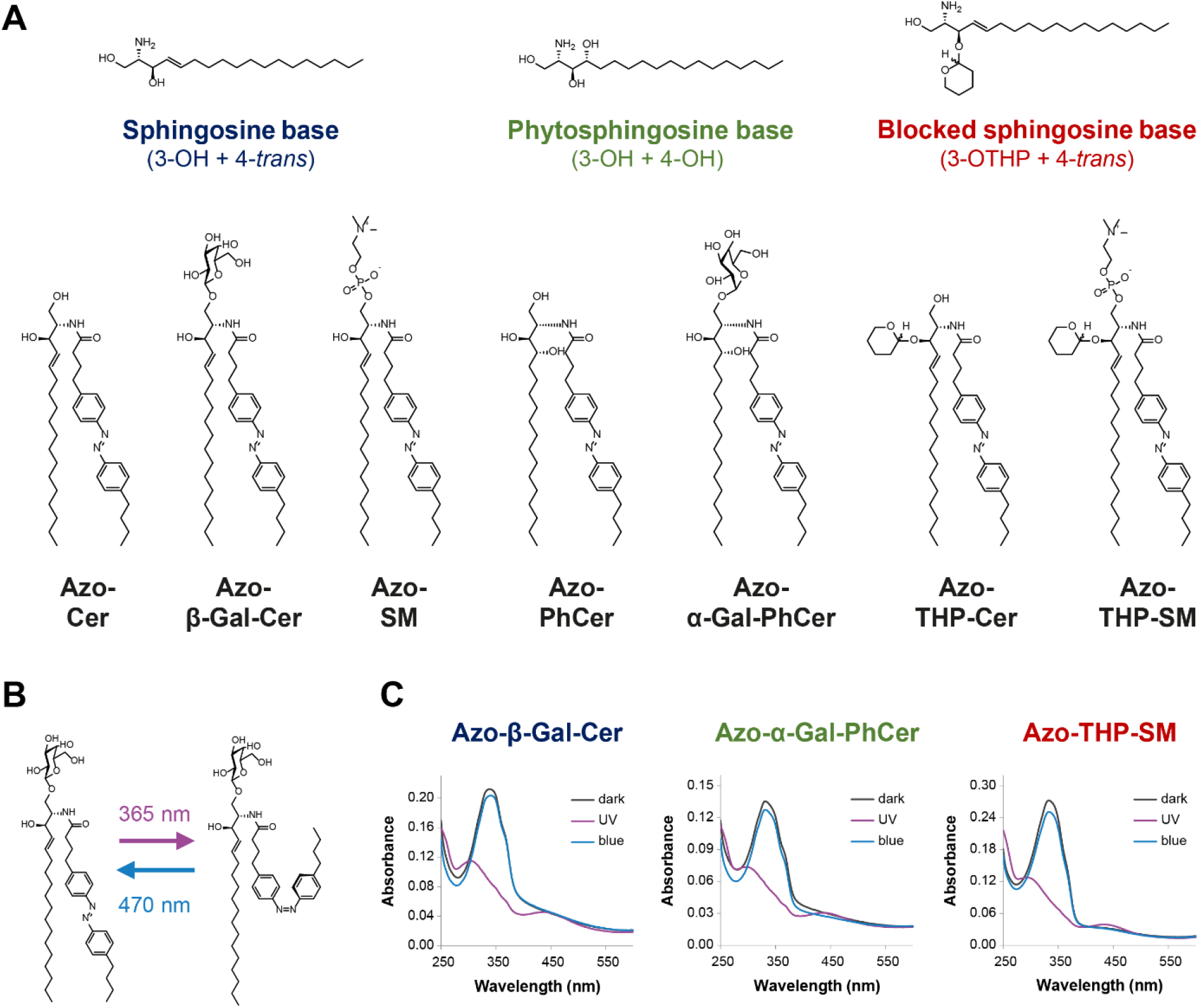
Structural and spectral properties of photoswitchable azo-sphingolipids. (A) chemical structures of the N-acyl azobenzene-modified (FAazo-4 fatty acid) sphingolipids here tested, subdivided according to their sphingoid backbone: Azo-Cer, Azo-β-Gal-Cer and Azo-SM with a sphingosine base, Azo-PhCer and Azo-α-Gal-PhCer with a phytosphingosine base, as well as Azo-THP-Cer and Azo-THP-SM displaying a 3-OH-blocked sphingosine base with a THP protecting group. (B) Schematics of light-induced trans-cis isomerization for an azo-sphingolipid, notably Azo-β-Gal-Cer. Application of UV-A light (λ = 365 nm) leads to the formation of cis-photolipid, while illumination with blue light (λ = 470 nm) leads to the formation of trans-photolipid. (C) UV-Vis absorbance spectra of photoswitchable sphingolipids (notably Azo-β-Gal-Cer, Azo-α-Gal-PhCer and Azo-THP-SM) incorporated in SUVs, at the dark-adapted state (black curves), as well as after the sequential shining of UV-A (purple curves) and blue light (blue curves).

The synthesis and characterization of **Azo-Cer** and **Azo-α-Gal-PhCer** were reported elsewhere^[36,37]^. **Azo-PhCer** was prepared analogously to **Azo-Cer** by the coupling of phytoceramide with FAAzo-4 using HBTU as a coupling agent (see SI). Additional protecting group manipulations yielded **Azo-THP-Cer**.

For the synthesis of **Azo-β-Gal-Cer**, we used a benzoyl protected alcohol and the azide as protecting groups^[47]^ (see SI). Azides do not coordinate to the primary alcohol and thereby the nucleophilicity of the sphingosine is greatly enhanced. Glycosylation of azidosphingosine with trichloroacetimidate yielded protected glycoside in 92% yield and excellent β-selectivity. Staudinger reduction using PBu_3_ and subsequent amide coupling with FAAzo-4^[34]^ using 1-ethyl-3-(3-dimethylaminopropyl)carbodiimide (EDCI), followed by global deprotection gave **Azo-β-Gal-Cer** (see SI).

The sphingomyelin derivatives were prepared from **Azo-THP-Cer**, which was phosphorylated using 2-cyanoethyl-*N*,*N*,*N*’,*N’*-tetraisopropylphosphorodiamidite and 1H-tetrazole, followed by reaction with choline tosylate (see SI). An oxidation directly yielded **Azo-THP-SM**. Finally, deprotection under acidic conditions gave the unprotected **Azo-SM** (see SI).

### 2.2. Light-responsiveness of membrane-embedded azo-sphingolipids

Next, we incorporated our newly synthesized **Azo-β-Gal-Cer**, **Azo-SM**, **Azo-PhCer**, **Azo-THP-SM** and **Azo-THP-Cer** photoswitchable lipids (or simply *photolipids*), as well as **Azo-Cer** and **Azo-α-Gal-PhCer**, into raft-mimicking L_d_-L_o_ phase-separated mixtures containing DOPC, Chol and SM (18:0-SM) and formed small unilamellar vesicles (SUVs) as previously described^[36,48]^. Unless otherwise stated, a molar ratio of 10:6.7:5:5 (DOPC:Chol:SM:*photolipid*) was typically chosen.

We started by collecting UV-Vis spectra of those various SUV suspensions and characterized the photodynamic properties of the different azobenzene-modified sphingolipids within a membrane environment. As seen in Figs. 1C and S1 (see SI), all azo-sphingolipids (independently of the headgroup and backbone type) displayed an absorbance maximum, λ_max_, at ~ 350 nm, when in the dark-adapted state prior irradiation with UV-A or blue light. This peak corresponds to the π → π* transition and is characteristic for the *trans*-azobenzene isoform. First illumination with UV-A light (λ = 365 nm) led to the reduction of the abovementioned absorbance peak, and appearance of a new λ_max_ at ~ 450 nm. This new peak corresponds to the n → π* transition and is characteristic for the *cis*-azobenzene isoform. Subsequent irradiation with blue light (λ = 470 nm) led then to a back-isomerization of the *N*-acyl azobenzene moieties into the *trans*-isoform, as confirmed by the disappearance of the absorbance peak at ~ 450 nm and reemergence of the absorbance peak at ~ 350 nm.

Our results indicate that all tested photoswitchable sphingolipids are in the *trans*-configuration for the dark- and blue light-adapted states, and mostly in *cis*-configuration for the UV light-adapted state. Similarly, no significant spectral shifts from the characteristic 350 nm and 450 nm excitation peaks were observed, pointing out that headgroup functionality and sphingoid base polarity do not critically interfere with the photodynamic properties of the photoswitch.

Subsequently, we deposited SUVs composed of quaternary DOPC:Chol:SM:*photolipid* mixtures doped with 0.1 mol% Atto655-DOPE (dye labelling the L_d_ regions) on top of freshly-cleaved mica, to form supported lipid bilayers (SLBs) via vesicle fusion. By collecting fluorescence confocal images, we first evaluated membrane integrity and presence of phase-separation for samples having sphingosine-(Azo-Cer, Azo-β-GalCer, Azo-SM), phytosphingosine-(Azo-PhCer, Azo-α-GalPhCer) and blocked THP-sphingosine-based (Azo-THP-SM) lipids. As seen in Figs. S2 and S3, all tested SLBs displayed L_d_-L_o_ phase-separation at the dark-adapted state, with micron-sized rigid L_o_ domains (dark areas in fluorescence images) segregated within a fluid L_d_ matrix (red areas in fluorescence images).

The photo-responsiveness and ability of the azo-sphingolipids to then remodel/reorganize phase-separated SLBs were further assessed directly after irradiation with UV-A light (λ = 365 nm). For membranes lacking photoswitchable lipid (control sample with a DOPC:Chol:SM composition at molar ratio 10:6.7:10), no light-induced remodeling was observed (Fig. S2A). In contrast, for SLBs containing azo-sphingolipids, stark lipid rearrangement dependent on the amount of *photolipid* present was reported. Here, lipid bilayers with the highest amount of azo-sphingolipid tested (18.7 mol%; DOPC:Chol:SM:*photolipid* with molar ratio 10:6.7:5:*5*) showed strong reorganization of the L_o_ domains with admixing of fluorescently-marked L_d_ lipids and blurring of the domain boundaries directly after UV-A irradiation (Fig. S2B-G). Intermediate amounts of photoswitchable lipid (11.2 mol%; DOPC:Chol:SM:*photolipid* with molar ratio 10:6.7:7:*3*) led to a significantly lower domain remodeling activity. Finally, for SLBs with the lowest amount of azo-sphingolipid tested (3.7 mol%; DOPC:Chol:SM:*photolipid* with molar ratio 10:6.7:5:*1*) no clearly perceptible membrane reorganization, such as admixing of L_d_-L_o_ domains, was observed (Fig. S3).

### 2.3. Remodeling of membrane domains by sphingosine-based azo-sphingolipids

After the abovementioned initial characterization, we systematically investigated the light-induced remodeling of phase-separated DOPC:Chol:SM:*photolipid* supported membranes (at molar ratio 10:6.7:5:*5*, meaning 18.7 mol% *photolipid*) using high-speed AFM. This technique enables us to capture minor dynamic changes in membrane architecture very accurately, due to its exquisite sub-nm resolution. Herein, we started by analyzing bilayers containing the sphingosine-based photoswitchable lipids Azo-Cer, Azo-β-GalCer and Azo-SM.

At the dark-adapted state, those various membranes displayed a height mismatch between the L_d_ and L_o_ regions of 1.1 – 1.7 nm; similarly to what we reported before for phase-separated bilayers with Azo-Cer (ACes) lipids^[36]^.

Application of UV-A light (λ = 365 nm) to bilayers having Azo-Cer, Azo-β-GalCer or Azo-SM led then to the generation of L_d_ phase, with no major alteration of the L_d_-L_o_ domain height mismatch (*i.e.* minor increase of ~ 0.1 nm). As seen in Figs. 2A, S4B and Movies S1-S4, the UV-induced isomerization of the *N*-acyl chains from a straight *trans*-form into a kinked *cis*-form, promoted an apparent “fluidization” of the phase-separated membranes, with an overall decrease of the total L_o_ area by ~ 25% (area L_o_(UV) / area L_o_(dark) = 0.75 ± 0.14), as depicted in Fig. S5. Directly after irradiation with UV-A light, small L_d_ “lakes” were formed within the more rigid thicker L_o_ domains on phase-separated membranes containing Azo-Cer, Azo-β-GalCer or Azo-SM. The number of fluid L_d_ “lakes” then rapidly dropped, in order to reduce surface tension. While the majority of the smaller L_d_ “lakes” seem to vanish towards the outer fluid L_d_ matrix, few larger fluid L_d_ “lakes” remained trapped inside the rigid L_o_ domains, appearing to grow primarily via Ostwald ripening^[49]^.

**Fig. 2.**
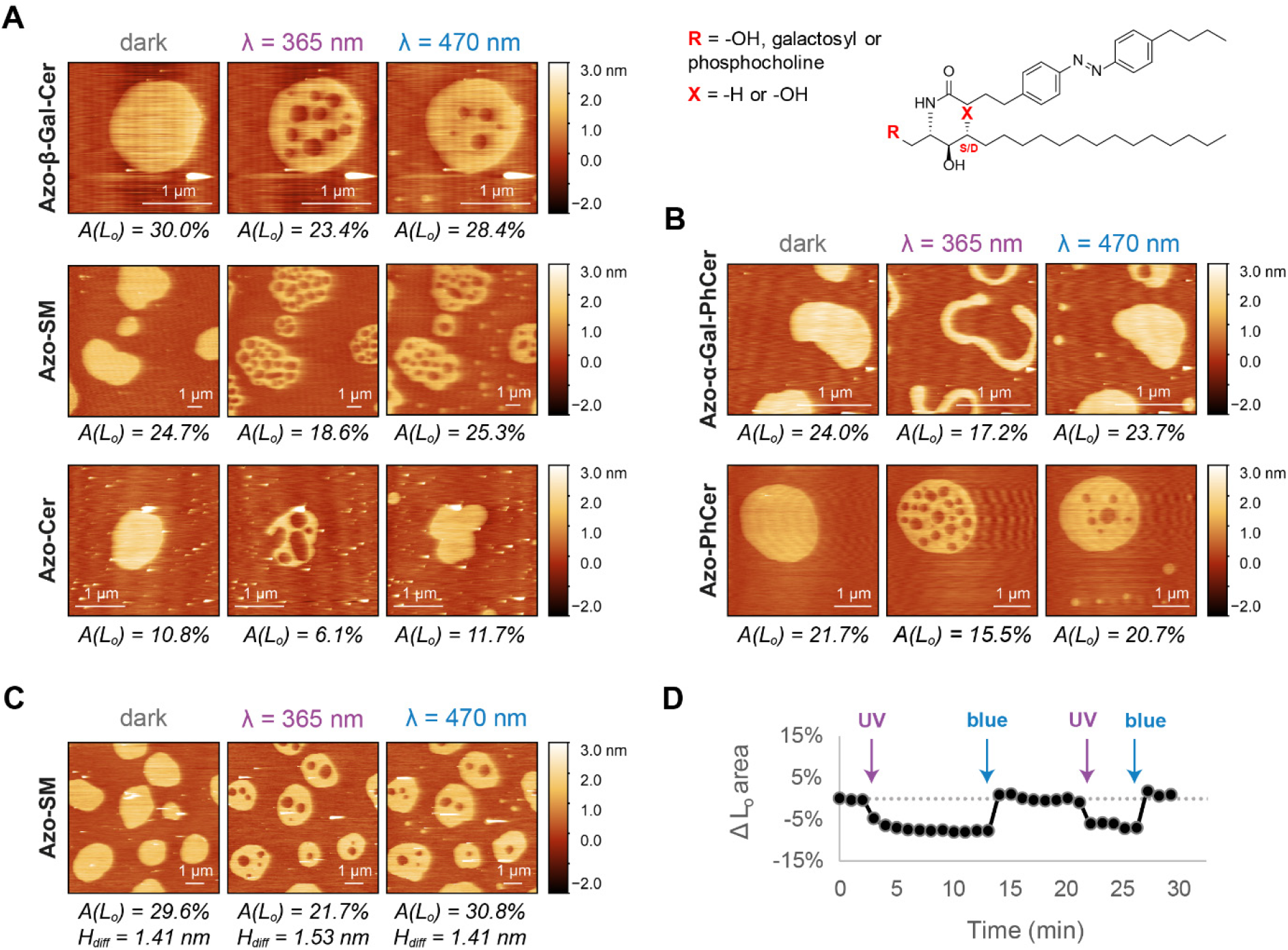
Lateral remodeling of phase-separated membranes containing non-blocked azo-sphingolipids upon light trigger analyzed by high-speed AFM. **(A-B)** Changes in the area of L_o_ domains before/after illumination with UV-A (λ = 365 nm) and blue (λ = 470 nm) lights on DOPC:Chol:SM:photolipid (10:6.7:5:5 mol ratio) SLBs having: (A) Azo-β-Gal-Cer, Azo-SM or Azo-Cer with a sphingosine backbone (X = -H) and varying headgroup functionality (R = galactosyl, phosphocholine or -OH, respectively); (B) Azo-α-Gal-PhCer or Azo-PhCer with a phytosphingosine backbone (X = -OH) and varying headgroup functionality (R = galactosyl or -OH, respectively). **(C-D)** Reversible lateral remodeling of a phase-separated SLB containing Azo-SM, upon UV-A/blue light irradiation, as seen in Movie S4: (C) AFM images of the SLB at the dark-, UV- and blue light-adapted states, displaying the area occupied by the L_o_ phase and the L_d_-L_o_ height mismatches. (D) Relative variation of total L_o_ area of the SLB over time, shown in Movie S4, upon shining short pulses (marked with arrows) of UV-A and blue light.

After few minutes of equilibration, subsequent application of a brief pulse of blue light (λ = 470 nm) to those phase-separated membranes reversed the effect, with L_o_ phase being generated. Here, back-isomerization of the *N*-acyl chains from a kinked *cis*-form into a straight *trans*-form, stimulated by blue light, promoted a rigidification of the membranes, with an increase of the total L_o_ area back to its original equilibrium dark-adapted value (area L_o_(blue) / area L_o_(dark) = 1.01 ± 0.18), as seen in Figs. 2A, S4B and S5. More specifically, upon irradiation with blue light, small rigid L_o_ “islands” were firstly formed within the fluid L_d_ matrix (Movies S1-S4). These taller L_o_ “islands” then vanished, as pre-existing L_o_ domains grew primarily via Ostwald ripening and domain fusion. Moreover, L_o_ domains displayed height values similar to the ones reported before for the initial dark-adapted state. Interestingly, those changes could be repeated over multiple cycles without dissipation effects, with the amount of L_d_-L_o_ phase-separation alternating between two defined steady-states (or area levels) (Fig. 2C–D).

Besides changes in L_d_-L_o_ phase-separation, we also observed sporadic generation of holes on our supported bilayers after the blue light-triggered conversion of the azo-sphingolipids’ *N*-acyl chains from *cis* to *trans* (Fig. S6A). The presence of holes allowed us to recover the total membrane thickness, which was ~ 5.2 nm (L_d_ thickness ~ 3.9 nm; Fig. S6B-C); in agreement with previously reported values for membranes of similar lipid composition^[50–52]^.

In summary, all the tested azo-sphingolipids with a sphingosine backbone display a similar photoswitching profile, independently of the type of headgroup. These lipids are able to increase the amount of L_d_ phase on phase-separated membranes upon conversion to the *cis*-isoform after UV-A light irradiation and increase the amount of L_o_ phase upon conversion to the *trans*-isoform after irradiation with blue light.

### 2.4. Remodeling of membrane domains by phytosphingosine-based azo-sphingolipids

Next, we recapitulated the same high-speed AFM procedures on membranes with photoswitchable phytosphingosine-based sphingolipids displaying two hydroxyl groups (3-OH + 4-OH) on the phytosphingosine backbone. More precisely, we investigated the ability of these photolipids to interfere with the L_d_-L_o_ phase-separation on supported lipid bilayers, when compared to sphingosine-based lipids having only one hydroxyl (3-OH) on their backbone.

In the dark-adapted state, phase-separated DOPC:Chol:SM:*photolipid* SLBs with Azo-PhCer and Azo-α-Gal-PhCer exhibited a domain height mismatch of 1.2 – 1.8 nm (Figs. 2B and S4C), very close to the L_d_-L_o_ height differences here reported for membranes with sphingosine analogs (Figs. 2A and S4B). Upon photoactivation, the L_d_-L_o_ phase-separated SLBs containing either Azo-PhCer (with a hydroxyl headgroup) and Azo-α-Gal-PhCer (with a bulkier galactosyl headgroup) behaved in a similar way to membranes with sphingosine-based azo-sphingolipids: exhibiting at the end an identical phenotype of membrane remodeling.

As seen in Figs. 2B, S4C and Movies S5-S6, after irradiation with UV-A light (λ = 365 nm), L_d_ “lakes” initially appeared inside existing L_o_ domains, and the total L_d_ phase membrane area lowered by ~ 23% (area L_o_(UV) / area L_o_(dark) = 0.77 ± 0.31, Fig. S5); while the domain height mismatch did not change majorly. Then, after irradiation with blue light (λ = 470 nm), L_o_ “islands” initially formed inside the L_d_ regions, and the total L_o_ phase membrane area subsequently increased to the initial equilibrium dark-adapted values (area L_o_(blue) / area L_o_(dark) = 1.10 ± 0.36, Fig. S5).

Our results confirm that the bulkiness of the neutral headgroup does not play a role in the membrane remodeling ability of our photoswitchable phytosphingolipids. Moreover, the increased backbone polarity of the phytosphingosine backbone does not seem to affect the way azo-phytosphingolipids engage in interactions with their neighboring lipids when compared to azo-sphingolipids. Hereby, we conclude that all the tested Azo-Cer, Azo-β-GalCer, Azo-SM, Azo-α-Gal-PhCer and Azo-PhCer establish stable interactions with other sphingolipids (such as SM) and sterols (such as Chol) inside L_o_ domains when their azobenzene acyl chain is in the *trans*-isoform (imitating a “straight” saturated acyl chain), and with unsaturated phosphatidylcholines (e.g. DOPC) inside L_d_ regions when the azobenzene is in the *cis*-isoform (mimicking a “bent” unsaturated chain).

### 2.5. Remodeling of membrane domains by 3-OH-blocked azo-sphingolipids

In order to infer the exact role of H-bonding and sphingoid base polarity for the mode of action of azo-sphingolipids, we used high-speed AFM to further investigate the photoswitching and lateral membrane remodeling activities of the azo-sphingolipids Azo-THP-Cer and Azo-THP-SM. These lipids have the 3-OH group on their sphingosine backbone protected with a bulky THP moiety. Noteably, the final protecting step resulted in an inseperable mixture of diastereomers at the THP linkage (see Fig. 3), which was used in all experiments.

**Fig. 3.**
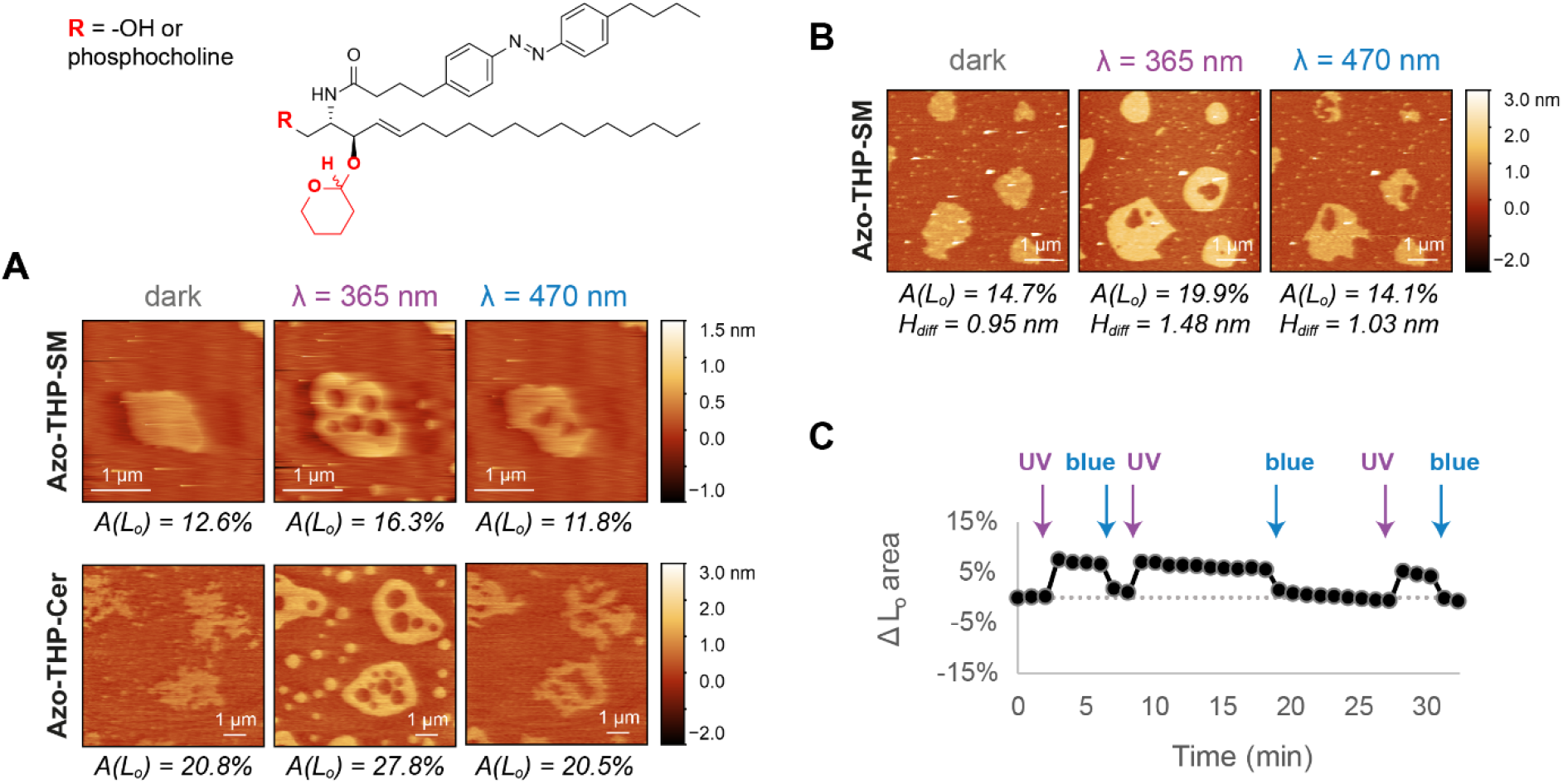
Lateral remodeling of phase-separated membranes containing 3-OH-blocked azo-sphingolipids upon light trigger analyzed by high-speed AFM. **(A)** Changes in the area of L_o_ domains before/after illumination with UV-A (λ = 365 nm) and blue (λ = 470 nm) lights on DOPC:Chol:SM:photolipid (10:6.7:5:5 mol ratio) SLBs having Azo-THP-SM or Azo-THP-Cer with a 3-OH-blocked (THP-protected) sphingosine backbone and varying headgroup functionality (R = phosphocholine or -OH, respectively). **(C-D)** Reversible lateral remodeling of a phase-separated SLB containing Azo-THP-SM, upon UV-A/blue light irradiation, as seen in Movie S9: (C) AFM images of the SLB at the dark-, UV- and blue light-adapted states, displaying the area occupied by the L_o_ phase and the L_d_-L_o_ height mismatches. (D) Relative variation of total L_o_ area of the SLB over time, shown in Movie S9, upon shining short pulses (marked with arrows) of UV-A and blue light.

In the dark-adapted *trans*-form, DOPC:Chol:SM:*photolipid* bilayers containing 3-OH-blocked Azo-THP-SM or Azo-THP-Cer (Figs. 3, S4D and Movies S7-S9) had L_o_ domains with irregular borders and lower height (~ 0.6 – 0.9 nm) when compared to SLBs with non-blocked counterparts (Fig. 2). Such a noticeable effect on the global architecture of L_o_ domains is evidence that 3-OH-blocked lipids are able to reduce the molecular packing within the L_o_ phase, as previously reported for 3-OH-blocked stearoyl-SM^[26,53]^.

When the membranes with THP-protected photoswitchable lipids were irradiated with UV-A light (λ = 365 nm), and the lipids converted to the *cis*-isoform, the total area of L_o_ phase markedly increased ~ 23% (area L_o_(UV) / area L_o_(dark) = 1.23 ± 0.21, Fig. S5), with L_o_ domains getting larger, rounder and noticeably heigher (~ 1.1 – 1.4 nm) (Fig. 3A–B). Quite strikingly, directly after exposure to UV-A light, L_d_ “lakes” inside pre-existing L_o_ domains, as well L_o_ “islands” within the L_d_ matrix were transiently formed. The pre-existing L_o_ domains then grew in total area, mainly via Ostwald ripening as L_o_ “islands” disappeared, whereas only few larger L_d_ “lakes” appeared at the end trapped inside the enlarged L_o_ domains.

Subsequent illumination with blue light (λ = 470 nm) led to an overall decrease of both, total L_d_-L_o_ height mismatch and L_o_ area back to the initial dark-adapted state values (area L_o_(blue) / area L_o_(dark) = 0.98 ± 0.20, Fig. S5), as the 3-OH-blocked photoswitchable lipids isomerized back to the *trans*-isoform. Interestingly, no noticeable formation of large L_d_ “lakes” or L_o_ “islands” was observed here. The L_o_ domains rapidly shrank with their domain borders becoming irregular (less rounded) and more unstable, as large rapid fluctuations were visible (Movies S7-S9). This clearly indicates that the THP-protected azo-sphingolipids, when in the *trans*-isoform, severely affect line tension of the phase-separated domains. Finally, the reported L_o_ domain height and area changes within the phase-separated bilayers could be repeated over multiple illumination cycles, as seen in Fig. 3C and Movies S7-S9.

To sum up, photoswitchable sphingolipids having their 3-OH sphingoid moiety blocked with a THP group promote a clearly distinct light-induced reorganization of L_d_-L_o_ phase-separated membranes, when compared to non-blocked counterparts. These blocked lipids are able to significantly increase the percentual amount and height of the L_o_ phase upon UV-triggered isomerization to the *cis*-isoform, and decrease both these parameters upon blue light-triggered isomerization to the *trans*-isoform.

### 2.6. Phase-separation area changes by photoswitchable sphingolipids

After demonstrating that the various non-blocked vs. hydroxyl-blocked photoswitchable sphingolipids reorganize phase-separated membranes differently, we set out to quantitatively compare the extent by which these lipids alter the total distribution of phase-separation, as well as other structural membrane parameters.

To begin, since AFM only allows us to follow a limited number of L_o_ domains simultaneously, we acquired additional large field-of-view fluorescence confocal images (Figs. S7, S8A and 4A), to obtain better statistics for determining ensemble area values; independent of domain size and number. We analyzed changes in L_o_ total area before/after UV/blue irradiation on DOPC:Chol:SM:*photolipid* SLBs containing 18.7 mol% Azo-Cer, Azo-β-GalCer, Azo-SM, Azo-PhCer, Azo-α-GalPhCer, Azo-THP-Cer or Azo-THP-SM, doped with 0.1 mol% Atto655-DOPE for fluorescent detection of the L_d_ phase. Usage of fluorescence allowed us to easily generate binary masks (Fig. S7), from which L_o_ phase areas could be straightforwardly estimated.

**Fig. 4.**
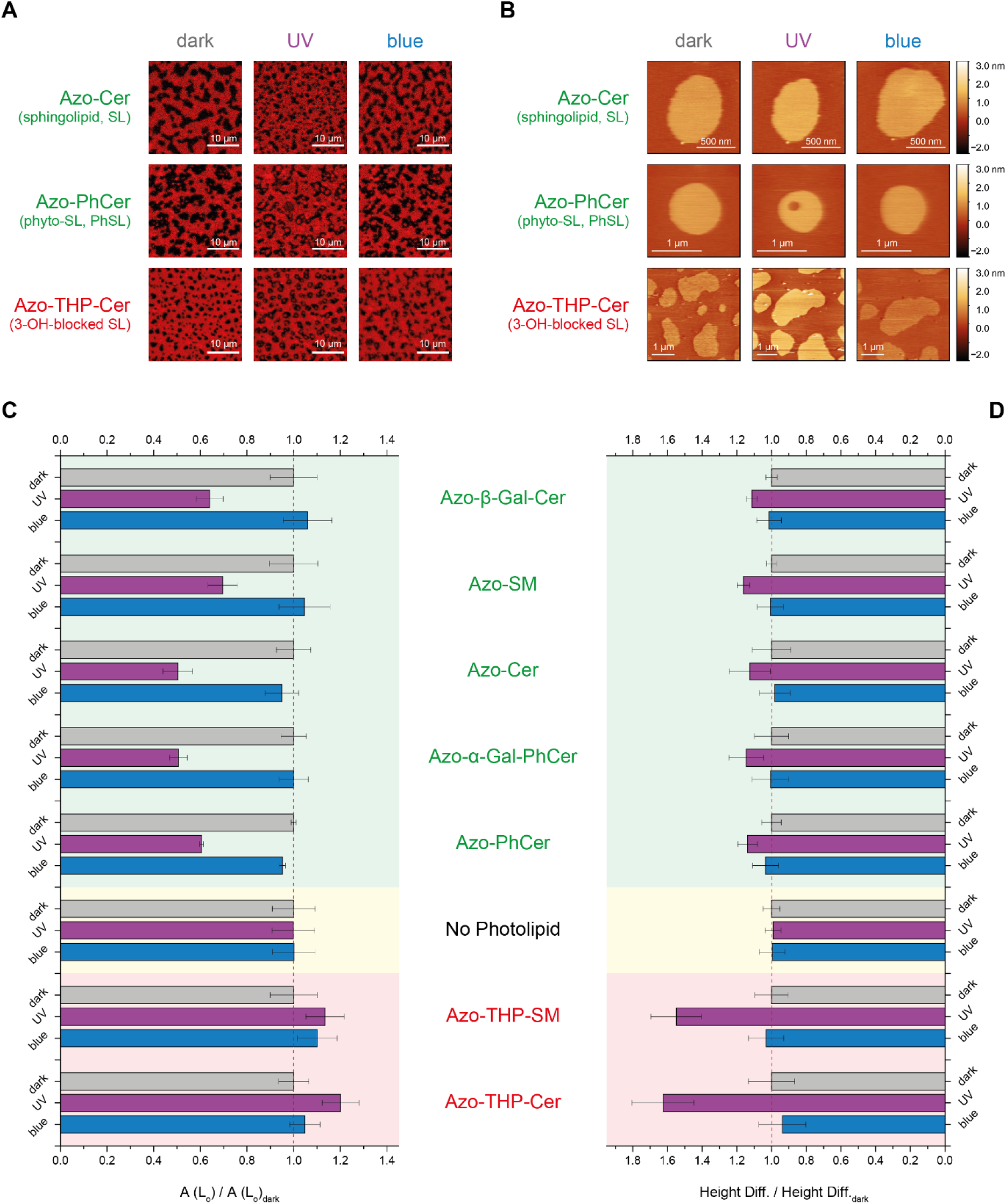
Normalized changes in the total L_o_ phase area and L_d_-L_o_ height difference on phase-separated SLBs having different types of azo-sphingolipids, upon application of UV-A (λ = 365 nm) and blue (λ = 470 nm) lights. Fluorescence confocal (A) and AFM (B) images of DOPC:Chol:SM:photolipid (10:6.7:5:5 mol ratio) SLBs with sphingosine-based Azo-Cer, phyto-sphingosine-based Azo-PhCer or 3-OH-blocked sphingosine-based Azo-THP-Cer, all having the same-OH headgroup but distinct sphingoid backbone. Normalized L_o_ areas (C) and L_d_-L_o_ height differences (D), respectively recovered from fluorescence confocal and AFM data, for phase-separated SLBs having either azo-(phyto)sphingolipids with free 3-OH (marked in green); no azo-sphingolipid (controls with SM, marked in yellow); or THP-protected azo-sphingolipids with the 3-OH blocked (marked in red). Error bars correspond to standard error of the mean (n = 5-9).

To simplify data comparison, the L_o_ areas were then normalized before/after UV/blue light illumination by the average L_o_ area for the various individual SLBs at the dark-adapted state (Fig. 4C). In addition, all non-normalized average L_o_ area values recovered for the various SLBs are represented in Fig. S8B.

As seen in Figs. 4C and S8B, the total L_o_ area for phase-separated SLBs having sphingosine- or phytosphingosine-based photoswitchable lipids decreased in average by 41% (area L_o_(UV) / area L_o_(dark) = 0.59 ± 0.04) after UV-A illumination and augmented back to the original dark-adapted state (area L_o_(blue) / area L_o_(dark) = 1.00 ± 0.02) upon blue light irradiation. For SLBs having Azo-THP-SM or Azo-THP-Cer, on the contrary, the total area of the L_o_ phase increase by 17% (area L_o_(UV) / area L_o_(dark) = 1.17 ± 0.03) after illumination with UV-A light, while the recorded amount of L_o_ phase decreased back to the original dark-adapted state value (area L_o_(blue) / area L_o_(dark) = 1.07 ± 0.03) after the application of blue light pulse.

Interestingly, if we compare the changes in L_o_ area after azo-sphingolipid photo-isomerization determined by fluorescence (Figs. 4C and S8) vs. high-speed AFM data (Fig. S5), the area changes for confocal microscopy seem to be skewed towards detecting higher amounts of L_d_ phase after UV irradiation. This skew may be a direct consequence of the limited pixel resolution of conventional laser-scanning confocal microscopy for detecting nanoscale L_o_ domains, when compared to AFM. Despite this instrumental bias, similar trends in membrane domain area variations were detected with both, fluorescence confocal and AFM techniques.

These experiments corroborate that sphingosine- and phytosphingosine-based photoswitchable lipids rely on the same principles for reshuffling membrane phase-separated domains; whereas the THP-protected counterparts, owing their distinct physicochemical properties, follow a markedly different mechanism.

### 2.7. Domain height mismatch changes by photoswitchable sphingolipids

Our results so far clearly point out that blocking the interfacial hydroxyl on the sphingoid backbone has a marked effect on the molecular organization of individual lipids and on the global architecture of L_o_ domains. Thus, to quantitatively ascertain how photoswitchable sphingolipids affect the structure and physicochemical properties of L_o_ domains within phase-separated membranes, we collected zoomed-in and high-resolution low-speed AFM images of individual L_o_ domains on DOPC:Chol:SM:*photolipid* SLBs, prior and after illumination with UV-A/blue light, as depicted in Figs. 4B, S9 and S10A. This acquisition mode allows us to follow the membrane contour with an increased signal-to-noise ratio and therefore determine more accurately the height differences between the L_o_ domains and the surrounding L_d_ matrix (Fig. S9). This is an important parameter, as it relates to the hydrophobic mismatch between the saturated (“non-bent” acyl chains) lipids in raft-mimicking L_o_ domains and the unsaturated (“bent” acyl chains) lipids in the more fluid and less packed L_d_ regions.

Altogether, the average height difference between L_o_ and L_d_ regions (at the dark-adapted state for SLBs having either sphingosine- or phytosphingosine-based photolipids was 1.26 ± 0.11 nm. This value corresponds to the mean of all average L_d_-L_o_ height differences (± standard error) obtained for membranes containing Azo-Cer, Azo-β-GalCer, Azo-SM, Azo-PhCer, Azo-α-GalPhCer (Fig. S10B); and was very close to the less precise values previously reported using high-speed AFM. Owing to the exquisite *z*-resolution of AFM, we also identified that the L_o_ domains of SLBs having azo-sphingolipids with smaller headgroups (e.g. Azo-Cer and Azo-PhCer) were slightly less elevated (1.04 ± 0.09 nm) than the L_o_ domains of SLBs having azo-sphingolipids with larger headgroups (e.g. Azo-SM, Azo-β-GalCer and Azo-α-GalPhCer). The later displayed L_d_-L_o_ height mismatches (1.41 ± 0.09 nm) closer to the values recovered (1.76 ± 0.05 nm) for control ternary mixtures without photoswitchable lipid (. Hence, our observations corroborate a preferred localization of the *trans*-azo-sphingolipids inside L_o_ domains, as these lipids could then engage hydrophobic packing and stable H-bonding interactions with SM and Chol, altering slightly the height of L_o_ domains due to the different headgroup size and *N*-acyl chain length (e.g. C18:0 acyl chain: 21.2 Å vs. FAAzo-4: 17.9 Å, retrieved from Chem3D, PerkinElmer).

Interestingly, upon applying UV-A light to SLBs having these non-blocked photoswitchable lipids, the L_d_-L_o_ height mismatch increased in average by 14% (1.44 ± 0.12 nm; Fig. 4D). This is in line with the exclusion of *cis*-azo-sphingolipids from the L_o_ phase and SM being then the predominant sphingolipid molecules inside those domains. Subsequent irradiation with blue light led to the decrease of the domain height by 13%, back to the original values reported for the dark-adapted state (Fig. 4D); corroborating a re-partitioning of *trans*-azo-sphingolipids back the L_o_ phase.

For membranes having hydroxyl-blocked photoswitchable lipids, the height of L_o_ domains at the dark-adapted state was lower and the domain boundaries were more irregular, when compared to membranes with non-blocked counterparts. Indeed, the L_d_-L_o_ height mismatch observed for SLBs with Azo-THP-SM or Azo-THP-Cer was below 1 nm (0.73 ± 0.05 nm; Fig. S10B); similar to the values observed using high-speed AFM, and nearly 0.5 nm and 1.0 nm lower than the height mismatch found for SLBs with Azo-SM or control membranes lacking azo-sphingolipids, respectively. The lower L_o_ height observed for membranes with 3-OTHP-lipids indeed corroborates a preferential localization of these lipids in the L_o_ phase when in the *trans-*isoform, but most importantly validates that blocking H-bonding severely alters inter-lipid interactions, molecular packing, as well line tension within L_o_ domains.

This destabilization effect can be overcome once the hydroxyl-blocked azo-sphingolipid is converted to its *cis*-isoform upon illumination with UV-A light. Indeed, after applying UV-A to the phase-separated SLBs with Azo-THP-SM, the L_d_-L_o_ height mismatch increased by 55% (Fig. 4D) to average values above 1 nm (1.13 ± 0.07 nm). This elevation in height suggests that Azo-THP-SM and Azo-THP-Cer lipids are expelled from the L_o_ phase when in the *cis-* isoform, leaving the L_o_ domains mainly composed by SM and Chol. Without the interference of these THP-protected lipids, SM and Chol molecules can then establish more stable H-bonding and tighter hydrophobic chain packing interactions, giving rise to taller, rounder and larger L_o_ domains. In opposition, irradiation of the phase-separated SLBs with blue light leads to a marked reduction of the L_o_ domain height (Fig. 4D), back to the initial dark-adapted state value (0.75 ± 0.05 nm). As the 3-OH-blocked azo-sphingolipids isomerize back to their *trans*-isoform, these lipids could then re-establish hydrophobic chain packing interactions with the other raft-localizing, destabilizing the existing H-bonding interactions between SM and Chol.

### 2.8. Domain line tension changes by photoswitchable sphingolipids

A parameter closely linked to the domain height mismatch is line tension, which can be perceived as the interfacial energy arising at the boundaries of coexisting phases and is an important driving force for membrane shape transformation (e.g. budding^[4,54,55]^ and fusion^[56]^). In order to estimate the approximate values of line tension for the various raft-mimicking membranes with distinct blocked and non-blocked photoswitchable sphingolipids, based on the height mismatch values measured using low-speed AFM, we used the theoretical model implemented by Cohen and coworkers (Eq. 1). This model describes a quadratic dependence of the line tension with the phase height mismatch^[57]^, and was previously used to estimate line tension on phase-separated membranes with similar lipid composition^[58]^.

Overall, as seen in Fig. S10C, line tension values ranged from 1.9 – 4.3 pN for L_d_-L_o_ phase-separated SLBs with sphingosine- and phytosphingosine-based lipids in the *trans*-isoform. When the lipids are in the *cis*-isoform and partition to the L_d_ phase instead, a small increase in line tension by 24% (Fig. S11) was recovered, being the values very close to the ones gauged for phase-separated SLBs lacking azo-sphingolipids (5.4 ± 0.6 pN).

In contrast, raft-like bilayers with THP-protected azo-sphingolipids (such as Azo-THP-SM and Azo-THP-Cer) in the *trans*-isoform possess noticeably reduced line tension values (~ 1.2 pN); 2.2 – 4.5-fold lower than the domain line tension measured for SLBs with non-blocked photolipid counterparts (Fig. S10C). Next, when the hydroxyl-blocked lipids are in their *cis*-isoform and locate in the L_d_ phase, the domain line tension greatly increases by ~ 120% (Fig. S11), reaching values close to the ones reported for non-blocked counterparts (2.6 pN). Hence, THP-protected photoswitchable lipids, when in *trans*, greatly reduce the line tension of L_o_ domains in opposition to the azo-sphingolipids with free interfacial hydroxyls, appearing to possess additional line-active (lineactant) properties.

Line-active molecules are known to concentrate at the boundaries of membrane phases^[59–62]^, reducing the hydrophobic mismatch and line tension around phase-separated domains (e.g. L_o_ vs. L_d_). Herein, hybrid lipids such as palmitoyl-oleyl-phosphatidylcholine (POPC), possessing both a saturated and unsaturated fatty acid chain, are of particular relevance. When added to ternary mixtures made of lipids with two saturated tails (e.g. DPPC or DSPC), two unsaturated tails (DOPC) and a sterol (Chol), POPC was shown to localize around the borders of the L_o_ phase promoting the reduction of line tension and formation of nanoscopic domains^[63–67]^. Interestingly, Azo-THP-SM and Azo-THP-Cer also appear to reduce line tension and promote the formation of nanoscopic domains in a similar way, sharing moreover key structural similarities with hybrid lipids: a) the *trans*-azobenzene *N*-acyl chains mimic saturated fatty acids prone to localize within L_o_ domains, b) the bulky THP moiety at the sphingosine base appears to interfere with the molecular packing of lipids and be therefore susceptible to preferentially localize in the less packed L_d_ phase.

To sum up, the hybrid chain properties in addition to the blockage of H-bonding could be possible explanations for the observed perturbation of the L_o_ domain boundaries and reduction of domain height, by THP-protected azo-sphingolipids in the *trans*-isoform.

### 2.9. Changes in the mechanics (indentation forces) of homogeneous membranes promoted by the photoisomerization of azo-spingolipids

Despite the differences in membrane domain remodeling by THP-protected vs. non-protected azo-sphingolipids, we also observed the generation of occasional membrane holes when isomerizing 3-OH-blocked photoswitchable lipids from the bent *cis*-isoform to the straight *trans*-isoform, as depicted in Fig. S12 for phase-separated membranes with Azo-THP-SM. This hole formation clearly indicates that SLBs containing THP-protected azo-sphingolipids, similarly to membranes with non-blocked counterparts (Fig. S6A), globally expand after UV-A illumination and subsequently compress upon illumination with blue light, due to the bending and unbending on the *N*-acyl chains respectively. Hence, photoswitching of azo-sphingolipids not only influences lipid phase-separation (as discussed so far), but irrefutably affects basic mechanical properties of the membrane, such as packing and stiffness/fluidity.

In order to evaluate whether photoswitchable sphingolipids indeed interfere with global membrane mechanics, irrespectively of phase-separation, we performed additional AFM-based force spectroscopy measurements^[68–75]^ on non-phase-separated SLBs composed of DOPC:Chol:*photolipid* (10:6.7:10 molar ratio). More precisely, we evaluated homogenous bilayers containing either Azo-Cer as non-blocked azo-sphingolipid (Fig. 5A), or Azo-THP-SM as 3-OH-blocked azo-sphingolipids (Fig. 5D). In short, when indenting the membranes with an AFM tip, a typical jump (or discontinuity) corresponding to the force required to pierce (or break through) the supported lipid bilayer can be easily identified within the collected force-displacement curves (as seen in Fig. 5B,E). The extent of such breakthrough forces is then directly linked to the mechanical properties of the membrane: lower forces are expected when the membranes are more fluid (or less compact); higher forces when these are stiffer (or more compact). Thus, upon recording a set of force curves prior and after illumination with UV-A (λ = 365 nm) and blue (λ = 470 nm) lights, we evaluated the breakthrough events and displayed the recovered forces needed to pierce the membranes (Fig. 5C,F) as histograms (non-normalized values depicted in Fig. S13).

**Fig. 5.**
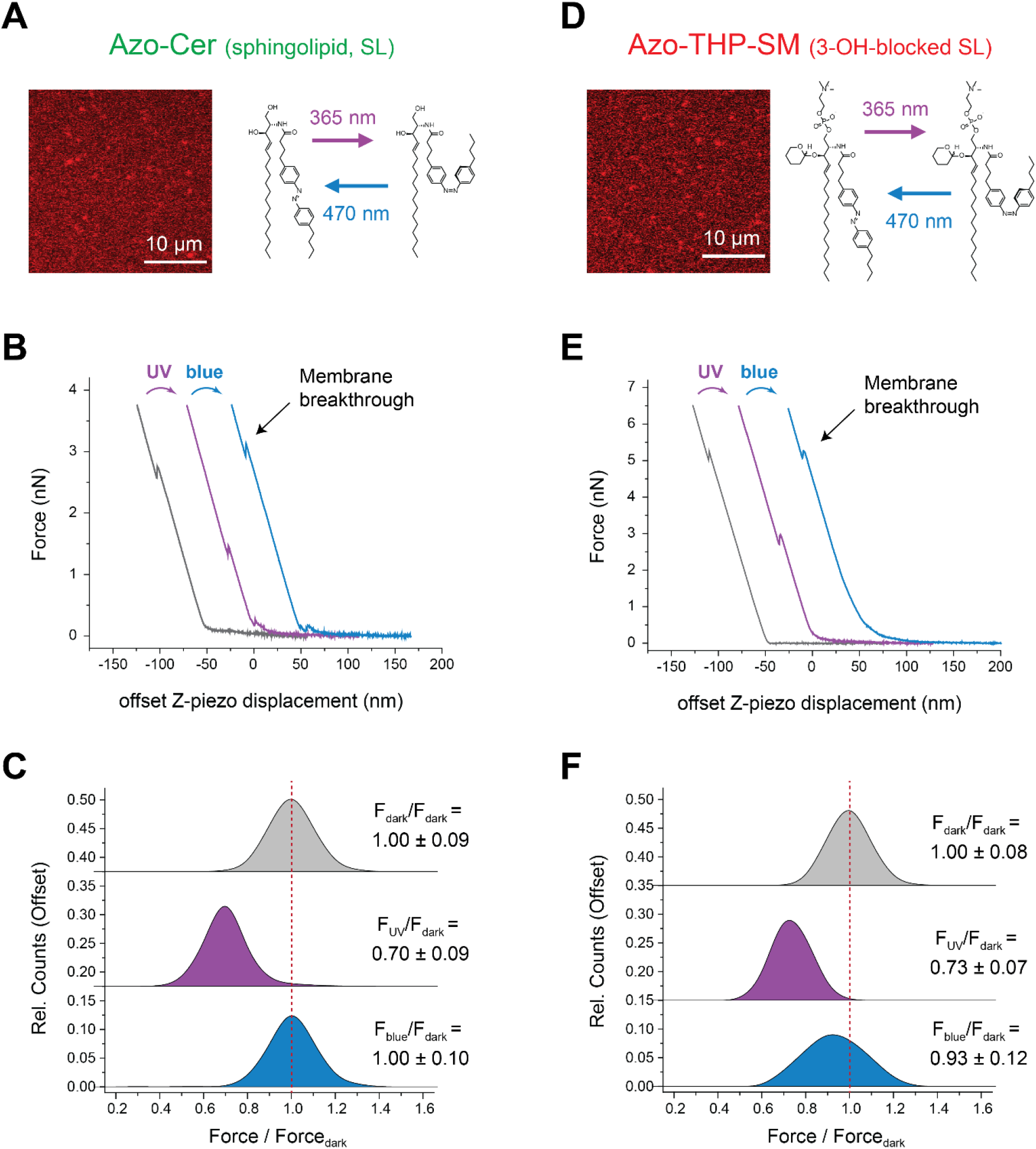
Breakthrough forces of homogeneous membranes containing non-blocked Azo-Cer (A-C) and 3-OH-blocked Azo-THP-SM (D-F) azo-sphingolipids. (A, D) Confocal images of DOPC:Chol:photolipid (10:6.7:10 mol ratio) supported membranes doped with 0.1 mol% Atto655-DOPE for fluorescence detection. (B, E) Force spectroscopy indentation curves (Z-piezo displacement) of homogeneous SLBs containing azo-sphingolipids upon illumination with UV-A (λ = 365 nm) and blue (λ = 470 nm) lights. Characteristic membrane breakthrough events for the AFM tip pinching through the SLB marked with arrows. (C, F) Membrane breakthrough histograms normalized to the dark-adapted state for SLBs containing Azo-Cer or Azo-THP-SM prior and after irradiation with UV-A and blue lights.

For homogenous DOPC:Chol:Azo-Cer SLBs, a breakthrough force of 2.93 ± 0.26 nN (Fig. S13A) was recorded in the dark-adapted state. After illumination with UV light, and consequent conversion of Azo-Cer to its bent *cis*-isoform, the force required for piercing the membrane reduced by ~ 30% (Fig. 5C), to 2.06 ± 0.27 nN (Fig. S13B). Irradiation with blue light, on the contrary, promoted an increase of the breakthrough force, back to its original average value (2.93 ± 0.28 nN; Fig. S13C), as Azo-Cer would back-isomerize to its *trans*-isoform. Thus, for non-blocked azo-sphingolipids, we confirmed that the *cis*-isoform expands/fluidifies the membrane, while *trans*-isoform compacts/stiffens it. In this context, our force data clearly backs up a previous study^[41]^ based on fluorescence recovery after photobleaching (FRAP), which showed that lateral lipid diffusion on a membrane made of Azo-PC (phosphatidylcholine analog with a FAAzo-4 acyl chain) was higher when the photoswitchable lipid was in its *cis*-isoform and slower when in the *trans*-isoform.

Likewise, for non-phase-separated DOPC:Chol:Azo-THP-SM bilayers, similar trends were recorded. More precisely, the piercing force needed to break through those membranes also reduced by ~ 30% (Fig. 5F), from 4.85 ± 0.39 nN (Fig. S13D) to 3.56 ± 0.34 nN (Fig. S13E), upon irradiation with UV light and formation of *cis*-Azo-THP-SM; in agreement with a global expansion or fluidification of the membrane. Subsequently, the membrane breakthrough force also reverted back close to the original value (4.51 ± 0.60 nN; Fig. S13F) upon illumination with blue light and formation of *trans*-Azo-THP-SM: in agreement with a global compaction or rigidification of those membranes.

Therefore, based on this force spectroscopy outcome for homogenous membranes, we can argue that the opposite changes in L_o_ area for phase-separated SLBs containing either 3-OH-blocked and non-blocked azo-sphingolipids (observed throughout the previous manuscript sections) are mainly due to different types of interactions these photoswitchable lipids engage via their sphingoid backbone with neighboring lipids within L_o_ domains, and are not directly linked to the structural properties of the *N*-acyl photoswitch *per se*.

## 3. Conclusions

In this work, we evaluated physicochemical foundations for the membrane remodeling ability by a family of photoswitchable sphingolipids, deciphering the relative contributions of the lipid headgroup and sphingoid backbone. We synthesized new types of *N*-acyl azobenzene sphingolipids with varying headgroup and sphingoid base functionalities. Then, we studied with the help of atomic force and fluorescence microscopies the propensity of these photoswitchable lipids to alter membrane properties and laterally remodel L_d_-L_o_ phase-separated supported membranes. Overall, we demonstrated that the headgroup type (simple hydroxyl vs. more complex galactosyl or phosphocholine) does not interfere with the photoswitching ability of the various azo-sphingolipids within raft-mimicking lipid mixtures. Owing to the photo-dynamic reversibility of the azobenzene *N*-acyl chain, we further highlighted that *trans*-*photolipids* (*i.e.* dark-adapted and blue light-illuminated states) predominantly localize within pre-existing raft-like L_o_ domains and compact membranes, while *cis*-*photolipids* (*i.e.* UV-A-illuminated state) preferentially locate within the more fluid L_d_ membrane regions and expand membranes.

Importantly, our results provide clear evidence that the nature of the sphingoid backbone, and their ability for engaging stable H-bonding interactions with other co-lipids, play a fundamental role in the way photoswitchable sphingolipids remodel L_d_-L_o_ phase-separated membranes and change the amount, size and height of L_o_ domains. Sphingosine- and phytospingosine-based lipids, with their free interfacial 3-OH and 4-OH hydroxyls, do not significantly alter the height of L_o_ domains when in the *trans*-isoform. In contrast, THP-protected lipids, with their interfacial 3-OH blocked, greatly interfere with the molecular packing and line tension of L_o_ domains, markedly reducing the overall L_o_ height mismatch. Whereas non-blocked azo-sphingolipids will promote a decrease of the total L_o_ phase area upon UV trigger, THP-protected azo-sphingolipids will increase the total L_o_ area, as well as induce a marked rise in L_o_ domain height after illumination with UV light.

Taken together, the structural diversity of the photoswitchable sphingolipids presented here, as well as exquisite understanding of how these lipids alter important membrane properties, may offer new strategies for controlling the structure of biological lipid bilayers and the localization of membrane-interacting proteins. Thus, by further expanding the headgroup repertoire of photoswitchable lipids, we may soon be in the position to target the fate of biologically-relevant proteins on membranes using light as trigger. Such endeavor would not only open up new exciting avenues for optodynamic applications in the fields of synthetic biology, structural biology or biophysics, but also offer novel perspectives towards the development of innovative photo-responsive drugs and pharmacological therapies.

## 4. Experimental section

### 4.1. Synthesis of *N*-acyl azobenzene-modified sphingolipids

A protocol for the synthesis and analysis of all photolipids can be found in the supporting information (SI).

### 4.2. Membrane model systems

Throughout this work, small unilamellar vesicles (SUVs) and supported lipid bilayers (SLBs) were used as lipid membrane model systems. These were primarily composed by *N*-stearoyl-D-*erythro*-sphingosylphosphorylcholine (C18-SM, or simply SM), 1,2-dioleoyl-*sn*-glycero-3-phosphocholine (DOPC) and cholesterol (Chol), which were purchased from Avanti Polar Lipids (Alabaster, AL, USA); to which different photoswitchable lipids, notably Azo-Cer, Azo-PhCer, Azo-THP-Cer, Azo-α-Gal-PhCer, Azo-β-Gal-Cer, Azo-SM or Azo-THP-SM were mixed. Unless otherwise stated, the typical lipid composition was DOPC:Chol:SM:*photolipid* with a 10:6.7:5:5 mol ratio. For fluorescence detected, lipid mixtures were also doped with 0.1 mol% Atto655-DOPE, purchased from ATTO Technology GmbH (Siegen, Germany).

SUVs were obtained through bath sonication of multilamellar vesicles. Briefly, the desired lipid mixtures dissolved in choloform:methanol (7:3) were added to a glass vial and the solvent was then evaporated using N_2_ flow, followed by vacuum-drying in a desiccator. Lipids were rehydrated by adding HEPES buffer (10 mM HEPES, 150 mM NaCl, pH 7.4), reaching a final lipid concentration of 10 mM, and then vigorously vortexed forming a suspension of multilamellar vesicles. These were then diluted to 1mM with HEPES buffer, and sonicated in an ultrasonic bath for 10-20 min until the suspension became clear, giving rise to SUVs.

SLBs were prepared by deposition and fusion of SUVs on top of freshly glued-mica glued on a borosilicate coverglass, as described elsewhere^[36]^. Shortly, SUV suspensions (at 1 mM lipid concentration) were deposited in the presence of 2 mM CaCl2 on freshly-cleaved mica. The samples were then incubated for 20 min at 65 °C, rinsed with HEPES buffer and allowed to slowly cool down to room temperature for at least 1 h.

### 4.3. UV-Vis spectra of membrane-embedded azo-sphingolipids

UV-Vis spectra of the various azo-sphingolipids embedded within SUVs were collected with Hellma SUPRASIL precision quartz cuvettes (10 mm light path) on a Jasco V-650 spectrophotometer (Tokyo, Japan), before and after illumination with UV-A or blue lights. More precisely, SUV suspensions at 150 μM lipid concentration, composed of DOPC:Chol:SM:*photolipid* (10:6.7:7:3 mol ratio) were here utilized.

### 4.4. Laser scanning confocal fluorescence microscopy

Fluorescence microscopy was performed on a Zeiss LSM 510Meta laser scanning microscope (Jena, Germany) using a water immersion objective (C-Apochromat, 40× 1.2W UV-VIS-IR). Samples were excited with the 633 nm line of a He-Ne laser for Atto655 excitation. Images were typically recorded with 1 Airy unit pinhole and 512×512 pixel resolution. Image analysis was performed using Fiji software (http://fiji.sc/Fiji).

#### Segmentation methods

In order to quantify the lipid domain (liquid-ordered phase), the confocal data was processed by a custom-made MATLAB script for batch processing. The algorithm performs basic segmentation operations based on thresholding (Otsu’s Method), morphological erosion and dilation operations. The output is an image in Portable Network Graphic format of the positive mask of the dark regions corresponding to the lipid domains and a text file containing the calculated area ratio of the domains for each image.

### 4.5. Atomic force microscopy and force spectroscopy

Atomic force microscopy (AFM) was performed on a JPK Instruments Nanowizard Ultra (Berlin, Germany) mounted on the Zeiss LSM510 Meta laser scanning confocal microscope (Jena, Germany). High-speed and normal-speed AFM, both in AC mode, were done with USC-F0.3-k0.3 ultra-short cantilevers from Nanoworld (Neuchâtel, Switzerland) with typical stiffness of 0.3 N/m. The cantilever oscillation was tuned to a frequency of 100-150 kHz and the amplitude was kept below 10 nm. Scan rate was set to 25-150 Hz for high-speed AFM and to 2-6 Hz for normal-speed AFM. For both modes, images were acquired with a typical 256×256-pixel resolution. All measurements were performed at room temperature. The force applied on the sample was minimized by continuously adjusting the set point and gain during imaging. Height, error, deflection and phase-shift signals were recorded and images were line-fitted as required. Data was analyzed using JPK data processing software Version 6.0.55 (JPK Instruments) and Gwyddion Version 2.49 (Czech Metrology Institute).

Line tension (γ) was determined as previously reported^[58]^ using the equation by Cohen and coworkers^[57]^:

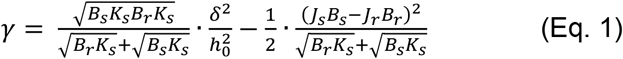

being *δ* the L_d_-L_o_ height mismatch, h the monolayer thickness with *h*_o_ = (*h*_r_ + *h*_s_)/2, *B* the elastic splay modulus, *K* the tilt modulus, and *J* the spontaneous curvature of the monolayer. Herein, the subscripts *r* and *s* refer to the L_o_ (*rigid*) and L_d_ (*soft*) membrane phases, respectively. For calculating the effective heights we used the height mismatches obtained for the various samples and considered a thickness of the L_d_ bilayer of 3.9 nm, as measured in Fig. S6. Finally, as described in García-Sáez et al. ^[58]^ we assumed *B_r_ = B_s_* = 10 k_B_T, *K_r_ = K_s_* = 40 mN/m, and *J_r_ = J_s_* = 0.

Force spectroscopy measurements were performed using uncoated silicon cantilevers CSC38 from MikroMasch (Tallinn, Estonia), with a spring constant of 0.12 N/m, as previously described^[73,74]^. Shortly, sensitivity and spring constant calibration were done via the thermal noise method. The total z-piezo displacement was then set to 300 nm, indenting approach speed to 800 nm/s, and the retraction speed was 200 nm/s, and maximal setpoint to 5-7 nN. Force measurements were carried out at different points of the lipid bilayers. Identification of the breakthrough events on an average of 200 approach force curves was done using the JPK data processing software Version 6.0.55 (JPK Instruments), whereas the retrieved yield forces were plotted in histograms using OriginPro2015 (OriginLab).

### 4.6. Compound switching on SUVs and SLBs

Photoswitching of the photolipid compounds was achieved using a CoolLED pE-2 LED light source (Andover, United Kingdom) for illumination at λ = 365 nm and 470 nm. The light source was typically operated for ~ 20 s at 80% power. For the UV-Vis spectroscopic experiments with SUVs inside cuvettes, the light beam was guided by a fiber-optic cable directly to the cuvette top. For microscopic experiments, the light beam was guided by an optical fiber directly through the objective of the LSM510 Meta microscope via a collimator at the backport.

## Acknowledgements

The project was funded by the Deutsche Forschungsgemeinschaft (DFG, German Research Foundation) – SFB-1032 – Project ID 201269156. Additional support was provided by the Center for NanoScience (CeNS). H.G.F and P.S acknowledge the financial support by the DFG within the SFB 863 (project ID 111166240). D.T acknowledges the European Research Council (ERC Advanced Grant #268795, “CARV”) and the Deutsche Forschungsgemeinschaft (SFB 749 and CIPSM) for generous funding. N.H. acknowledges financial support by the Deutsche Telekom Foundation and the LMUMentoring program. Further support was given by the Max Planck Society to P.S. The authors thank Sigrid Bauer for assistance in lipid handling and Alena Khmelinskaia for helpful discussions.

## Contributions

H.G.F and S.M.L conducted the biophysical experiments and analyzed data. N.H. synthesized and purified most photoswitchable lipids, in cooperation with N.W, H.T-R and J.A.F. P.S and D.T provided funding. D.T supervised the synthesis of photoswitchable lipids and H.G.F their biophysical characterization. H.G.F, N.H and S.M.L prepared the first manuscript draft. All authors contributed to revision of the manuscript.

## Conflicts of interest

There are no conflicts of interest.

## Notes

### Competing Interest Statement

The authors have declared no competing interest.

