## Supporting Information for "Structural Diversity of Photoswitchable Sphingolipids for Optodynamic Control of Lipid Raft Microdomains"

#### Supplementary figures

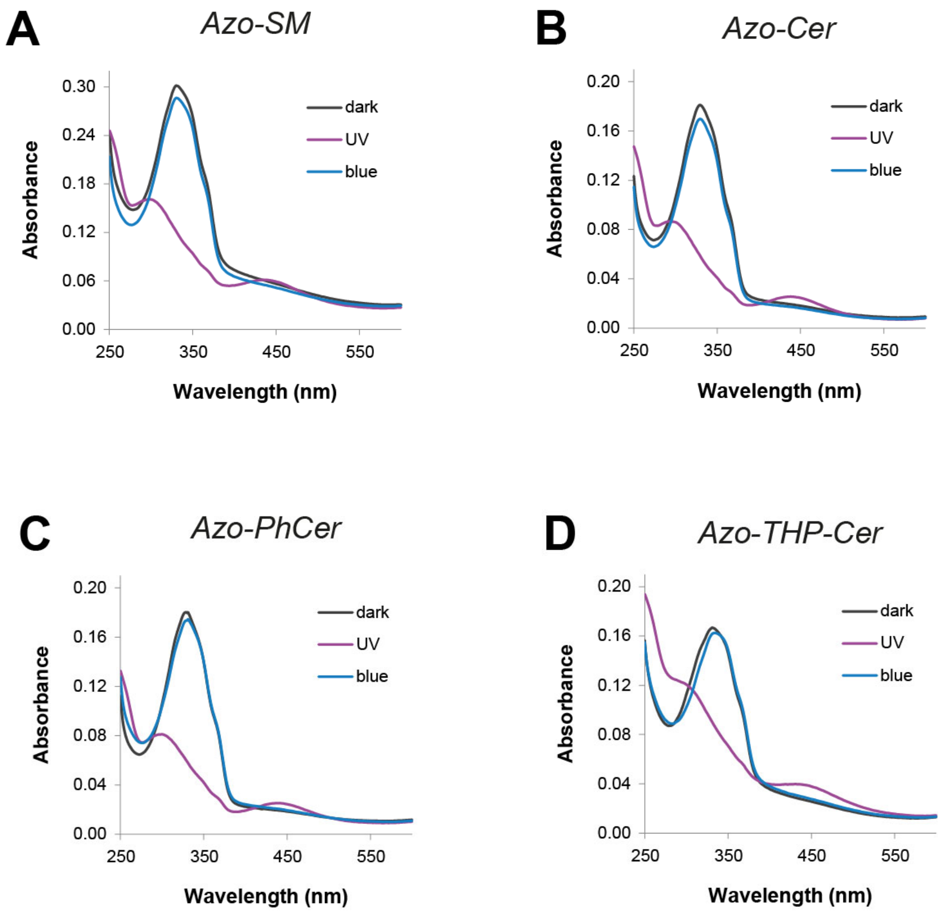

**Fig. S1 – UV-Vis absorbance spectra of azo-sphingolipids incorporated in small unilamellar vesicles (SUVs).** (A) Azo-SM, (B) Azo-Cer, (C) Azo-PhCer and (D) Azo-THP-Cer. Black curves correspond to the spectra of the photolipids at their dark-adapted state (black curves), purple curves to the spectra after shining with UV-A light (λ = 365 nm) and blue curves after applying blue light (λ = 470 nm).

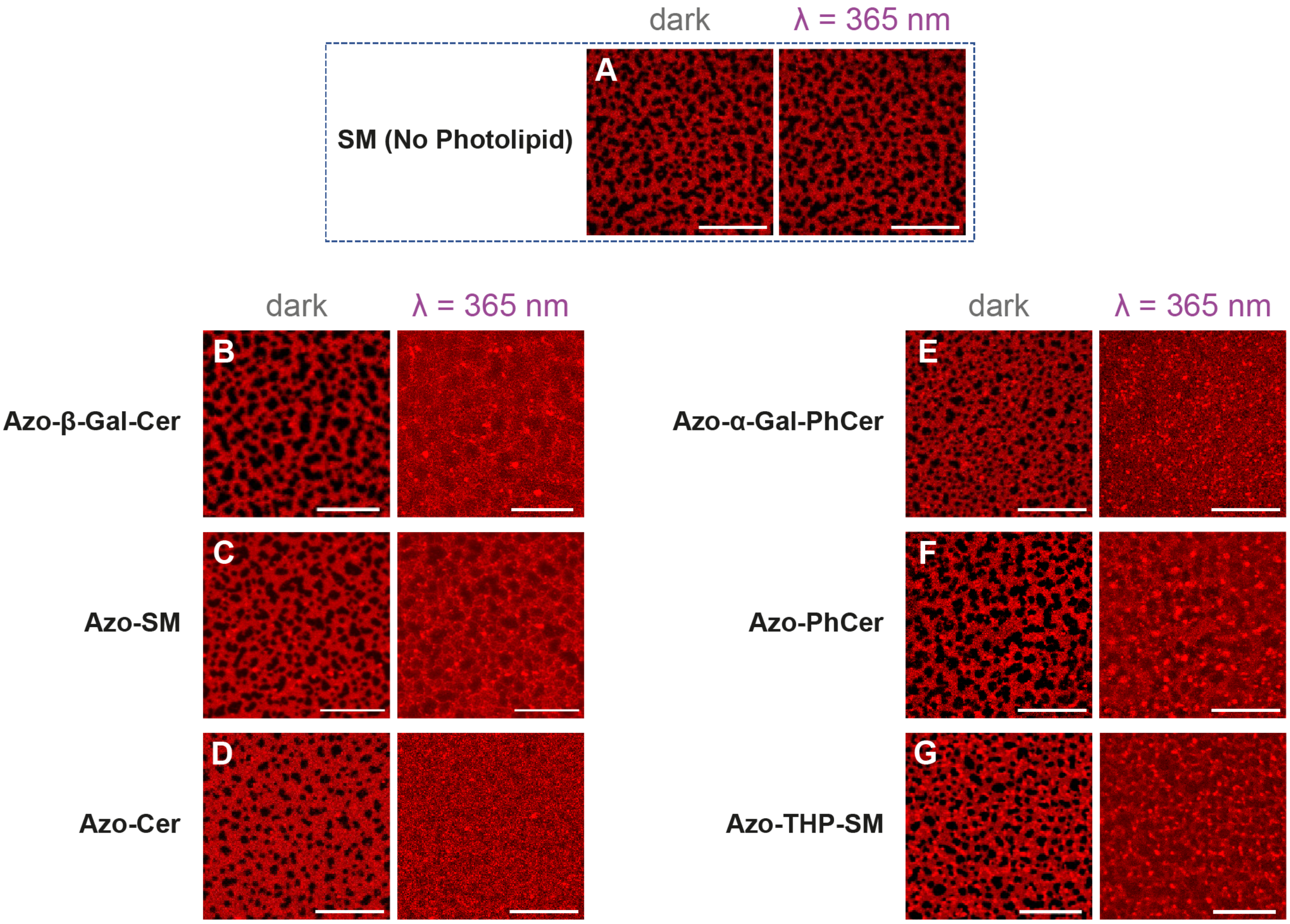

**Fig. S2 – Remodeling of phase-separated supported lipid bilayers (SLBs) containing azo-sphingolipids directly after irradiation with UV-A light.** Fluorescence confocal images showing the admixing of the $L_d$-$L_o$ lipid phases on SLBs composed of DOPC:Chol:SM:photolipid (10:6.7:5:5 mol ratio), doped with 0.1 mol% Atto655-DOPE (for fluorescence detection of $L_d$ phase), before and directly after illumination with UV-A light (λ = 365 nm). (A) Control sample without photolipid (DOPC:Chol:SM (10:6.7:10 mol ratio)). Samples with 18.7 mol% (B) Azo-β-Gal-Cer, (C) Azo-SM, (D) Azo-Cer, (E) Azo-α-Gal-PhCer, (F) Azo-PhCer and (G) Azo-THP-SM. Scale-bar is 20 µm.

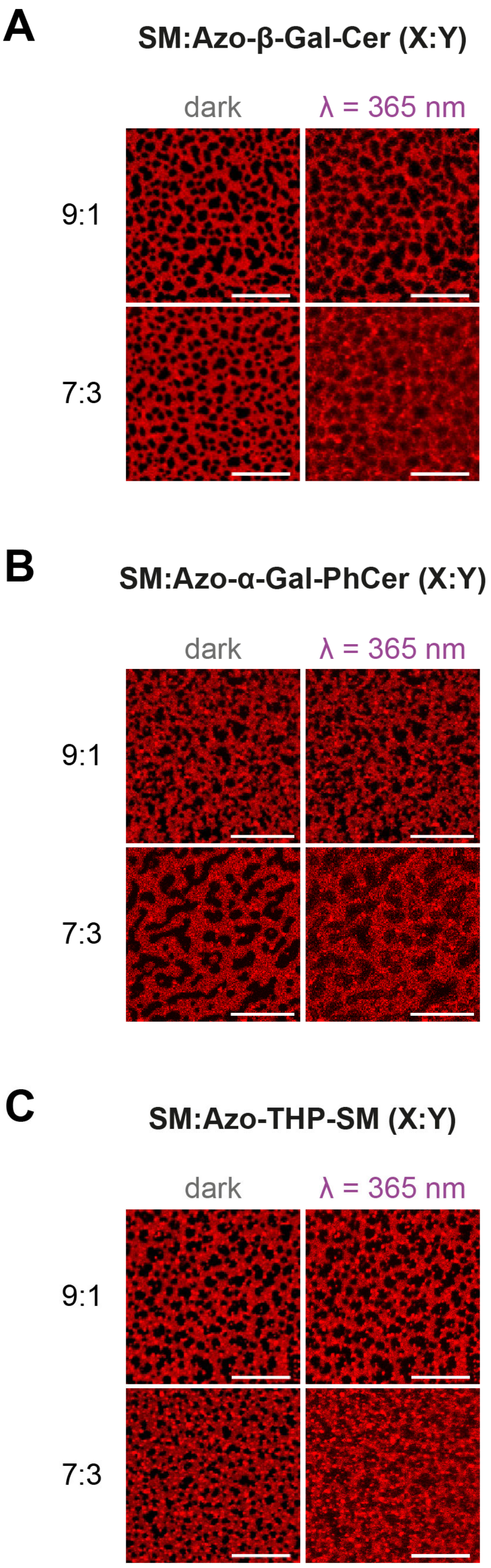

**Fig. S3 – Admixing of $L_d$-$L_o$ directly after irradiation with UV-A light on membranes with lower amounts of azo-sphingolipids.** Fluorescence confocal images of SLBs composed of DOPC:Chol:SM:photolipid (10:6.7:X:Y mol ratio), doped with 0.1 mol% Atto655-DOPE (for fluorescence detection of $L_d$ phase), before and directly after illumination with UV-A light (λ = 365 nm). Samples with (A) Azo-β-Gal-Cer, (B) Azo-α-Gal-PhCer and (C) Azo-THP-SM at 3.7 mol% (SM:photolipid, X:Y = 9:1) or 11.2 mol% (SM:photolipid, X:Y = 7:3). Scale-bar is 20 μm.

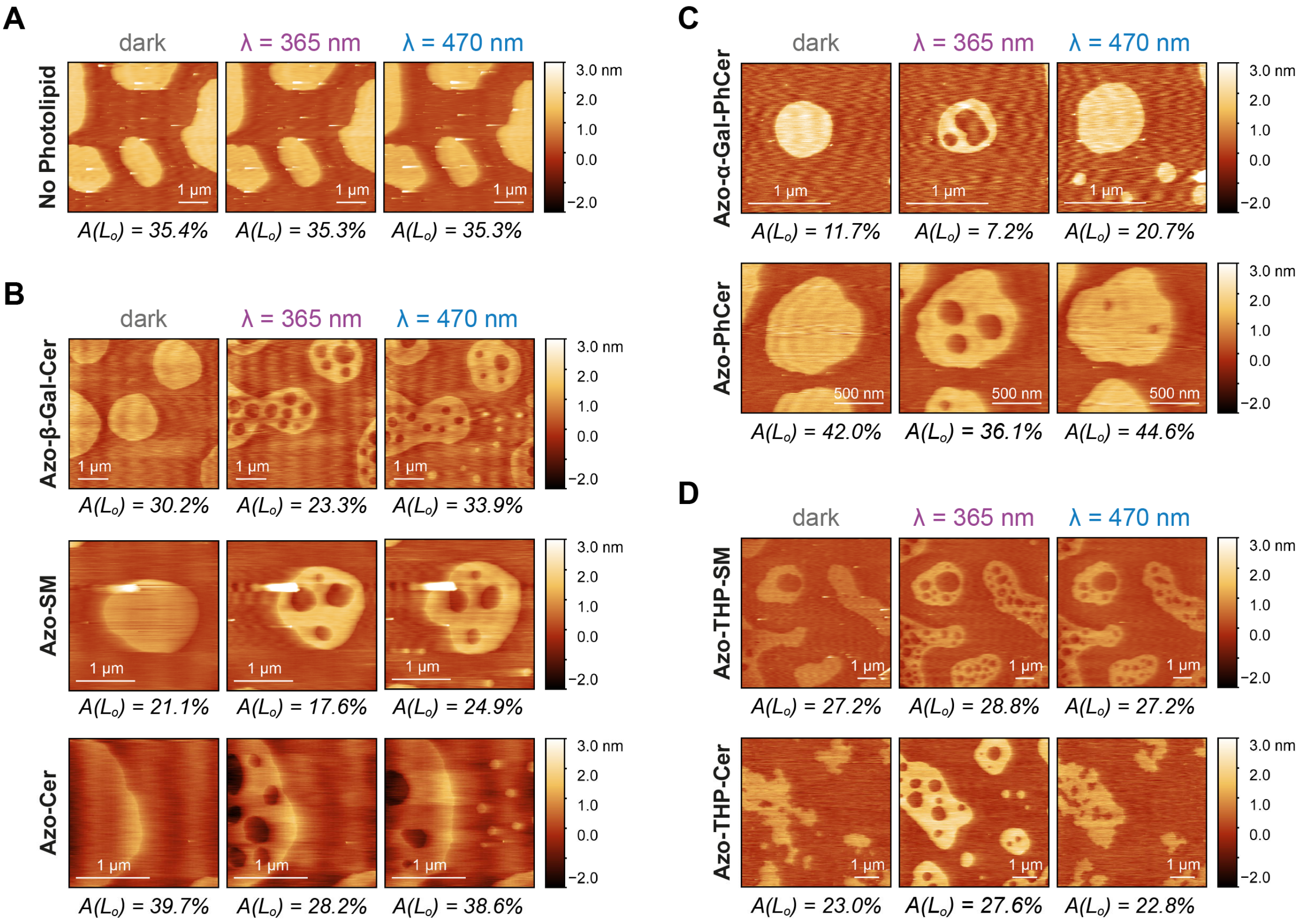

**Fig. S4 – Additional high-speed AFM height images of phase-separated supported bilayers containing different types of azo-sphingolipids upon light trigger**. Changes in the area of $L_o$ domains before/after illumination with UV-A (λ = 365 nm) and blue (λ = 470 nm) lights on DOPC:Chol:SM:photolipid (10:6.7:5:5 mol ratio) SLBs having different types of photoswitchable lipids. ***(A) without azo-sphingolipid***: control with SM; mixture being DOPC:Chol:SM (10:6.7:10 mol ratio). ***(B) with spingosine-based azo-sphingolipids***: Azo-β-Gal-Cer, Azo-SM or Azo-Cer. ***(C) with phytospingosine-based azo-sphingolipids***: Azo-α-Gal-PhCer or Azo-PhCer. ***(D) samples with 3-OH-blocked azo-sphingolipids***: Azo-THP-SM or Azo-THP-Cer.

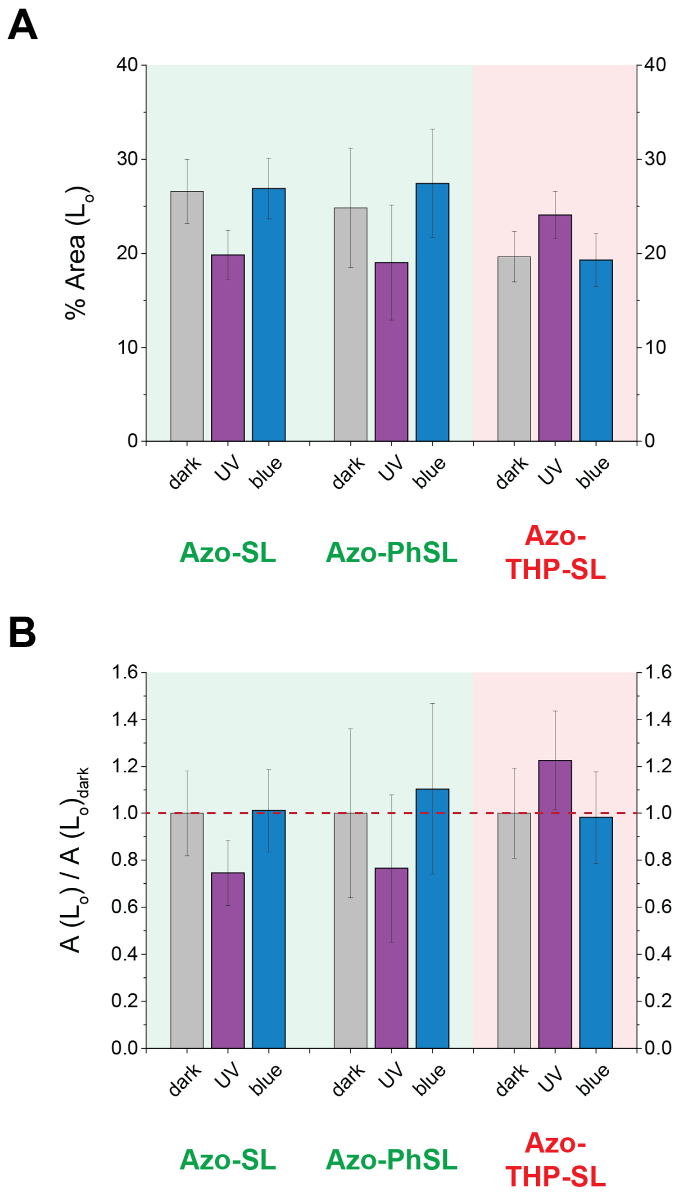

**Fig. S5 – Average L$_o$ areas of phase-separated SLBs containing photoswitchable sphingolipids, grouped by type, recovered from high-speed AFM images in Figs. 2, 3 and S4.** Non-normalized (A) and normalized (B) L$_o$ areas of grouped SLBs containing azo-sphingolipids Azo-β-Gal-Cer, Azo-SM and Azo-Cer (Azo-SL), azo-phytosphingolipids Azo-α-Gal-PhCer and Azo-PhCer (Azo-PhSL), and THP-protected azo-sphingolipids Azo-THP-SM and Azo-THP-Cer (Azo-THP-SL). Grey bars corresponds to average values at the dark-adapted state, purple bars to the average values after irradiation with UV-A light (λ = 365 nm), and blue bars to values after irradiation with blue light (λ = 470 nm). Columns relative to photolipids with free 3-OH (i.e. Azo-SL and Azo-PhSL) are marked in green, while columns relative to photolipids with blocked-3-OH (i.e. Azo-THP-SL) are marked in red. Error bars correspond to standard error of the mean (*n* = 4-7).

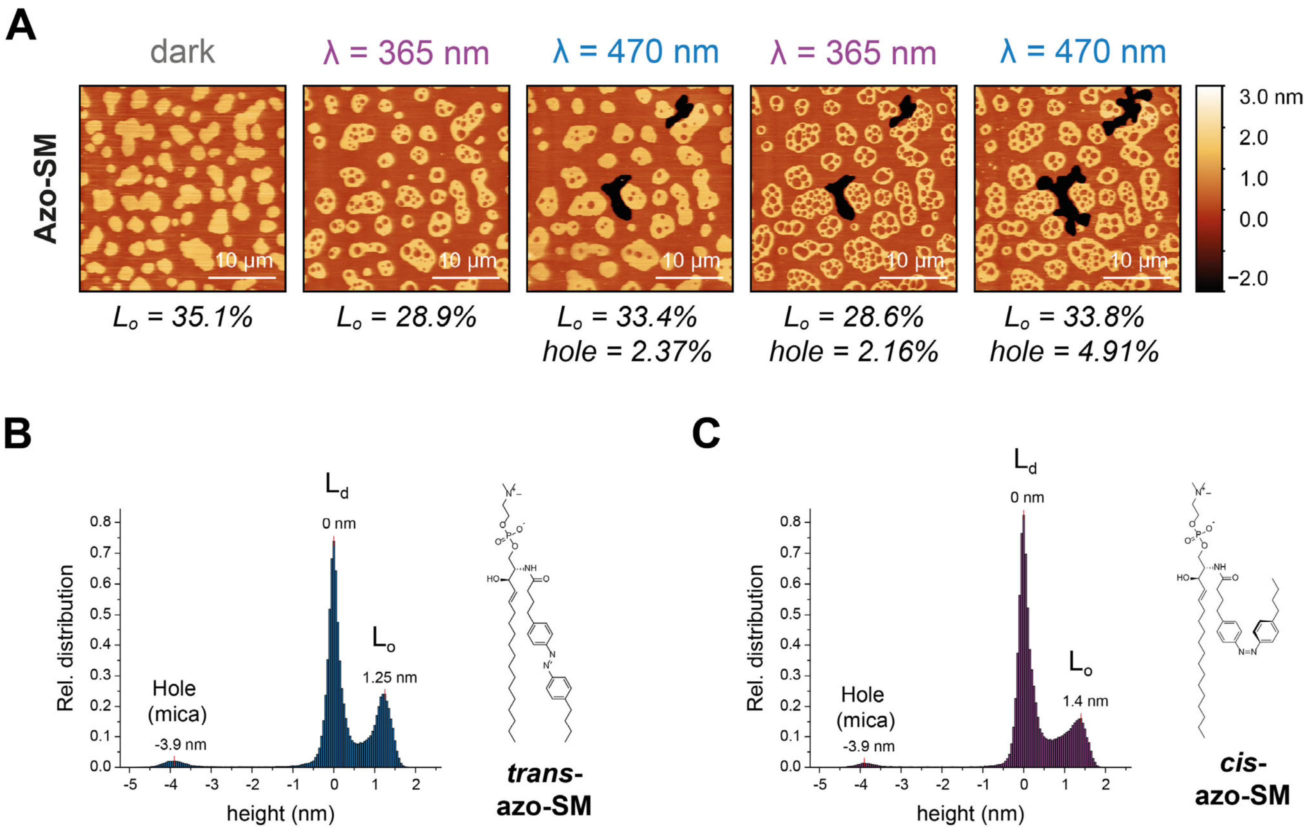

**Fig. S6 – Reshuffling of $L_d$-$L_o$ phase separation and membrane expansion/compaction triggered by the photo-isomerization of Azo-SM.** (A) Sequential AFM images of DOPC:Chol:SM:Azo-SM (10:6.7:5:5 mol ratio) SLB undergoing phase reshuffling and hole expansion/compaction upon applying UV-A (λ = 365 nm) and blue (λ = 470 nm) lights. Areas of $L_o$ phase and membrane holes are additionally depicted. (B, C) Height distribution histograms extracted from images above when Azo-SM was in the (B) *trans*- (blue-adapted) and (C) *cis*- (UV-adapted) states, respectively. Peaks correspond to the height level of the holes (mica surface, -3.9 nm), $L_d$ (0 nm) and $L_o$ (1.25 / 1.4 nm) phases, as marked.

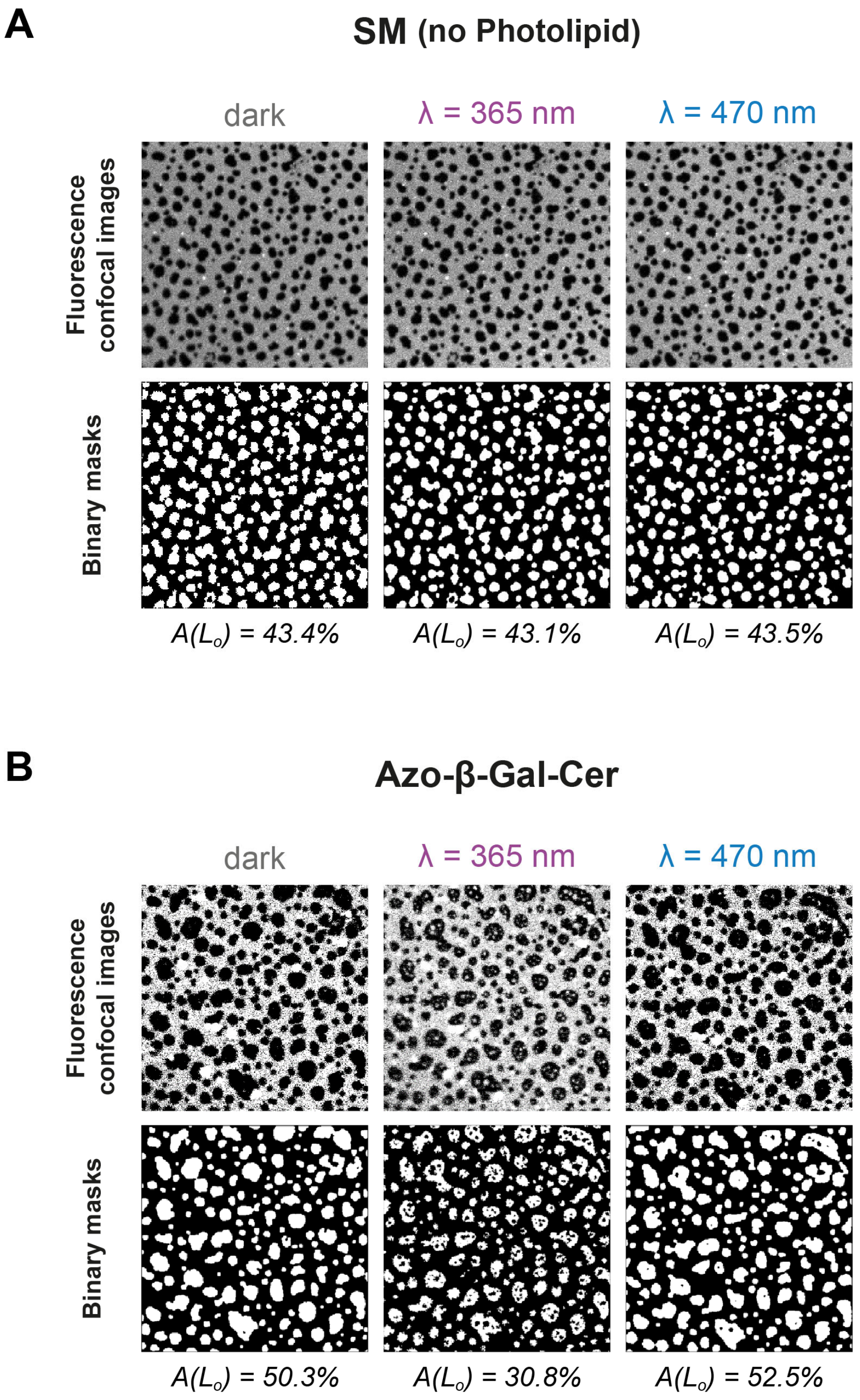

**Fig. S7 – Quantification of the amount of $L_d$-$L_o$ area from fluorescence confocal data.** Example of fluorescence confocal images, generated binary masks and recovered $L_o$ phase area values for (A) DOPC:Chol:SM (10:6.7:10) control SLBs lacking photoswitchable sphingolipids, as well as (B) DOPC:Chol:SM:Azo-β-GalCer (10:6.7:5:5) SLBs having a sphingosine-based photoswitchable sphingolipid, prior/after illumination with UV-A (λ = 365 nm) and blue (λ = 470 nm) lights. Microscopy images correspond to large fields-of-view (56.7 × 56.7 μm$^2$) of SLBs having 0.1 mol% Atto655-DOPE for fluorescence detection of the $L_d$ phase.

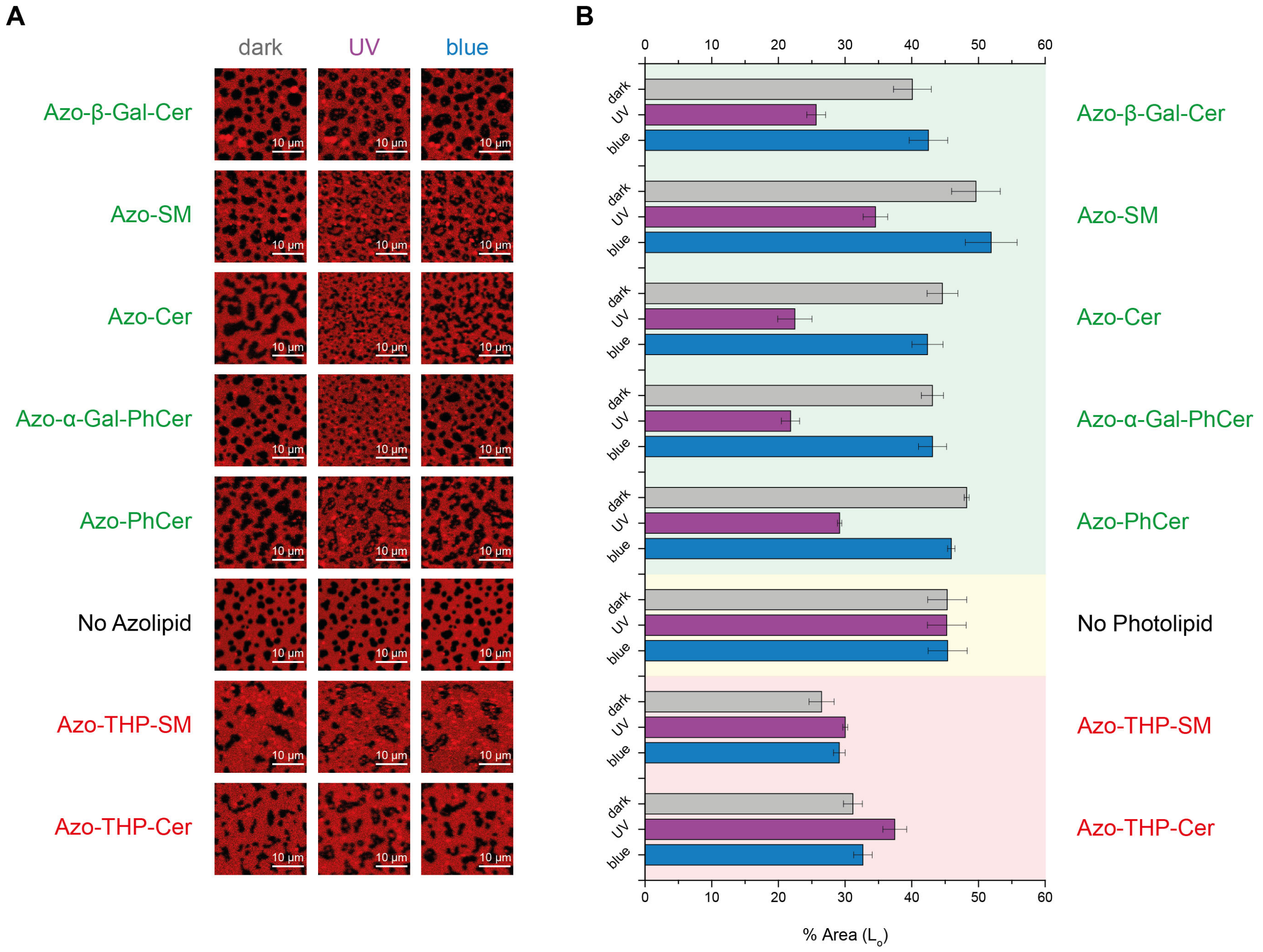

**Fig. S8 – Percentage of $L_o$ phase area retrieved from florescence confocal images prior/after illumination with UV-A (λ = 365 nm) and blue (λ = 470 nm) lights.** (A) Microscopy images of DOPC:Chol:SM:photolipid (10:6.7:5:5 mol ratio) and control (no photolipid; 10:6.7:10 mol ratio) SLBs, doped with 0.1 mol% Atto655-DOPE for fluorescence detection. (B) Average percentage of $L_o$ phase retrieved for phase-separated SLBs containing either azo-(phyto)sphingolipids with free 3-OH (marked in green), no photolipid (controls with SM, marked in yellow), or THP-protected azo-sphingolipids with the 3-OH blocked (marked in red). Error bars correspond to the standard error of the mean (*n* = 5-8).

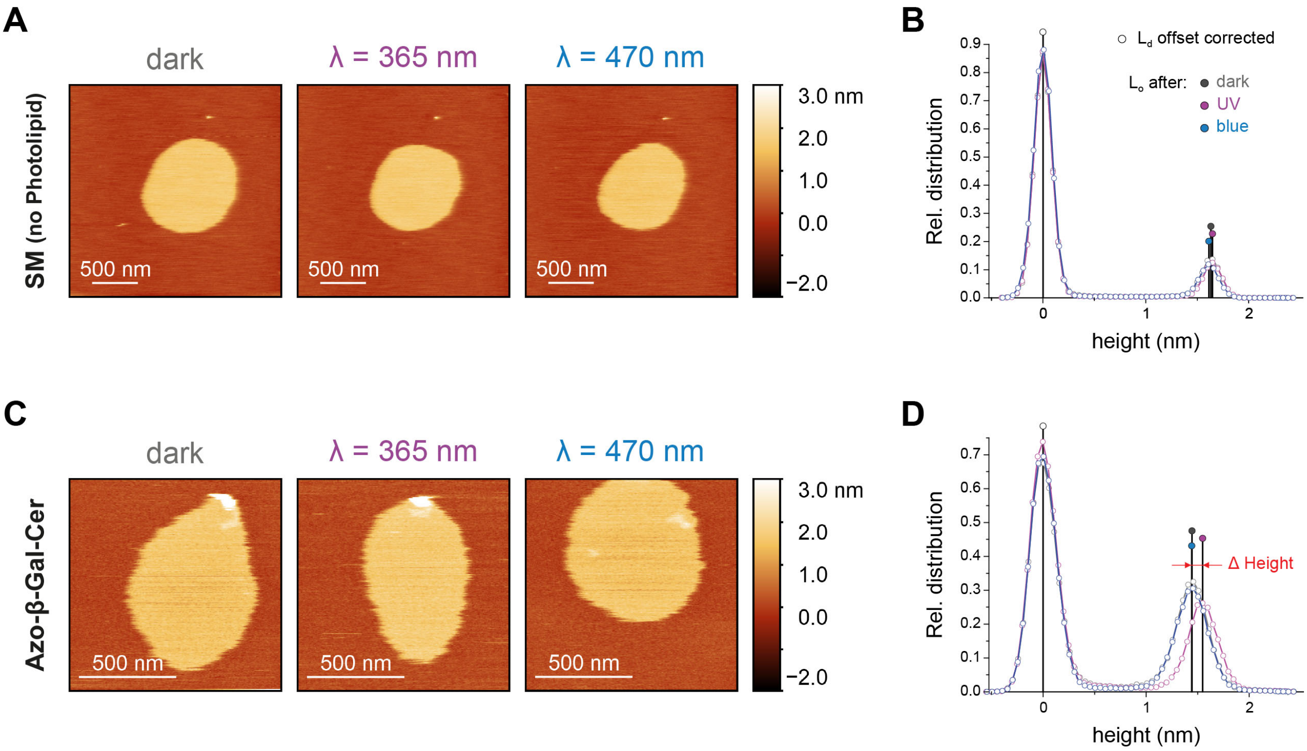

**Fig. S9 – Quantification of the $L_d$-$L_o$ height difference from AFM data. (A, C)** Example of AFM height images for phase-separated (A) DOPC:Chol:SM (10:6.7:10) control SLBs lacking photoswitchable sphingolipids, as well as (C) DOPC:Chol:SM:Azo-β-GalCer (10:6.7:5:5) SLBs having a sphingosine-based photoswitchable sphingolipid, prior/after illumination with UV-A (λ = 365 nm) and blue (λ = 470 nm) lights. **(B, D)** Height distribution profiles for the displayed membranes lacking photolipid (B) and with Azo-β-GalCer (D) at the dark-, UV- and blue-adapted states, with the $L_d$ (centered at 0 nm) and $L_o$ (1.4-1.7 nm) phase height peaks accordingly marked.

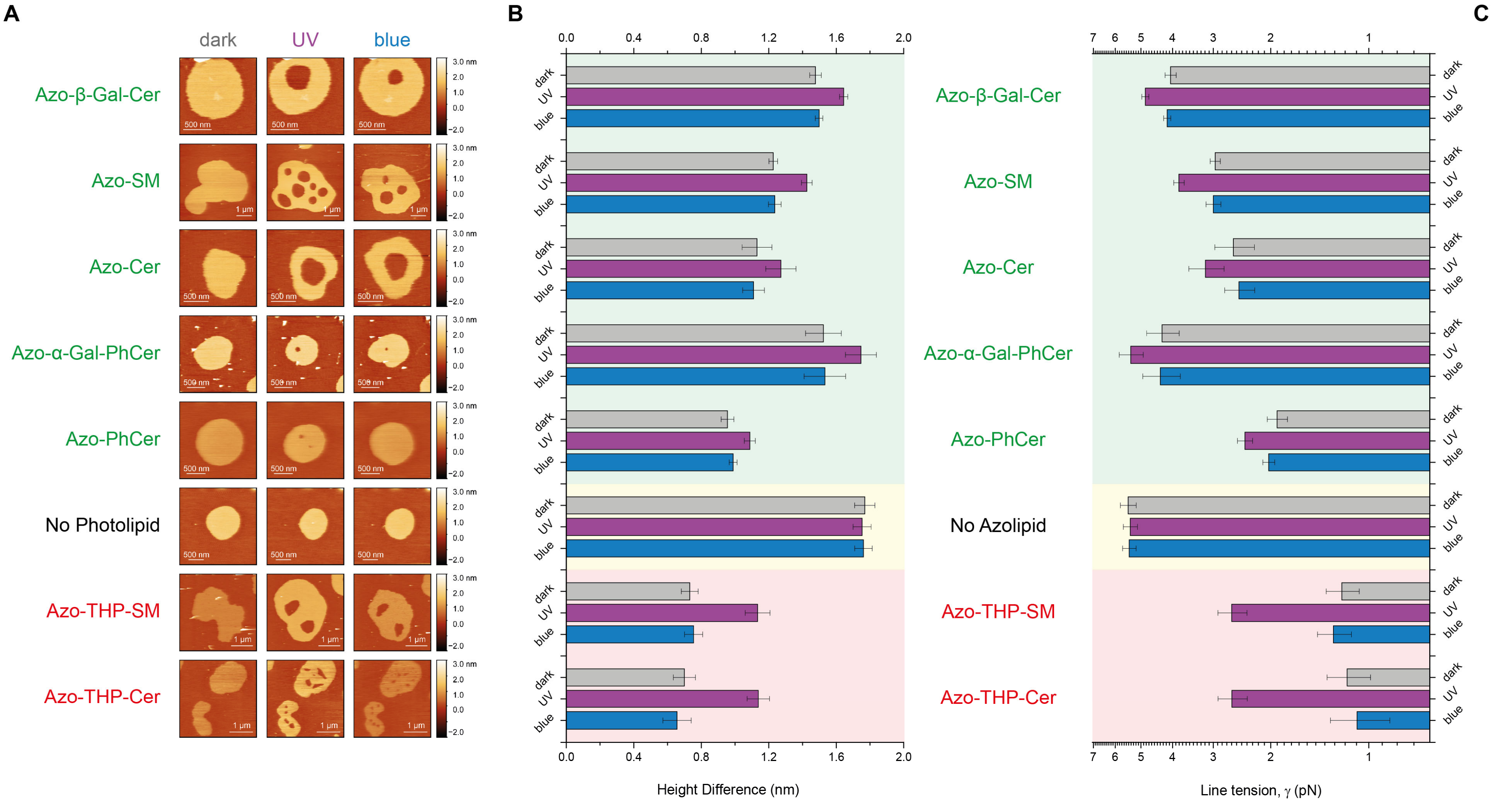

**Fig. S10 – $L_d$-$L_o$ height difference and line tension values retrieved from AFM images prior/after illumination with UV-A (λ = 365 nm) and blue (λ = 470 nm) lights.** (A) Slow-speed AFM height images of DOPC:Chol:SM:photolipid (10:6.7:5:5 mol ratio) and control (no photolipid; 10:6.7:10 mol ratio) SLBs. Average height mismatches (B) and calculated line tension values for phase-separated SLBs containing either azo-(phyto)sphingolipids with free 3-OH (marked in green), no photolipid (controls with SM, marked in yellow), or THP-protected azo-sphingolipids with the 3-OH blocked (marked in red). Error bars correspond to standard error of the mean (*n* = 5-9).

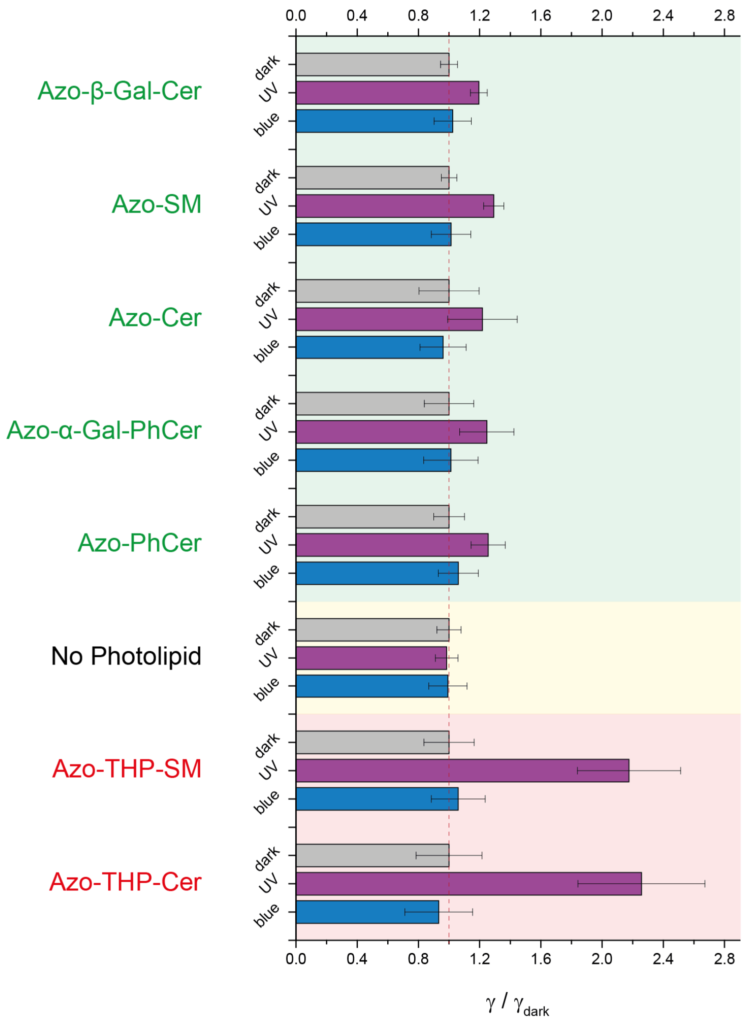

**Fig. S11 – Normalized changes in the line tension values of $L_o$ domains on phase-separated membranes with different types of azo-sphingolipids, upon application of UV-A (λ = 365 nm) and blue (λ = 470 nm) light.** Average line tension values (normalized to dark-adapted state) calculated from height mismatches in Fig. S10 for SLBs containing either azo-(phyto)sphingolipids with free 3-OH (marked in green), no photolipid (controls with SM, marked in yellow), or THP-protected azo-sphingolipids with the 3-OH blocked (marked in red). Error bars correspond to the standard error of the mean (*n* = 5-9).

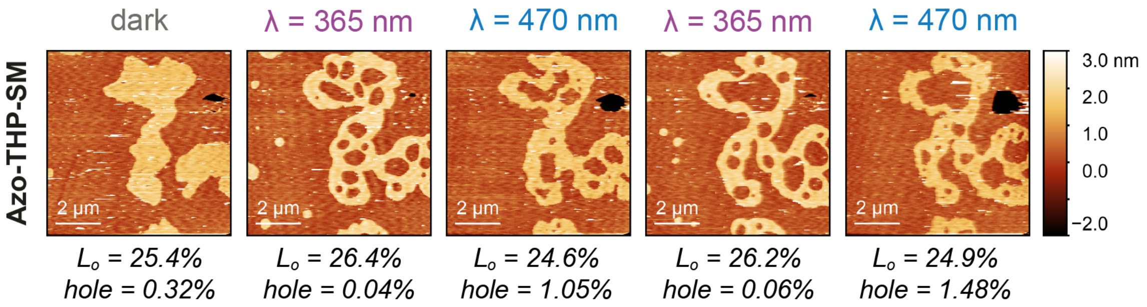

**Fig. S12 – Membrane expansion/compaction triggered by the photo-isomerization of Azo-THP-SM.** Sequential AFM images of DOPC:Chol:SM:Azo-THP-SM (10:6.7:5:5 mol ratio) SLB undergoing phase reshuffling and hole expansion/compaction upon applying UV-A (λ = 365 nm) and blue (λ = 470 nm) lights. Areas of $L_o$ phase and membrane holes are additionally depicted.

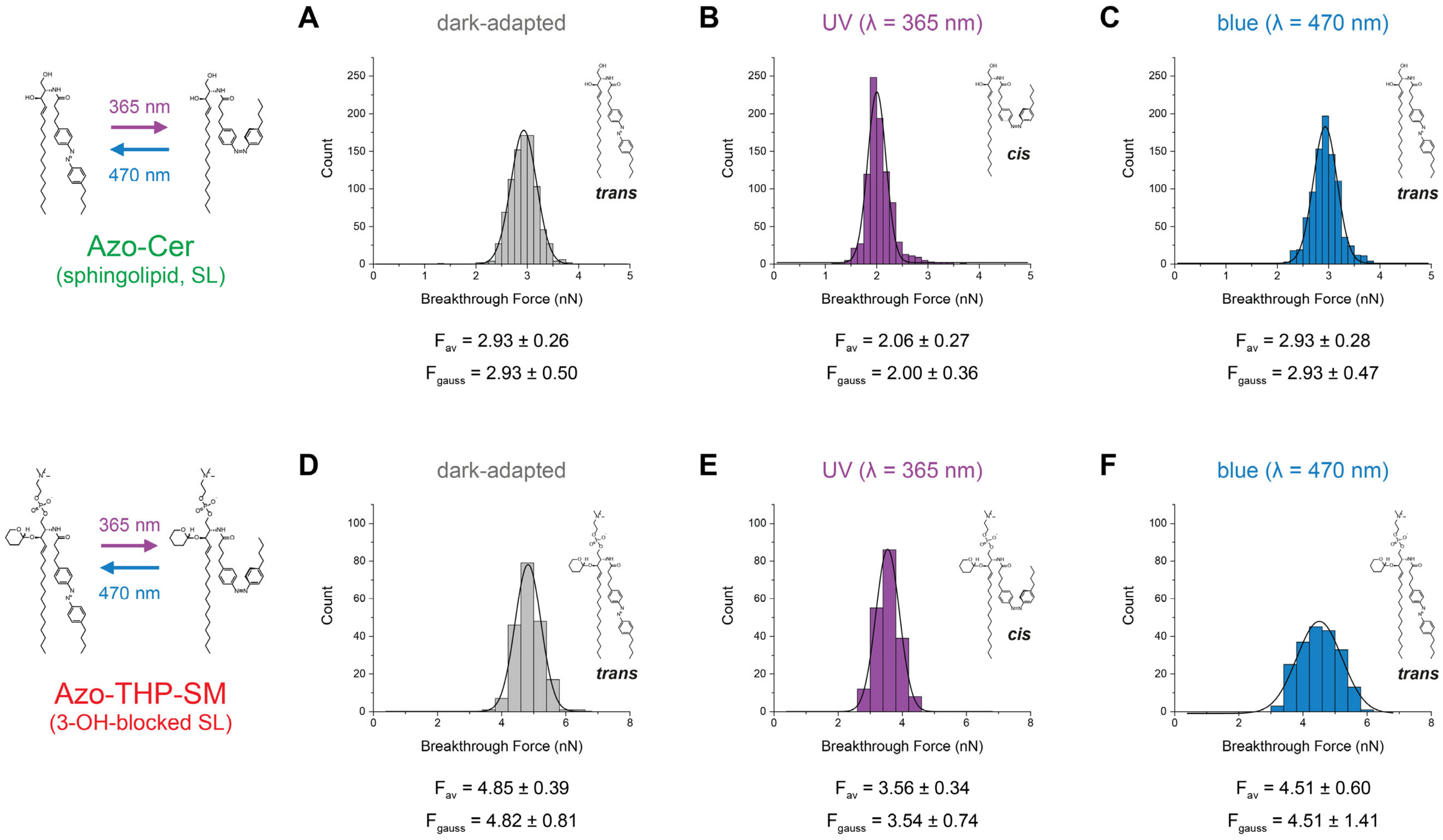

**Fig. S13 – Non-normalized histograms of breakthrough forces obtained via force spectroscopy on homogeneous SLBs containing non-blocked Azo-Cer (*A-C*) and 3-OH-blocked Azo-THP-SM (*D-F*) azo-sphingolipids.** Nominal breakthrough forces recovered from membrane piercing experiments on (A-C) DOPC:Chol:Azo-Cer and (D-F) DOPC:Chol:Azo-THP-SM homogenous membranes (at 10:6.7:10 mol ratio) at the (A, D) dark-, (B, E) UV light- and (C, F) blue light-adapted states. Average breakthrough forces (± standard deviation), as well as Gaussian peak fits (± half-width) are displayed.

#### Supplementary movie legends

**Movie S1 – Remodeling of lipid domains by the sphingosine-based Azo-β-GalCer on a phase-separated SLB made of DOPC:Chol:SM:Azo-β-GalCer (10:6.7:5:5 mol ratio), recorded using high-speed AFM.** Images correspond to height signal. Initial dark-adapted state is marked with a gray circle. Isomerization to *cis*-Azo-β-GalCer upon irradiation with UV-A light (λ = 365 nm) is marked with a purple circle. Isomerization back to *trans*-Azo-β-GalCer upon irradiation with blue light (λ = 470 nm) is marked with a blue circle. Acquisition = 3.2 s/frame. Video frame rate = 11 fps.

**Movie S2 – Remodeling of lipid domains by the sphingosine-based Azo-SM on a phase-separated SLB made of DOPC:Chol:SM:Azo-SM (10:6.7:5:5 mol ratio), recorded using high-speed AFM.** Images correspond to height signal. Initial dark-adapted state is marked with a gray circle. Isomerization to *cis*-Azo-SM upon irradiation with UV-A light (λ = 365 nm) is marked with a purple circle. Isomerization back to *trans*-Azo-SM upon irradiation with blue light (λ = 470 nm) is marked with a blue circle. Acquisition = 16.2 s/frame. Video frame rate = 11 fps.

**Movie S3 – Remodeling of lipid domains by the sphingosine-based Azo-Cer on a phase-separated SLB made of DOPC:Chol:SM:Azo-Cer (10:6.7:5:5 mol ratio), recorded using high-speed AFM.** Images correspond to height signal. Initial dark-adapted state is marked with a gray circle. Isomerization to *cis*-Azo-Cer upon irradiation with UV-A light (λ = 365 nm) is marked with a purple circle. Isomerization back to *trans*-Azo-Cer upon irradiation with blue light (λ = 470 nm) is marked with a blue circle. Acquisition = 5.2 s/frame. Video frame rate = 11 fps.

**Movie S4 – Reversible remodeling of lipid domains by Azo-SM on a phase-separated SLB made of DOPC:Chol:SM:Azo-SM (10:6.7:5:5 mol ratio), recorded using high-speed AFM.** Images correspond to height signal. Initial dark-adapted state is marked with a gray circle. Isomerization to *cis*-Azo-SM upon irradiation with UV-A light (λ = 365 nm) is marked with a purple circle. Isomerization back to *trans*-Azo-SM upon irradiation with blue light (λ = 470 nm) is marked with a blue circle. Acquisition = 20.2 s/frame. Video frame rate = 11 fps. Quantification of the variation in $L_o$ area is depicted in Fig. 2D.

**Movie S5 – Remodeling of lipid domains by the phytosphingosine-based Azo-α-Gal-PhCer on a phase-separated SLB made of DOPC:Chol:SM:Azo-α-Gal-PhCer (10:6.7:5:5 mol ratio), recorded using high-speed AFM.** Images correspond to height signal. Initial dark-adapted state is marked with a gray circle. Isomerization to *cis*-Azo-α-Gal-PhCer upon irradiation with UV-A light (λ = 365 nm) is marked with a purple circle. Isomerization back to *trans*-Azo-α-Gal-PhCer upon irradiation with blue light (λ = 470 nm) is marked with a blue circle. Acquisition = 5.2 s/frame. Video frame rate = 11 fps.

**Movie S6 – Remodeling of lipid domains by the phytosphingosine-based Azo-PhCer on a phase-separated SLB made of DOPC:Chol:SM:Azo-PhCer (10:6.7:5:5 mol ratio), recorded using high-speed AFM.** Images correspond to height signal. Initial dark-adapted state is marked with a gray circle. Isomerization to *cis*-Azo-PhCer upon irradiation with UV-A light (λ = 365 nm) is marked with a purple circle. Isomerization back to *trans*-Azo-PhCer upon irradiation with blue light (λ = 470 nm) is marked with a blue circle. Acquisition = 5.2 s/frame. Video frame rate = 11 fps.

**Movie S7 – Remodeling of lipid domains by the 3-OH-blocked sphingosine-based Azo-THP-SM on a phase-separated SLB made of DOPC:Chol:SM:Azo-THP-SM (10:6.7:5:5 mol ratio), recorded using high-speed AFM.** Images correspond to phase signal. Initial dark-adapted state is marked with a gray circle. Isomerization to *cis*-Azo-THP-SM upon irradiation with UV-A light (λ = 365 nm) is marked with a purple circle. Isomerization back to *trans*-Azo-THP-SM upon irradiation with blue light (λ = 470 nm) is marked with a blue circle. Acquisition = 4.1 s/frame. Video frame rate = 11 fps.

**Movie S8 – Remodeling of lipid domains by the 3-OH-blocked sphingosine-based Azo-THP-Cer on a phase-separated SLB made of DOPC:Chol:SM:Azo-THP-Cer (10:6.7:5:5 mol ratio), recorded using high-speed AFM.** Images correspond to height signal. Initial dark-adapted state is marked with a gray circle. Isomerization to *cis*-Azo-THP-Cer upon irradiation with UV-A light (λ = 365 nm) is marked with a purple circle. Isomerization back to *trans*-Azo-THP-Cer upon irradiation with blue light (λ = 470 nm) is marked with a blue circle. Acquisition = 16.2 s/frame. Video frame rate = 11 fps.

**Movie S9 – Reversible remodeling of lipid domains by Azo-THP-SM on a phase-separated SLB made of DOPC:Chol:SM:Azo-THP-SM (10:6.7:5:5 mol ratio), recorded using high-speed AFM.** Images correspond to height signal. Initial dark-adapted state is marked with a gray circle. Isomerization to *cis*-Azo-THP-SM upon irradiation with UV-A light (λ = 365 nm) is marked with a purple circle. Isomerization back to *trans*-Azo-THP-SM upon irradiation with blue light (λ = 470 nm) is marked with a blue circle. Acquisition = 20.2 s/frame. Video frame rate = 11 fps. Quantification of the variation in $L_o$ area is depicted in Fig. 3C.

### Compound synthesis and characterization

#### Methods and equipment

Unless otherwise noted, all reactions were magnetically stirred and performed under an atmosphere of inert gas (Ar or $N_2$) using standard Schlenk techniques. The reactions were carried out in oven-dried glassware (200 °C oven temperature). External bath temperatures were used to record all reaction mixture temperatures. Diethyl ether ($Et_2O$) and tetrahydrofuran (THF) were distilled prior to use under an atmosphere of $N_2$ from sodium and benzophenone, triethylamine ($NEt_3$) from calcium hydride. *N*,*N*-dimethylformamide (DMF), toluene and methanol (MeOH) were purchased from Acros Organics as 'extra dry' reagents under inert gas atmosphere and over molecular sieves. Solvents for flash column chromatography and crystallization experiments were purchased in technical grade and distilled under reduced pressure prior to use. Degassed solvents were degassed under $N_2$ atmosphere by using either three successive freeze-pump-thaw cycles or by purging the solvent for 30 min with $N_2$. Petroleum ether (PE) refers to fractions of *iso*-hexanes which boil between 40 and 80 °C. All other reagents were purchased from commercial sources and used without further purification.

**Chromatography**. Analytical thin-layer chromatography (TLC) was performed on pre-coated glass plates (silica gel 60 F254) from Merck, and visualized by exposure to ultraviolet light (UV, 254 nm) and by staining with aqueous acidic ceric ammonium molybdate(IV) (CAM) solution. Flash column chromatography was performed using Merck silica gel (40–63 µm particle size).

**NMR Spectroscopy.** Proton nuclear magnetic resonance ($^1H$ NMR) spectra were recorded in 5 mm tubes on a Varian 300, Varian 400, Inova 400 or Varian 600 spectrometer in deuterated solvents at room temperature. Chemical shifts ($\delta$ scale) are expressed in parts per million (ppm) and are calibrated using residual protic solvent as an internal reference ($CHCl_3$: $\delta$ = 7.26 ppm). Data for $^1H$ NMR spectra are reported as follows: chemical shift ($\delta$ ppm) (multiplicity, coupling constants (Hz), integration). Couplings are expressed as: s = singlet, d = doublet, t = triplet, q = quartet, m = multiplet or combinations thereof. Carbon nuclear magnetic resonance ($^{13}C$ NMR) spectra were recorded at 75, 100 and 150 MHz, respectively. Carbon chemical shifts ($\delta$ scale) are also expressed in parts per million (ppm) and are referenced to the central carbon resonances of the solvents ($CDCl_3$: $\delta$ = 77.16 ppm). In order to assign the $^1H$ and $^{13}C$ NMR spectra, a range of 2D NMR experiments (COSY, HSQC, HMBC, NOESY) were used as appropriate. The numbering of the proton and carbon atoms does not correspond to the IUPAC nomenclature. Diastereotopic protons in the $^1H$ NMR spectra are referenced with a and b, nomenclature is arbitrarily and does not correspond to the spin system.

**High performance liquid chromatography (HPLC)**. HPLC was performed with HPLC grade solvents and deionized $H_2O$ that was purified on a TKA MicroPure $H_2O$ purification system. All solvents were degassed with helium gas prior to use. Unless noticed otherwise, all experiments were carried out at room temperature.

Analytical HPLC spectra were recorded on a ultra-high performance liquid chromatography (UHPLC) system from the Agilent 1260 Infinity series (1260 degasser, 1260 Binary Pump VL, 1260 ALS auto sampler, 1260 TCC thermostated column compartment, 1260 DAD diode array detector), which was computer-controlled through Agilent ChemStation software.

Chiral HPLC spectra were recorded on a high performance liquid chromatography (HPLC) system from the Shimadzu 20A series (DGU-20A3R degasser, LC-20AD Binary Pump VL, SIL-20AHT autosampler, CTO-20A thermostated column compartment, SPD-M20A DAD diode array detector), which was computer controlled through Shimadzu LabSolutions Software (Version 5.42 SP5). Enantiomeric excess (*ee*) was calculated by using the following equation; $m_1$ refers to the integral of the major peak and $m_2$ to the integral of the minor peak:

$$ee = \frac{|\text{-}m_1 - m_2|}{m_1 + m_2} \cdot 100\%$$

**High-resolution mass spectrometry (HRMS).** A Varian MAT CH7A mass spectrometer was used to obtain high-resolution electron ionization (EI) mass. High-resolution electronspray (ESI) mass spectra were recorded on a Varian MAT 711 MS spectrometer operation in either positive or negative ionization modes.

**Infrared spectroscopy (IR).** Infrared spectra (IR) were recorded on a Perkin Elmer Spectrum BX II (FTIR System) equipped with an attenuated total reflection (ATR) measuring unit. IR data is reported in frequency of absorption ($cm^{-1}$). The IR bands are characterized as: w = weak, m = medium, s = strong, br = broad, or combinations thereof.

**Melting points (mp)**. Melting points were measured on a Büchi Melting Point B-540 or SRS MPA120 EZ-Melt apparatus and are uncorrected.

**Optical rotation**. Perkin-Elmer 241 or Krüss P8000-T polarimeter were used to measure optical rotation at the Sodium D-line (589 nm) at the given temperature (*T* in °C) and concentrations (*c* in g/100 mL) using spectroscopic grade solvents. The measurements were carried out in a cell with a path length (*d*) of 0.5 dm. Specific rotations were calculated using the following equation:

$$[\alpha]_D = \frac{\alpha}{c \cdot d} \frac{10^{-1} \cdot \text{deg} \cdot \text{cm}^2}{\text{g}}$$

#### Experimental procedures

**(2*S*,3*R*,*E*)-2-(4-(4-((*E*)-(4-butylphenyl)diazenyl)phenyl)butanamido)-3-hydroxyoctadec-4-en-1-yl benzoate (SI1)**

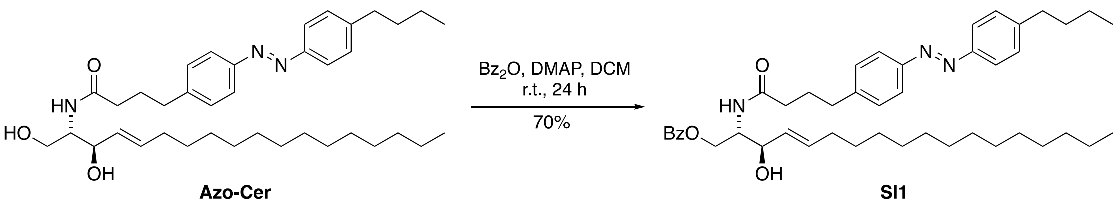

To a solution of **Azo-Cer** (previously named ACe-1[R1]) (100 mg, 0.165 mmol, 1.0 eq.) in dry DCM (10 mL) were added benzoic anhydride (41.2 mg, 0.182 mmol, 1.11 eq.) and DMAP (2.02 mg, 16.5 µmol, 0.10 eq.) and the reaction was stirred for 24 h at room temperature. Then, $NaHCO_3$ solution (100 mL) was added and the organic layer was extracted with EtOAc (3 × 100 mL) and dried ($Na_2SO_4$). Purification *via* flash column chromatography [PE/EtOAc, 9:1 to 1:2] afforded benzoyl-protected ceramide **SI1** (81.7 mg, 0.115 mmol, 70%) as a light orange solid.

$\mathbf{R}_f$ = 0.69 [PE: EtOAc 1:1]

**$^1$H NMR** (400 MHz, $CDCl_3$): $\delta$ = 8.04–7.97 (m, 2H), 7.84–7.76 (m, 4H), 7.59 – 7.53 (m, 1H), 7.42 (t, *J* = 7.7 Hz, 2H), 7.35–7.28 (m, 2H), 7.25 (d, *J* = 8.1 Hz, 2H), 5.93 (d, *J* = 7.9 Hz, 1H), 5.77 (dtd, *J* = 14.9, 6.7, 1.2 Hz, 1H), 5.53 (ddt, *J* = 15.4, 6.6, 1.5 Hz, 1H), 4.62–4.53 (m, 1H), 4.47–4.37 (m, 2H), 4.26 (dd, *J* = 6.7, 4.0 Hz, 1H), 2.69 (t, *J* = 7.9 Hz, 4H), 2.22 (t, *J* = 7.4 Hz, 2H), 2.00 (m, 4H), 1.70–1.59 (m, 2H), 1.45–1.15 (m, 24H), 0.94 (t, *J* = 7.3 Hz, 3H), 0.87 (t, *J* = 6.8 Hz, 3H) ppm.

**$^{13}$C NMR** (101 MHz, $CDCl_3$): $\delta$ = 173.2, 167.1, 151.4, 151.1, 146.5, 144.6, 135.0, 133.5, 129.9, 129.3, 129.2, 128.7, 128.2, 123.0, 122.9, 73.5, 63.4, 53.5, 36.0, 35.74, 35.1, 33.6, 32.4, 32.1, 29.8, 29.8, 29.8, 29.6, 29.5, 29.4, 29.2, 27.0, 22.9, 22.5, 14.3, 14.1 ppm.

**IR** (ATR): $\tilde{\nu}$ = 2925 (s), 2853 (m), 2358 (w), 2340 (w), 1722 (m), 1648 (m), 1602 (w), 1498 (w), 1452 (w), 1388 (w), 1273 (s), 965 (w), 844 (w), 712 (m) $cm^{-1}$.

**HRMS** (ESI): calcd. for $C_{45}H_{64}N_3O_4^+$ : 710.4891 $[M+H]^+$
found: 710.4887 $[M+H]^+$.

**(2*S*,3*R*,*E*)-2-(4-(4-((*E*)-(4-butylphenyl)diazenyl)phenyl)butanamido)-3-((tetrahydro-2H-pyran-2-yl)oxy)octadec-4-en-1-yl benzoate (SI2)**

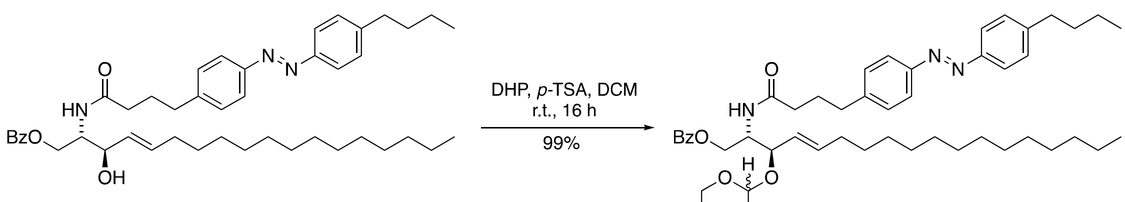

**SI1** **SI2**

To a solution of secondary alcohol **SI1** (75.4 mg, 0.106 mmol, 1.0 eq.) in dry DCM (5 mL) were added freshly distilled dihydropyran (42.9 mL, 0.475 mmol, 4.5 eq.) and *p*-TSA monohydrate (1.63 mg, 9.49 µmol, 0.09 eq.) and the reaction mixture was stirred for 16 h at room temperature. Then, $NaHCO_3$ solution (10 mL) was added and the aqueous layer was extracted with EtOAc (3 × 20 mL) and dried ($Na_2SO_4$). Purification *via* flash column chromatography [PE/EtOAc, 6:1 to 3:1] afforded a diastereomeric mixture of benzoyl-protected ceramide **SI2** (83.0 mg, 0.105 mmol, 99%) as a light orange solid.

$\mathbf{R_f}$ = 0.45 [Pent: EtOAc 3:1]

**$^1$H NMR** (400 MHz, $CDCl_3$): $\delta$ = 8.06–8.01 (m, 2H), 7.84–7.76 (m, 4H), 7.56–7.50 (m, 1H), 7.44–7.37 (m, 2H), 7.33–7.28 (m, 2H), 7.25–7.22 (m, 2H), 6.27 (d, $J$ = 8.2 Hz, 1H), 5.79–5.70 (m, 1H), 5.39 (ddt, $J$ = 15.4, 7.2, 1.5 Hz, 1H), 4.57 (dtd, $J$ = 8.8, 4.2, 1.9 Hz, 1H), 4.54–4.42 (m, 3H), 4.24 (dd, $J$ = 7.3, 4.0 Hz, 1H), 3.95 (dq, $J$ = 11.2, 2.4 Hz, 1H), 3.86 (ddp, $J$ = 10.5, 5.9, 2.8 Hz, 1H), 3.74 (dddt, $J$ = 16.5, 14.4, 6.9, 4.1 Hz, 1H), 3.56–3.46 (m, 1H), 3.46–3.29 (m, 1H), 2.67 (td, $J$ = 7.9, 5.6 Hz, 4H), 2.19 (td, $J$ = 7.2, 2.4 Hz, 2H), 2.00 (dq, $J$ = 22.1, 7.2 Hz, 4H), 1.80 (d, $J$ = 5.8 Hz, 2H), 1.75–1.59 (m, 4H), 1.59–1.44 (m, 6H), 1.43–1.18 (m, 24H), 0.94 (t, $J$ = 7.3 Hz, 3H), 0.89–0.84 (m, 3H) ppm.

**$^{13}$C NMR** (101 MHz, $CDCl_3$): $\delta$ = 172.2, 166.8, 151.3, 151.1, 146.4, 144.9, 136.3, 133.1, 129.8, 129.2, 129.2, 128.5, 126.3, 122.9, 122.8, 98.8, 97.2, 78.1, 77.2, 68.4, 67.8, 67.6, 63.7, 63.6, 62.4, 51.6, 36.2, 35.7, 35.1, 33.6, 32.4, 32.0, 31.0, 29.8, 29.8, 29.7, 29.6, 29.5, 29.3, 29.2, 27.1, 25.6, 25.4, 22.8, 22.5, 20.4, 19.8, 14.3, 14.1 ppm.

**IR** (ATR): $\tilde{\nu}$ = 2925 (s), 2853 (s), 2358 (w), 2340 (w), 2190 (w), 1722 (m), 1684 (m), 1654 (m), 1602 (w), 1540 (w), 1453 (m), 1378 (m), 1352 (m), 1272 (s), 1201 (m), 1177 (m), 1157 (m), 1120 (s), 1076 (s), 1032 (s), 971 (m), 906 (m), 869 (m), 844 (m), 814 (m), 712 (m) cm$^{-1}$.

**HRMS** (ESI): calcd. for $C_{50}H_{72}N_3O_5^+$: 795.5500 [M+H]$^+$
found: 795.5502 [M+H]$^+$.

**4-(4-((*E*)-(4-butylphenyl)diazenyl)phenyl)-N-((2*S*,3*R*,*E*)-1-hydroxy-3-((tetrahydro-2*H*-pyran-2-yl)oxy)octadec-4-en-2-yl)butanamide (Azo-THP-Cer)**

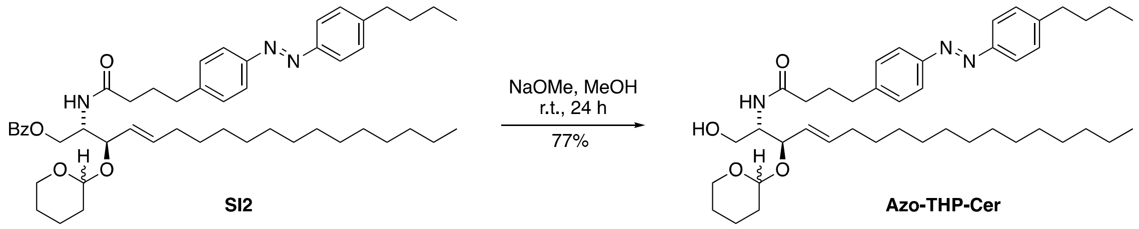

To a solution of fully protected **SI2** (30.3 mg, 38.2 µmol, 1.0 eq.) in dry MeOH (2 mL) was added catalytic sodium methoxide in MeOH (prepared from 1 mL of MeOH and 12 mg MeONa, 200 µL used) and the reaction was stirred for 24 h at room temperature. Then, $H_2O$ (10 mL) was added and the aqueous layer was extracted with EtOAc (3 × 20 mL) and dried ($Na_2SO_4$). Purification *via* flash column chromatography [PE/EtOAc, 6:1 to 3:1] afforded a diastereomeric mixture of **Azo-THP-Cer** (20.3 mg, 29.4 µmol, 77%) as a light orange solid.

$\mathbf{R}_f$ = 0.38 [PE: EtOAc 1:2]

**$^1$H NMR** (400 MHz, $CDCl_3$): $\delta$ = 7.82 (dq, *J* = 8.7, 2.1 Hz, 4H), 7.35–7.28 (m, 4H), 6.42 (d, *J* = 7.6 Hz, 1H), 5.76–5.66 (m, 1H), 5.38 (ddt, *J* = 15.4, 7.1, 1.6 Hz, 1H), 4.45–4.38 (m, 1H), 4.19 (dd, *J* = 7.1, 4.8 Hz, 1H), 4.02–3.91 (m, 2H), 3.88–3.80 (m, 1H), 3.63 (q, J = 6.9, 6.4 Hz, 1H), 3.46 (dp, J = 13.4, 6.3, 5.6 Hz, 1H), 3.26 (d, J = 8.8 Hz, 1H), 2.71 (dt, J = 19.5, 7.7 Hz, 4H), 2.23 (td, J = 7.2, 2.4 Hz, 2H), 2.03 (dtd, J = 10.9, 7.5, 6.4, 3.1 Hz, 4H), 1.85 – 1.70 (m, 1H), 1.70 – 1.59 (m, 2H), 1.59 – 1.42 (m, 4H), 1.42 – 1.17 (m, 25H), 0.94 (t, J = 7.3 Hz, 3H), 0.89 – 0.85 (m, 3H) ppm.

**$^{13}$C NMR** (101 MHz, $CDCl_3$): $\delta$ = 172.8, 151.4, 151.1, 146.4, 144.9, 136.0, 129.3, 129.2, 126.5, 123.0, 122.9, 98.3, 79.1, 64.8, 62.6, 54.0, 36.0, 35.7, 35.2, 33.6, 32.4, 32.1, 31.2, 29.8, 29.8, 29.8, 29.6, 29.6, 29.5, 29.3, 29.2, 27.1, 25.3, 22.8, 22.5, 21.1, 14.3, 14.1 ppm.

**IR** (ATR): $\tilde{\nu}$ = 3303 (bm), 2924 (s), 2853 (m), 1645 (m), 1602 (w), 1546 (w), 1499 (w), 1466 (m), 1378 (w), 1184 (w), 1118 (m), 1074 (m), 1022 (m), 970 (m), 844 (m) $cm^{-1}$.

| | | |
|---|---|---|
| **HRMS** (EI): | calcd. for $C_{43}H_{68}N_3O_4^+$: | 690.5204 $[M+H]^+$ |
| | found: | 690.5208 $[M+H]^+$. |

#### Tetrahydropyrane-protected azosphingomyelin (Azo-THP-SM)

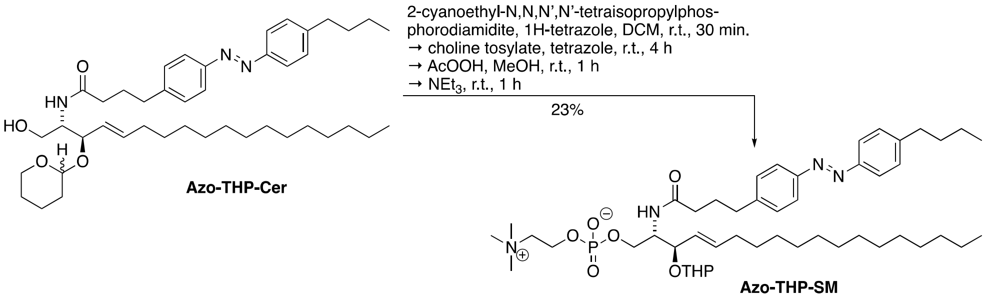

To a solution of a diastereomeric mixture of **Azo-THP-Cer** (26 mg, 37 µmol, 1.0 eq) in dry DCM (1.5 mL) were added 4 Å molecular sieves (0.30 g) and 2-cyanoethyl-*N*,*N*,*N'*,*N'*-tetraisopropylphosphorodiamidite (17 mg, 18 µL, 56 µmol, 1.5 eq.) and 1*H*-tetrazole (0.45 M, 0.10 mL, 44 µmol, 1.2 eq.) at room temperature under Ar. The solution was stirred for 30 min at room temperature. To the reaction mixture was added 1*H*-tetrazole (0.45 M, 0.24 mL, 0.11 mmol, 3.0 eq.), followed by choline tosylate (41 mg, 0.15 mmol, 4.0 eq.) at room temperature. The reaction mixture was stirred for 4 h at room temperature, then MeOH (0.8 mL) and AcOOH (39% in AcOH, 11 µL, 56 µmol, 1.5 eq.) were added, followed by stirring for a further 1 h at room temperature. After that time, 30% aq $NH_3$ (1 mL) was added to the mixture, and the reaction was stirred for 1 h at room temperature. The solution was filtered and concentrated. The product was purified by first elution through TMD-8 resin (THF/water, 90:10), then subsequently by flash column chromatography (silica gel, $CHCl_3$/MeOH/water, 65:25:1 to 65:25:4) to give as **Azo-THP-SM** (7.4 mg, 8.65 mmol, 23%) as an orange solid.

$\mathbf{R_f}$ = 0.28 [$CHCl_3$:MeOH:$H_2O$ 65:25:4]

**$^1$H NMR** (400 MHz, $CD_3OD$): $\delta$ = 7.87 – 7.78 (m, 4H), 7.44 – 7.32 (m, 4H), 5.71 (dt, J = 15.3, 6.6 Hz, 1H), 5.37 – 5.17 (m, 1H), 4.29 – 4.22 (m, 2H), 4.16 – 4.12 (m, 2H), 4.08 – 4.00 (m, 1H), 3.88 (qd, J = 7.2, 6.2, 3.5 Hz, 1H), 3.61 (dd, J = 5.6, 3.7 Hz, 2H), 3.45 (s, 1H), 3.20 (d, J = 2.2 Hz, 9H), 2.72 (q, J = 7.3 Hz, 4H), 2.27 (t, J = 7.5 Hz, 2H), 2.04 – 1.92 (m, 4H), 1.88 – 1.82 (m, 1H), 1.66 (td, J = 7.4, 2.0 Hz, 2H), 1.55 – 1.50 (m, 3H), 1.40 (q, J = 7.5 Hz, 3H), 1.32 – 1.19 (m, 24H), 0.97 (t, J = 7.4 Hz, 3H), 0.88 (t, J = 6.8 Hz, 3H) ppm.

**$^{13}$C NMR** (151 MHz, $CD_3OD$): $\delta$ = 175.4, 152.5, 152.3, 147.8, 146.7, 139.1, 130.4, 130.2, 130.0, 129.9, 127.8, 123.9, 123.8, 121.8, 121.8, 95.8, 76.6, 67.4, 65.7 (d, *J* = 5.5 Hz), 63.5, 60.5 (d, *J* = 4.9 Hz), 54.69, 54.66, 54.2 (d, *J* = 7.6 Hz), 36.8, 36.5, 36.2, 34.8, 33.4, 33.1, 31.8, 30.83, 30.82, 30.80, 30.79, 30.6, 30.5, 30.4, 30.3, 28.9, 26.7, 23.8, 23.4, 20.5, 14.5, 14.3 ppm.

**IR** (ATR): $\tilde{\nu}$ = 3389 (br), 2923 (s), 2851 (m), 2359 (w), 1367 (m)=, 1454 (m), 1247 (m), 1087 (m), 968 (m), 836 (w) cm$^{-1}$.

**HRMS** (EI): calcd. for $C_{44}H_{70}N_3O_8^+$: 855.5759 [M+H]$^+$
found: 855.5752 [M+H]$^+$.

**Azosphingomyelin (Azo-SM)**

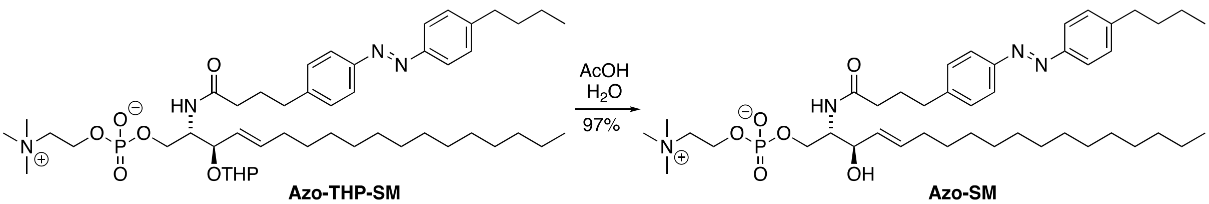

A solution of **Azo-THP-SM** (4.2 mg, 4.91 µmol) in AcOH (0.7 mL) and $H_2O$ (0.35 mL) was stirred for 6 h at 40 °C and concentrated under reduced pressure. Purification of the resulting residue by flash column chromatography ($CHCl_3$:MeOH:$H_2O$ 65:25:4) gave **Azo-SM** (3.7 mg, 4.76 µmol, 97%) as an orange solid.

**R*f*** = 0.18 [$CHCl_3$:MeOH:$H_2O$ 65:25:4]

**$^1$H NMR** (400 MHz, $CD_3OD$): $\delta$ = 7.82 (dd, J = 8.3, 6.4 Hz, 4H), 7.37 (dd, J = 11.7, 8.4 Hz, 4H), 5.70 (dt, J = 15.3, 6.6 Hz, 1H), 5.45 (ddt, J = 15.2, 7.7, 1.5 Hz, 1H), 4.32 – 4.23 (m, 2H), 4.13 – 3.96 (m, 4H), 3.62 (dd, J = 5.3, 3.9 Hz, 2H), 3.21 (s, 9H), 2.71 (td, J = 7.8, 3.9 Hz, 4H), 2.27 (t, J = 7.5 Hz, 2H), 2.00 – 1.90 (m, 4H), 1.70 – 1.62 (m, 2H), 1.40 (q, J = 7.5 Hz, 2H), 1.32 – 1.18 (m, 22H), 0.97 (t, J = 7.4 Hz, 3H), 0.88 (t, J = 6.9 Hz, 3H) ppm.

**$^{13}$C NMR** (151 MHz, $CD_3OD$): $\delta$ = 175.4, 152.5, 152.3, 147.8, 146.7, 135.3, 131.2, 130.3, 130.2, 123.9, 123.8, 72.6, 67.5, 65.8 (d, *J*= 5.0 Hz), 60.4 (d, *J* = 5.0 Hz), 55.3 (d, *J* = 7.4 Hz), 54.7, 54.7, 54.7, 36.8, 36.5, 36.3, 34.8, 33.4, 33.1, 30.8, 30.8, 30.8, 30.8, 30.7, 30.5, 30.5, 30.4, 28.9, 23.8, 23.4, 14.5, 14.3 ppm.

**IR** (ATR): $\tilde{\nu}$ = 3299 (br), 2920 (s), 2850 (m), 1643 (m), 1601 (w), 1556 (w), 1466 (m), 1375 (w), 1229 (m), 1148 (w), 1088 (s), 1048 (s), 968 (s), 838 (m) cm$^{-1}$.

**HRMS** (EI): calcd. for $C_{43}H_{72}N_4O_6P^+$: 771.5184 [M+H]$^+$
found: 771.5183 [M+H]$^+$.

**(2*S*,3*R*,*E*)-2-Azidooctadec-4-ene-1,3-diol (SI3)**

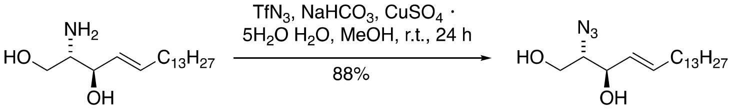

**D-*erythro* sphingosine** **SI3**

D-*erythro*-sphingosine (275 mg, 0.920 mmol, 1 eq.), $NaHCO_3$ (312 mg, 3.72 mg, 4 eq.) and $CuSO_4$•$H_2O$ (8.80 mg, 40.0 µmol, 5 mol%) were dissolved in $H_2O$ (1.2 mL). The emulsion was cooled to 0 °C and freshly prepared $TfN_3$ (2 M in toluene, 2.00 mL, 4.00 mmol, 4.3 eq.) was added. MeOH (2 mL) was added and the reaction was slowly allowed to warm to r.t. After 24 h, $H_2O$ (20 mL) was added and the reaction mixture was extracted with EtOAc (3 × 20 mL), dried ($MgSO_4$) and concentrated under reduced pressure. Flash column chromatography [PE/EtOAc, 10:1 to 0:1] afforded azidosphingosine (**SI3**, 263 mg, 0.806 mmol, 88%) as a yellow oil.

$\mathbf{R_f}$ **=** 0.66 [PE/EtOAc, 1:1].

$[\alpha]_D^{20}$ = –0.14 (*c* = 1, DCM).

**$^1$H NMR** (400 MHz, $CDCl_3$): $\delta$ = 5.86–5.78 (m, 1H), 5.53 (ddt, *J* = 15.4, 7.4, 1.5 Hz, 1H), 4.27–4.23 (m, 1H), 3.78 (dd, *J* = 5.2, 3.9 Hz, 2H), 3.51 (q, *J* = 5.4 Hz, 1H), 2.14–2.00 (m, 2H), 1.47–1.32 (m, 2H), 1.32–1.19 (m, 21H), 0.88 (t, *J* = 6.8 Hz, 3H) ppm.

**$^{13}$C NMR** (101 MHz, $CDCl_3$): $\delta$ = 136.1, 127.9, 73.8, 66.7, 62.6, 32.3, 31.9, 29.7 – 29.2, 28.9, 14.1 ppm.

**IR** (ATR): $\tilde{\nu}$ = 3351 (w), 2919 (s), 2851 (s), 2100 (m), 1669 (w), 1467 (m), 1379 (m), 1266 (m), 1235 (m), 1195 (m), 1154 (m), 1003 (m), 971 (m), 704 (w) cm$^{-1}$.

**HRMS** (EI): calcd. for $C_{18}H_{34}O_2N_3^-$: 324.2657 [M–H]$^-$
found: 324.2658 [M–H]$^-$.

**(2*S*,3*R*,*E*)-2-Azido-1-(trityloxy)octadec-4-en-3-ol (SI4)**

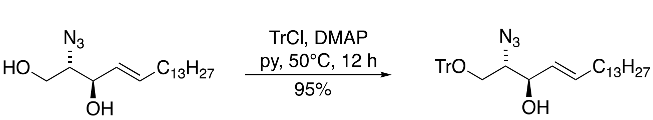

**SI3** **SI4**

Azide **SI3** (263 mg, 0.806 mmol, 1eq.) was dissolved in pyridine (3 mL), then TrCl (274 mg, 0.887 mmol, 1.1 eq.) and DMAP (4.92 mg, 40.3 µmol, 0.05 eq.) were added. The mixture was stirred at 50 °C for 12 h. Afterwards, the solvent was removed under reduced pressure. The crude product was purified by flash column chromatography on silica gel [PE/EtOAc, 10:1 to 3:1] to give protected alcohol **SI4** (437 mg, 0.771 mmol, 95%) as a colorless oil.

**R*f*** = 0.65 [PE/EtOAc, 7:1].

**[α]$_D^{20}$** = 0.01 (*c* = 1, DCM).

**$^1$H NMR** (400 MHz, $CD_3CN$): *δ* = 7.37–7.32 (m, 6H), 7.21 (q, *J* = 6.8, 6.1 Hz, 6H), 7.16 (t, *J* = 7.1 Hz, 3H), 5.46 (dt, *J* = 14.3, 6.8 Hz, 1H), 5.22 (dd, *J* = 15.5, 7.1 Hz, 1H), 3.97 (t, *J* = 6.2 Hz, 1H), 3.49 (dt, *J* = 8.6, 4.4 Hz, 1H), 3.14–3.06 (m, 1H), 3.01 (dd, *J* = 9.9, 7.7 Hz, 1H), 3.14–3.06 (m, 2H), 3.01 (dd, *J* = 9.9, 7.7 Hz, 1H), 1.83 (dq, *J* = 5.2, 2.6 Hz, 2H), 1.01–1.21 (d, *J* = 13.9 Hz, 22H), 0.77 (t, *J* = 6.6 Hz, 3H) ppm.

**$^{13}$C NMR** (101 MHz, $CD_3CN$): *δ* = 144.8, 134.7, 129.9, 129.6, 129.4, 128.1, 87.8, 73.0, 67.2, 64.2, 32.7, 32.6–29.7, 23.3, 14.4 ppm.

**IR** (ATR): $\tilde{\nu}$ = 3422 (bw), 3059 (w), 3033 (w), 2924 (s), 2954 (m), 2362 (w), 2098 (m), 1669 (w), 15098 (w), 1491 (w=, 1448 (m), 1271 (w), 1221 (w), 1184 (w), 1155 (w), 1077 (m), 1033 (w), 1015 (w), 972 (w), 989 (w), 764 (m), 746 (m), 702 (s) cm$^{-1}$.

| | | |
|---|---|---|
| **HRMS** (EI): | calcd. For $C_{37}H_{48}N_3O_2^-$: | 566.3752 [M–H]$^-$ |
| | found: | 566.3746 [M–H]$^-$. |

##### (2*S*,3*R*,*E*)-2-Azido-1-(trityloxy)octadec-4-en-3-yl benzoate (SI5)

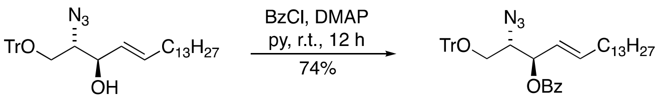

**SI4** **SI5**

To a solution of secondary alcohol **SI4** (437 mg, 0.771 mmol, 1 eq.) in pyridine (12 mL) were added benzoylchloride (0.187 mL, 1.54 mmol, 2 eq.) and DMAP (4.71 mg, 38.6 µmol, 0.05 eq.). The mixture was stirred at room temperature for 12 h. The solvent was removed under reduced pressure and flash column chromatography on silica gel [PE/EtOAc, 1:0 to 2:1] afforded protected D-*erythro*-sphingosine **SI5** (383 mg, 0.570 mmol, 74%) as a colorless oil.

**R*f* =** 0.79 [PE/EtOAc, 7:1]

**[α]$_D$** = –0.003 (*c* = 1, DCM).

**$^1$H NMR** (400 MHz, $CDCl_3$): *δ* = 7.91–7.85 (m, 2H), 7.53–7.45 (m, 1H), 7.39–7.32 (m, 6H), 7.24–7.17 (m, 6H), 7.17–7.11 (m, 3H), 5.74 (dt, *J* = 15.4, 6.7 Hz, 1H), 5.56 (dd, *J* = 7.9, 4.8 Hz, 1H), 5.35 (ddt,

$J$ = 15.4, 7.9, 1.5 Hz, 1H), 3.79 (dt, $J$ = 6.8, 5.0 Hz, 1H), 3.22 (dd, $J$ = 9.8, 6.8 Hz, 1H), 3.12 (dd, $J$ = 9.8, 5.2 Hz, 1H), 1.89 (qt, $J$ = 7.0, 1.7 Hz, 2H), 1.29–1.06 (m, 22H), 0.86–0.74 (m, 3H) ppm.

**$^{13}$C NMR** (101 MHz, $CDCl_3$): $\delta$ = 165.3, 143.6, 138.5, 133.2, 129.9, 129.8, 128.7, 128.5, 128.2, 128.0, 127.3, 123.2, 87.3, 74.9, 64.6, 63.0, 32.4, 32.1, 29.9–29.3, 28.8, 22.9, 14.3 ppm.

**IR** (ATR): $\tilde{\nu}$ = 2924 (m), 2853 (m), 2098 (m), 1723 (m), 1602 (w), 1491 (w), 1466 (w), 1450 (m), 1315 (w), 1263 (s), 1177 (w), 1154 (w), 1092 (m), 1069 (m), 1026 (m), 970 (m), 899 (w), 774 (w), 764 (m), 741 (m), 703 (s) $cm^{-1}$.

| | | |
|---|---|---|
| **HRMS** (ESI): | calcd. for $C_{44}H_{57}N_4O_3^+$ | 689.4425 [$M+NH_4^+$] |
| | found: | 689.4442 [$M+NH_4^+$]. |

##### (2*S*,3*R*,*E*)-2-Azido-1-hydroxyoctadec-4-en-3-yl benzoate (SI6)

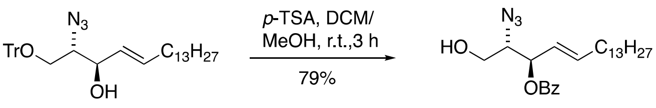

**SI5** **SI6**

To a solution of protected D-*erythro*-sphingosine (**SI5**, 72.6 mg, 0.108 mmol, 1 eq.) in DCM (1 mL) and MeOH (1 mL) was added *p*-toluenesulforic acid hydrate (20.5 mg, 0.108 mmol, 1.0 eq.) and the reaction was stirred at room temperature for 3 h. All volatiles were removed under reduced pressure and the crude product was purified by flash column chromatography [PE/EtOAc, 10:1 to 3:1] to give primary alcohol (**SI6**, 27.6 mg, 85.1 µmol, 79%) as a colorless oil.

**$R_f$** = 0.66 [PE/EtOAc, 4:1].

**$[\alpha]_D^{20}$** = –0.38 ($c$ = 1, DCM).

**$^1$H NMR** (400 MHz, $CDCl_3$): $\delta$ = 8.06 (d, $J$ = 7.7 Hz, 2H), 7.59 (t, $J$ = 7.4 Hz, 1H), 7.46 (t, $J$ = 7.6 Hz, 2H), 5.96 (ddd, $J$ = 13.8, 8.7, 4.3 Hz, 1H), 5.68–5.57 (m, 2H), 3.85–3.70 (m, 2H), 3.63 (dd, $J$ = 11.6, 7.0 Hz, 1H), 2.08 (q, $J$ = 7.1 Hz, 2H), 1.57 (bs, 1H), 1.39 (t, $J$ = 7.2 Hz, 2H), 1.24 (d, $J$ = 3.7 Hz, 16H), 0.88 (t, $J$ = 6.7 Hz, 3H) ppm.

**$^{13}$C NMR** (101 MHz, $CDCl_3$): $\delta$ = 165.5, 138.7, 133.4, 129.8 (C-5), 129.7, 128.5, 123.2, 74.6, 66.2, 62.0, 32.4, 31.9–28.7, 22.7, 14.1 ppm.

**IR** (ATR): $\tilde{\nu}$ = 3428 (bw), 2923 (s), 2853 (s), 2168 (w), 2101 (s), 1722 (s), 1602 (w), 1452 (m), 1316 (m), 1265 (s), 1177 (m), 1110 (s), 1068 (s), 1026 (m), 970 (m), 860 (w), 710 (s), 686 (m) $cm^{-1}$.

| | | |
|---|---|---|
| **HRMS** (EI): | calcd. for $C_{25}H_{43}N_4O_3^-$: | 447.3330 [$M+NH_4^+$] |
| | found: | 447.3336 [$M+NH_4^+$] |

**(2*R*,3*S*,4*S*,5*R*,6*R*)-2-(Acetoxymethyl)-6-(((2*S*,3*R*,*E*)-2-azido-3-(benzoyloxy)octadec-4-en-1-yl)oxy)tetrahydro-2*H*-pyran-3,4,5-triyl triacetate (SI8)**

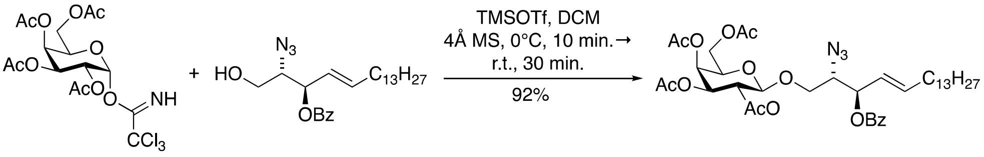

Trichloroacetimidate **SI7**[R1] (225 mg, 0.456 mmol, 2.3 eq.) and acceptor **SI6** (100 mg, 0.198 mmol, 1.0 eq.) were combined and co-evaporated with toluene (3 × 5 mL) and with THF (1 × 5 mL), dried under high vacuum and then dissolved in DCM (0.8 mL). The mixture was stirred with freshly activated 4Å MS at room temperature for 30 min, before the reaction vessel was cooled to 0 °C and TESOTf (0.8 M in DCM, 31.7 µL, 39.6 µmol, 0.2 eq.) was added. After 10 min the reaction was allowed to warm to room temperature and after an additional 30 min the reaction was diluted with DCM and washed with saturated aqueous $NaHCO_3$ solution (10 mL). The aqueous layer was extracted with DCM (3 × 20 mL) and the combined organic phases were dried ($MgSO_4$) and concentrated under reduced pressure. Flash column chromatography (PE/EtOAc, 100:0 to 2:1) afforded protected glycoside (**SI8**, 139 mg, 0.182 mmol, 92%) as a colorless oil.

**$R_f$ =** 0.50 [PE/EtOAc, 2:1].

**$[\alpha]_D^{20}$** = –0.13 (*c* = 1, DCM)

**$^1$H NMR** (600 MHz, $CDCl_3$): *δ* = 8.08–8.02 (m, 2H), 7.57 (t, *J* = 7.4 Hz, 1H), 7.45 (t, *J* = 7.6 Hz, 2H), 5.93 (dt, *J* = 13.6, 6.7 Hz, 1H), 5.63–5.51 (m, 2H), 5.38 (d, J = 3.4 Hz, 1H), 5.23 (dd, *J* = 10.5, 7.9 Hz, 1H), 5.01 (dd, J = 10.4, 3.5 Hz, 1H), 4.49 (d, *J* = 8.0 Hz, 1H), 4.17–4.04 (m, 2H), 3.92 (m, 3H), 3.58 (dd, *J* = 9.1, 4.8 Hz, 1H), 2.15 (s, 3H), 2.10 (s, 3H), 2.09–2.03 (m, 3H), 2.02 (s, 3H), 1.98 (s, 4H), 1.37 (q, *J* = 7.0 Hz, 2H), 1.24 (d, *J* = 4.7 Hz, 23H), 0.87 (t, J = 6.7 Hz, 3H).

**$^{13}$C NMR** (101 MHz, $CDCl_3$): *δ* = 170.3, 170.2, 170.1, 169.3, 165.1, 139.1, 133.2, 129.9, 129.7, 128.4, 122.6, 101.0, 74.7, 70.8, 68.5, 68.0, 66.9, 63.5, 61.1, 32.4, 31.9–29.1, 28.7, 22.7, 20.7, 20.7, 20.6, 14.1 ppm.

**IR (ATR)**: $\tilde{\nu}$ = 3428 (w), 3353 (w), 3296 (w), 1926 (m), 2854 (m), 2108 (m), 1726 (s), 1726 (s), 1601 (w), 1452 (w), 1370 (m), 1317 (w), 1252 (s), 1224 (s), 1176 (w), 1070 (m), 1026 (w), 973 (w), 957 (w), 916 (w), 827 (m), 713 (m) $cm^{-1}$.

| | | |
|---|---|---|
| **HRMS (EI)**: | calcd. for $C_{39}H_{61}N_4O_{12}^+$: | 777.4280 $[M+NH_4]^+$ |
| | found: | 777.4297 $[M+NH_4]^+$. |

**4-(4-((*E*)-(4-Butylphenyl)diazenyl)phenyl)-*N*-((2*S*,3*R*,*E*)-3-hydroxy-1-(((2*R*,3*R*,4*S*,5*R*,6*R*)-3,4,5-trihydroxy-6-(hydroxymethyl)tetrahydro-2*H*-pyran-2-yl)oxy)octadec-4-en-2-yl)butanamide (SI9)**

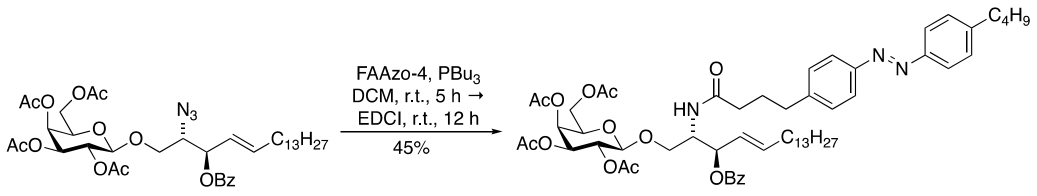

**SI8** **SI9**

Glycoside **SI8** (23.9 mg, 31.5 µmol, 1 eq.) and FAAzo-4[R3] (15.3 mg, 47.2 µmol, 1.5 eq.) were dissolved in DCM (1 mL). $Bu_3P$ (11.6 µL, 47.2 µmol, 1.5 eq.) was added and the reaction stirred for 6 h at room temperature. Then EDCI (22.0 mg, 142 µmol, 3 eq.) was added and the reaction mixture stirred at room temperature for another 12 h. The solvent was removed under reduced pressure and purification *via* flash column chromatography [PE/EtOAc, 10:1 to 0:1] afforded amide **SI9** (14.8 mg, 14.2 µmol, 45%) as a yellow oil.

$\mathbf{R_f}$ **=** 0.60 [PE/EtOAc, 1:2].

**$^1$H NMR** (600 MHz, $CDCl_3$): $\delta$ = 8.05–8.01 (m, 2H), 7.83–7.80 (m, 4H), 7.58–7.53 (m, 1H), 7.46–7.42 (m, 2H), 7.33–7.29 (m, 4H), 5.89 (dtd, $J$ = 15.2, 6.7, 0.8 Hz, 1H), 5.78 (d, $J$ = 9.1 Hz, 1H), 5.59–5.54 (m, 1H), 5.50 (ddt, $J$ = 15.3, 7.6, 1.5 Hz, 1H), 5.35 (dd, $J$ = 3.4, 1.2 Hz, 1H), 5.15 (dd, $J$ = 10.5, 7.8 Hz, 1H), 4.99 (dd, $J$ = 10.5, 3.4 Hz, 1H), 4.54–4.48 (m, 1H), 4.44 (d, $J$ = 7.9 Hz, 1H), 4.07–4.00 (m, 2H), 3.96 (dd, $J$ = 11.3, 6.3 Hz, 1H), 3.85 (ddd, $J$ = 7.4, 6.4, 1.3 Hz, 1H), 3.68 (dd, $J$ = 10.1, 4.3 Hz, 1H), 2.78–2.71 (m, 2H), 2.71–2.65 (m, 3H), 2.10–2.13 (m, 3H), 2.07–1.99 (m, 4H), 1.97 (s, 2H), 1.96 (s, 3H), 1.94 (s, 3H), 1.68–1.61 (m, 2H), 1.38 (h, $J$ = 7.4 Hz, 3H), 1.35–1.17 (m, 27H), 0.94 (t, $J$ = 7.4 Hz, 3H), 0.87 (t, $J$ = 7.1 Hz, 3H) ppm.

**$^{13}$C NMR** (151 MHz, $CDCl_3$): $\delta$ = 172.3, 170.4, 170.3, 170.2, 169.7, 165.4, 151.4, 151.1, 146.5, 144.8, 137.7, 133.2, 129.8, 129.3, 129.2, 128.6, 124.8, 123.0, 122.9, 101.1, 74.5, 70.9, 70.8, 69.0, 67.3, 67.0, 61.2, 51.0, 36.0, 35.7, 35.2, 33.6, 32.5, 32.1, 29.8, 29.6, 29.5, 29.4, 29.1, 27.1, 22.8, 22.5, 20.9, 20.8, 20.7, 14.3, 14.1 ppm.

**IR** (ATR): $\tilde{\nu}$ = 2926 (m), 2854 (w), 1753 (s), 1672 (w), 1602 (w), 1531 (w), 1452 (w), 1369 (m), 1224 (s), 1176 (w), 1071 (m), 968 (w), 846 (w), 714 (m) $cm^{-1}$.

**HRMS** (ESI): calcd. for $C_{59}H_{82}N_3O_{13}^+$: 1040.5842 $[M+H]^+$
found: 1040.5880 $[M+H]^+$.

**(2*R*,3*S*,4*S*,5*R*,6*R*)-2-(Acetoxymethyl)-6-(((2*S*,3*R*,*E*)-3-(benzoyloxy)-2-(4-(4-((*E*)-(4-butylphenyl)-di-azenyl)phenyl)butanamido)octadec-4-en-1-yl)oxy)tetrahydro-2*H*-pyran-3,4,5-triyl triacetate (Azo-β-Gal-Cer)**

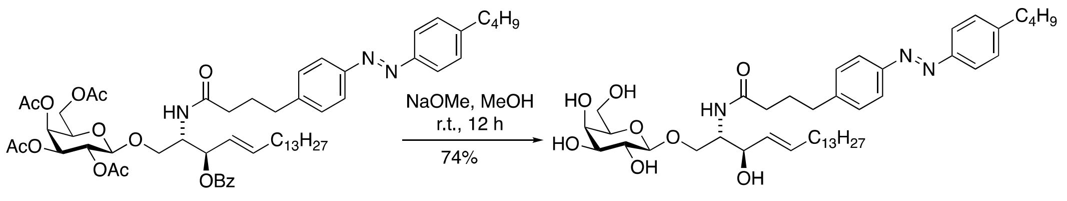

Protected glycosphingolipid (**SI9**, 13.6 mg, 13.1 µmol, 1.0 eq.) was dissolved in MeOH (1.5 mL) and NaOMe was added until pH 9–10. The reaction mixture was stirred at room temperature for 12 h. The reaction was stopped by the addition of DOWEX 50WX 2-100 ($H^+$ form) and stirred for another 30 min at room temperature. All solid material was removed by filtration through a pad of Celite®, which was washed with MeOH (5 mL) and the filtrate was concentrated under reduced pressure. Flash column chromatography [$CHCl_3$/MeOH, 10:1] afforded **Azo-β-Gal-Cer** (7.4 mg, 9.63 µmol, 74%) as yellow viscous oil.

$R_f$ = 0.34 [DCM: MeOH 10:1]

**$^1$H NMR** (800 MHz, $CD_3OD$): $\delta$ = 7.85–7.80 (m, 4H), 7.42–7.33 (m, 4H), 5.69 (dtd, $J$ = 15.3, 6.7, 0.9 Hz, 1H), 5.46 (ddt, $J$ = 15.3, 7.8, 1.5 Hz, 1H), 4.23 (d, J = 7.7 Hz, 1H), 4.18 (dd, J = 10.1, 4.9 Hz, 1H), 4.10 (t, J = 8.1 Hz, 1H), 4.01 (ddd, $J$ = 8.3, 4.9, 3.3 Hz, 1H), 3.83 (dd, $J$ = 3.4, 1.1 Hz, 1H), 3.77 (dd, $J$ = 11.4, 7.0 Hz, 1H), 3.72 (dd, $J$ = 11.4, 5.2 Hz, 1H), 3.63 (dd, $J$ = 10.2, 3.3 Hz, 1H), 3.59–3.55 (m, 1H), 3.55–3.51 (m, 1H), 3.48 (dd, $J$ = 9.7, 3.4 Hz, 1H), 2.76–2.68 (m, 4H), 2.26 (t, $J$ = 7.5 Hz, 2H), 2.02–1.93 (m, 4H), 1.71–1.64 (m, 2H), 1.41 (dt, $J$ = 15.0, 7.4 Hz, 2H), 1.35–1.16 (m, 33H), 0.97 (t, J = 7.4 Hz, 3H), 0.88 (t, $J$ = 7.2 Hz, 3H) ppm.

**$^{13}$C NMR** (101 MHz, $CDCl_3$): $\delta$ = 175.5, 152.5, 152.3, 147.8, 146.7, 135.2, 131.3, 130.3, 130.2, 123.9, 123.8, 105.4 (C-1), 76.8, 74.9, 73.1, 72.7, 70.3, 70.0, 62.6, 54.9, 49.0, 36.8, 36.5, 36.2, 34.8, 33.4, 33.1, 30.8, 30.8, 30.7, 30.5, 30.4, 30.4, 28.8, 23.8, 23.4, 14.5, 14.3 ppm.

**IR** (ATR): $\tilde{\nu}$ = 3288 (bm), 2924 (s), 2852 (m), 2168 (m), 1745 (m), 1558 (w), 1465 (m), 1003 (s), 727 (m) $cm^{-1}$.

| | | |
|---|---|---|
| **HRMS** (EI): | calcd. for $C_{44}H_{70}N_3O_8^+$: | 768.5167 [M+H]$^+$ |
| | found: | 768.5157 [M+H]$^+$. |

### NMR data

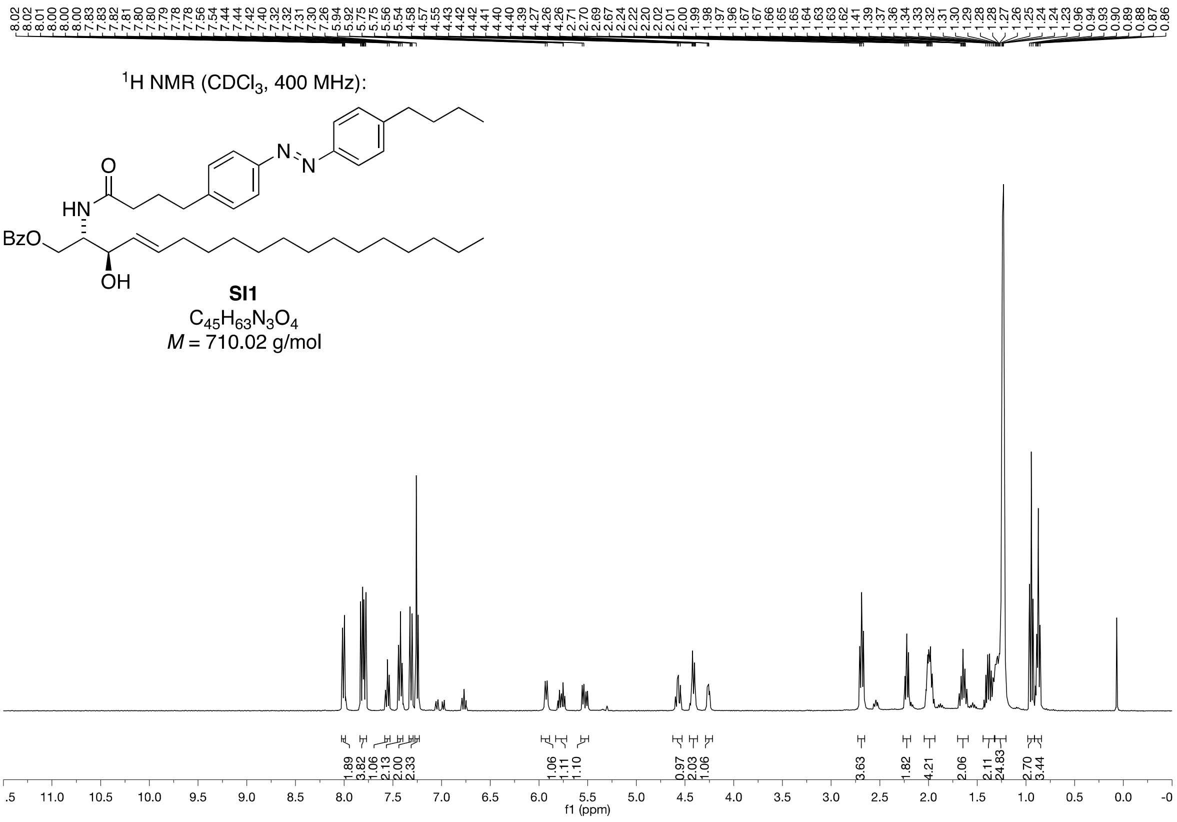
8.02 8.02 8.01 8.00 8.00 7.83 7.82 7.81 7.80 7.79 7.78 7.56 7.54 7.44 7.42 7.40 7.32 7.31 7.30 7.26 5.94 5.92 5.75 5.75 5.56 5.54 4.58 4.57 4.55 4.43 4.42 4.41 4.40 4.39 4.27 4.26 4.26 2.71 2.70 2.69 2.67 2.24 2.22 2.20 2.02 2.01 2.00 1.99 1.98 1.97 1.96 1.67 1.67 1.66 1.65 1.65 1.64 1.63 1.63 1.62 1.41 1.39 1.37 1.36 1.34 1.33 1.32 1.31 1.30 1.29 1.28 1.28 1.27 1.26 1.25 1.24 1.24 1.23 0.96 0.94 0.93 0.90 0.89 0.88 0.87 0.86
$^{1}$H NMR (CDCl$_3$, 400 MHz):
O
HN
N
N
BzO
OH
SI1
$C_{45}H_{63}N_3O_4$
*M* = 710.02 g/mol
1.89 3.82 1.06 2.13 2.00 2.33 1.06 1.11 1.10 0.97 2.03 1.06 3.63 1.82 4.21 2.06 2.11 24.83 2.70 3.44
.5 11.0 10.5 10.0 9.5 9.0 8.5 8.0 7.5 7.0 6.5 6.0 5.5 5.0 4.5 4.0 3.5 3.0 2.5 2.0 1.5 1.0 0.5 0.0 -0
f1 (ppm)

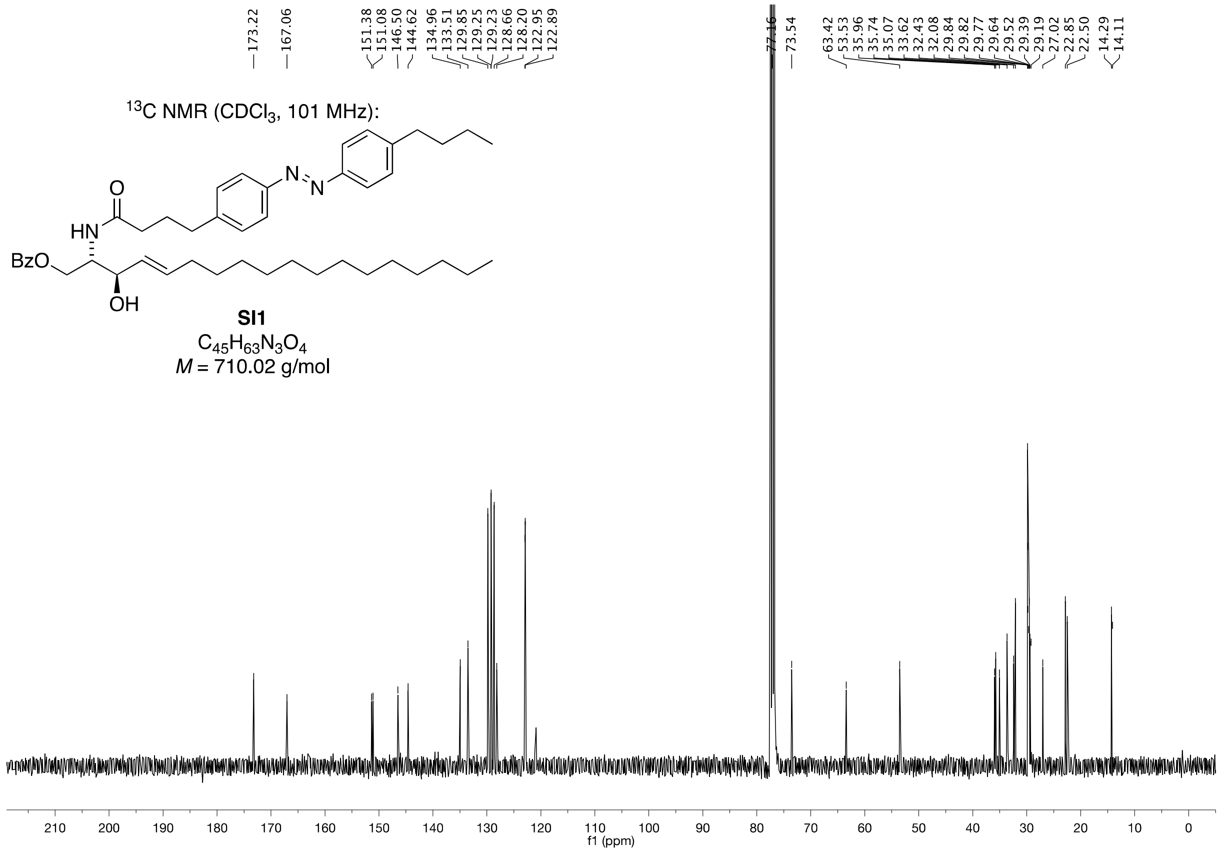
173.22 167.06 151.38 151.08 146.50 144.62 134.96 133.51 129.85 129.25 129.23 128.66 128.20 122.95 122.89 77.16 73.54 63.42 53.53 35.96 35.74 35.07 33.62 32.43 32.08 29.84 29.82 29.77 29.64 29.52 29.39 29.19 27.02 22.85 22.50 14.29 14.11
$^{13}$C NMR (CDCl$_3$, 101 MHz):
O
HN
N
N
BzO
OH
SI1
$C_{45}H_{63}N_3O_4$
*M* = 710.02 g/mol
210 200 190 180 170 160 150 140 130 120 110 100 90 80 70 60 50 40 30 20 10 0
f1 (ppm)

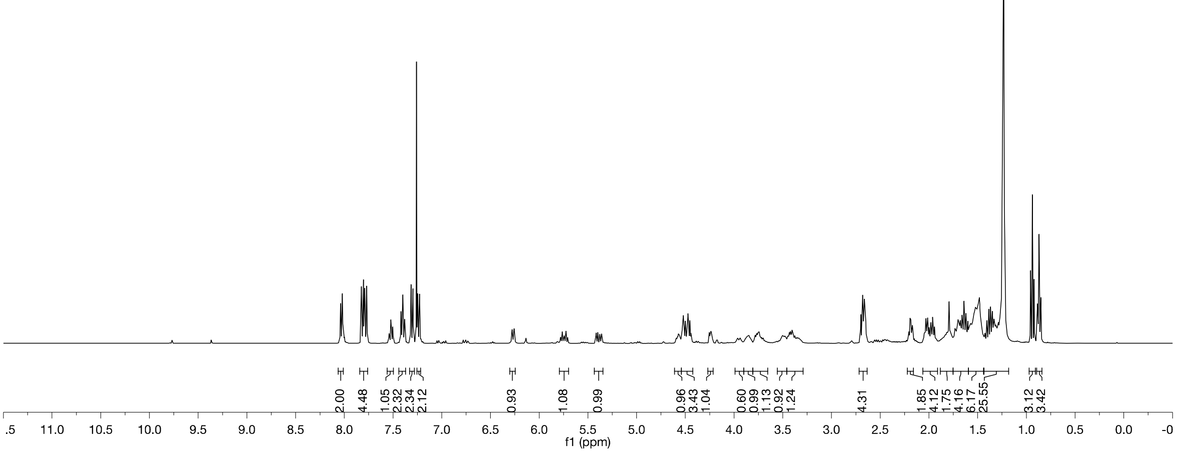

$^{1}H$ NMR ($CDCl_3$, 400 MHz):

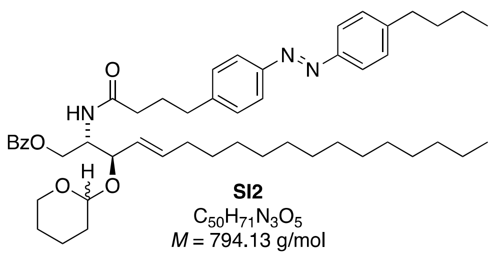

**SI2**
$C_{50}H_{71}N_3O_5$
*M* = 794.13 g/mol

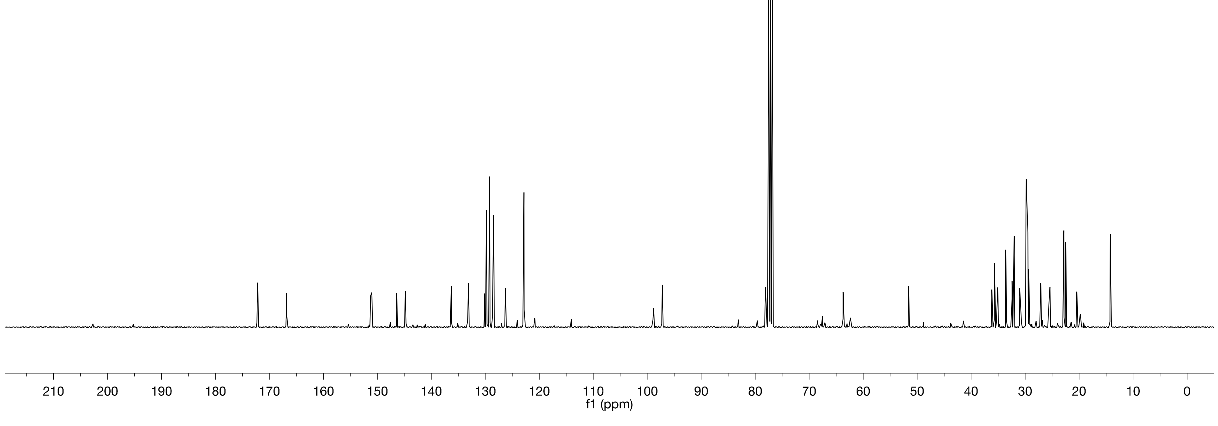

$^{13}C$ NMR ($CDCl_3$, 101 MHz):

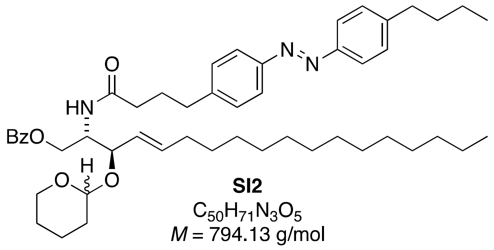

**SI2**
$C_{50}H_{71}N_3O_5$
*M* = 794.13 g/mol

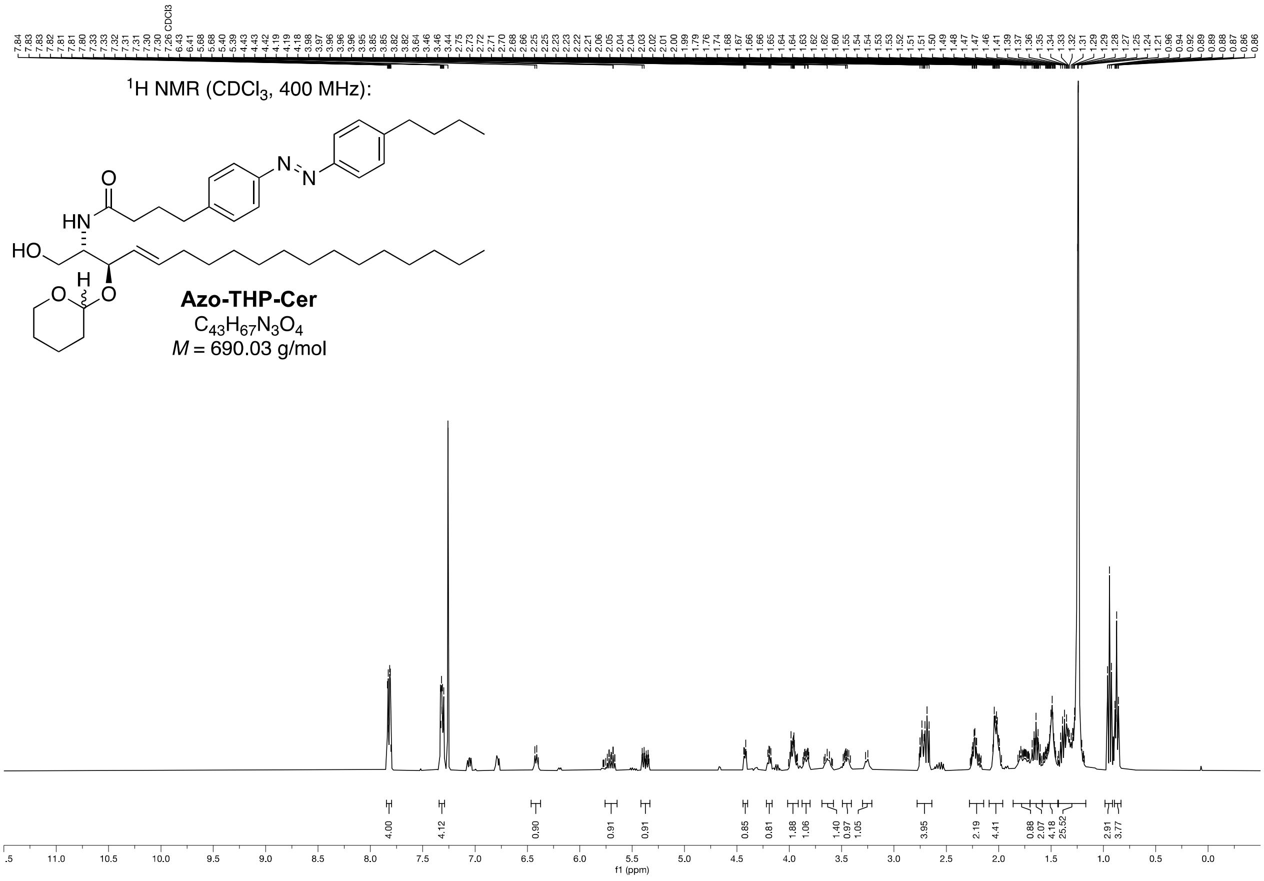

$^{1}H$ NMR ($CDCl_3$, 400 MHz):
O
HN
HO
H
O
N
N
**Azo-THP-Cer**
$C_{43}H_{67}N_3O_4$
$M$ = 690.03 g/mol
f1 (ppm)

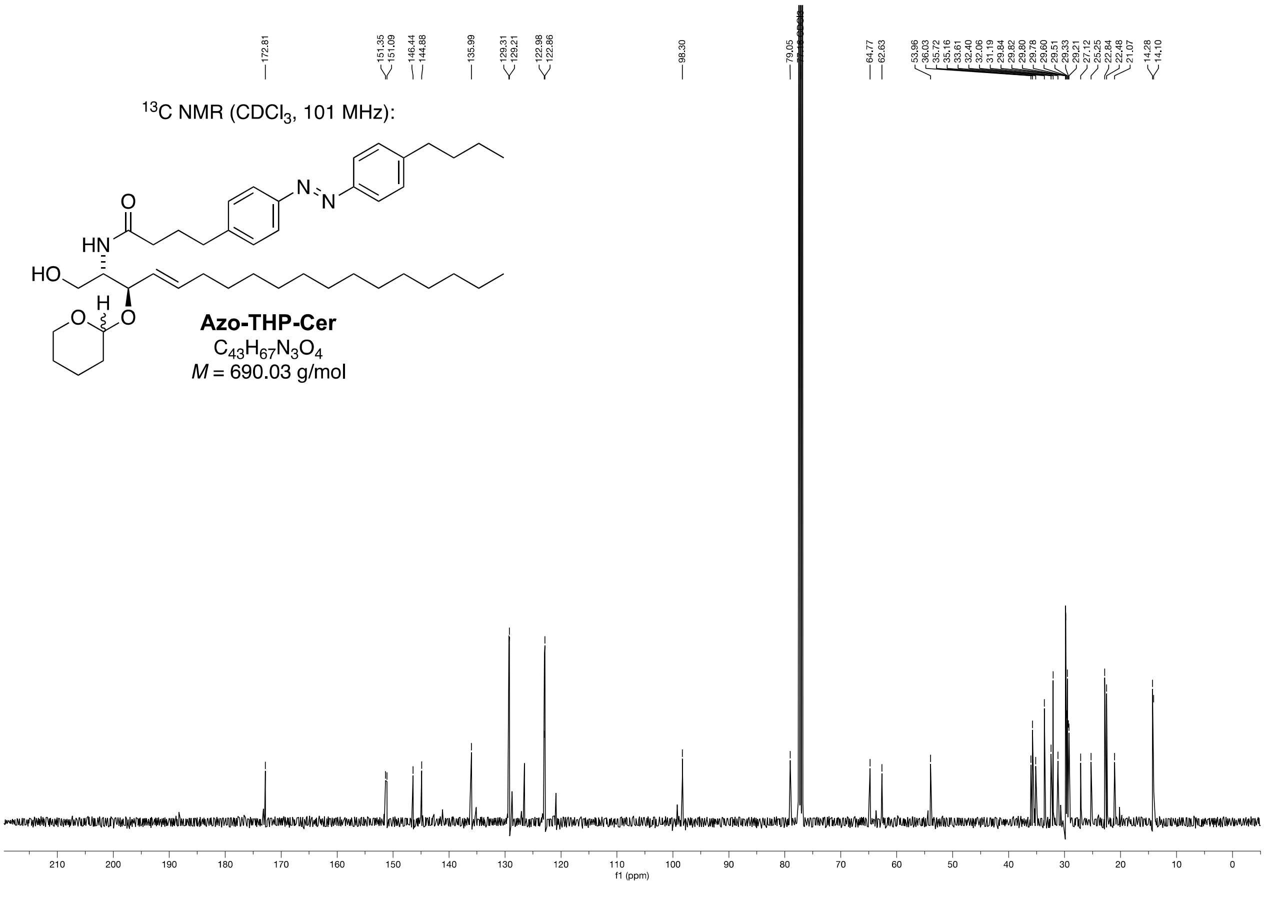

$^{13}C$ NMR ($CDCl_3$, 101 MHz):
O
HN
HO
H
O
N
N
**Azo-THP-Cer**
$C_{43}H_{67}N_3O_4$
$M$ = 690.03 g/mol
f1 (ppm)

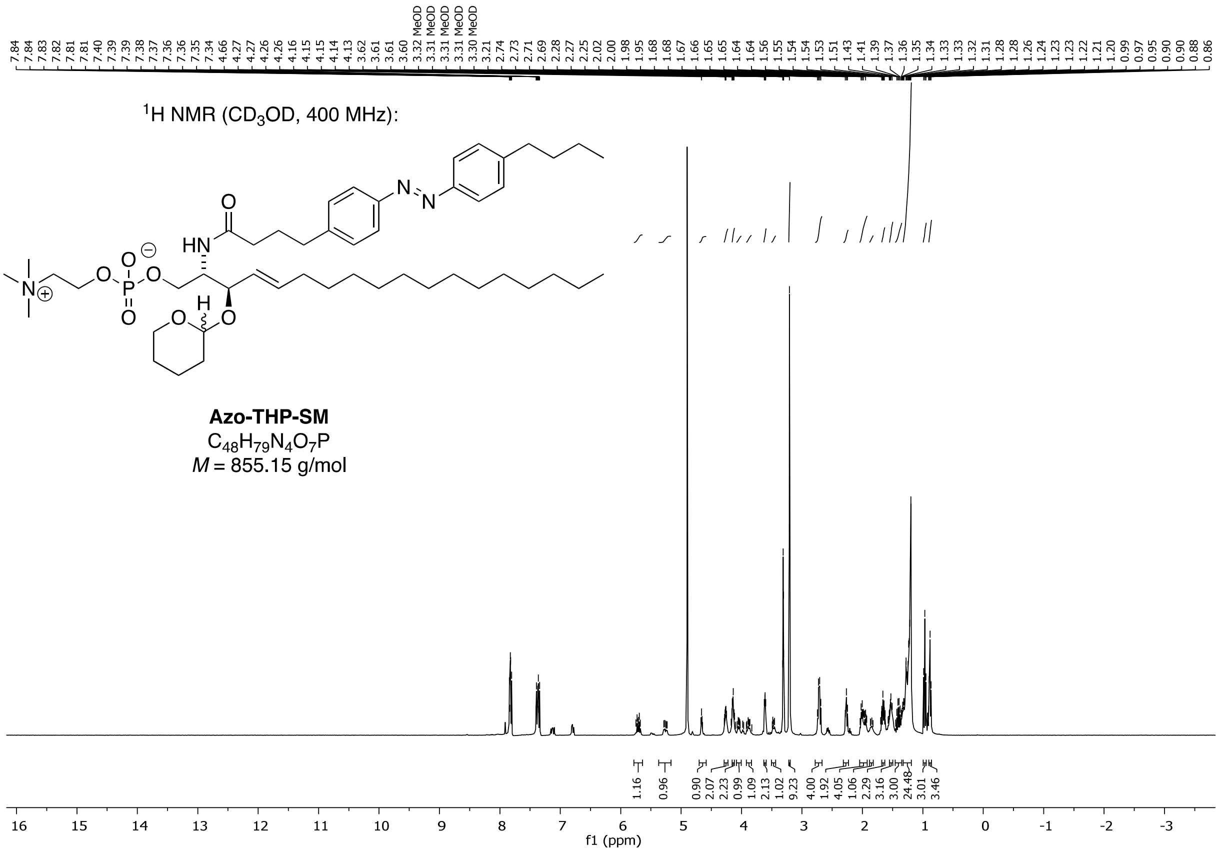

$^1$H NMR (CD$_3$OD, 400 MHz):
Azo-THP-SM
$C_{48}H_{79}N_4O_7P$
$M$ = 855.15 g/mol
f1 (ppm)

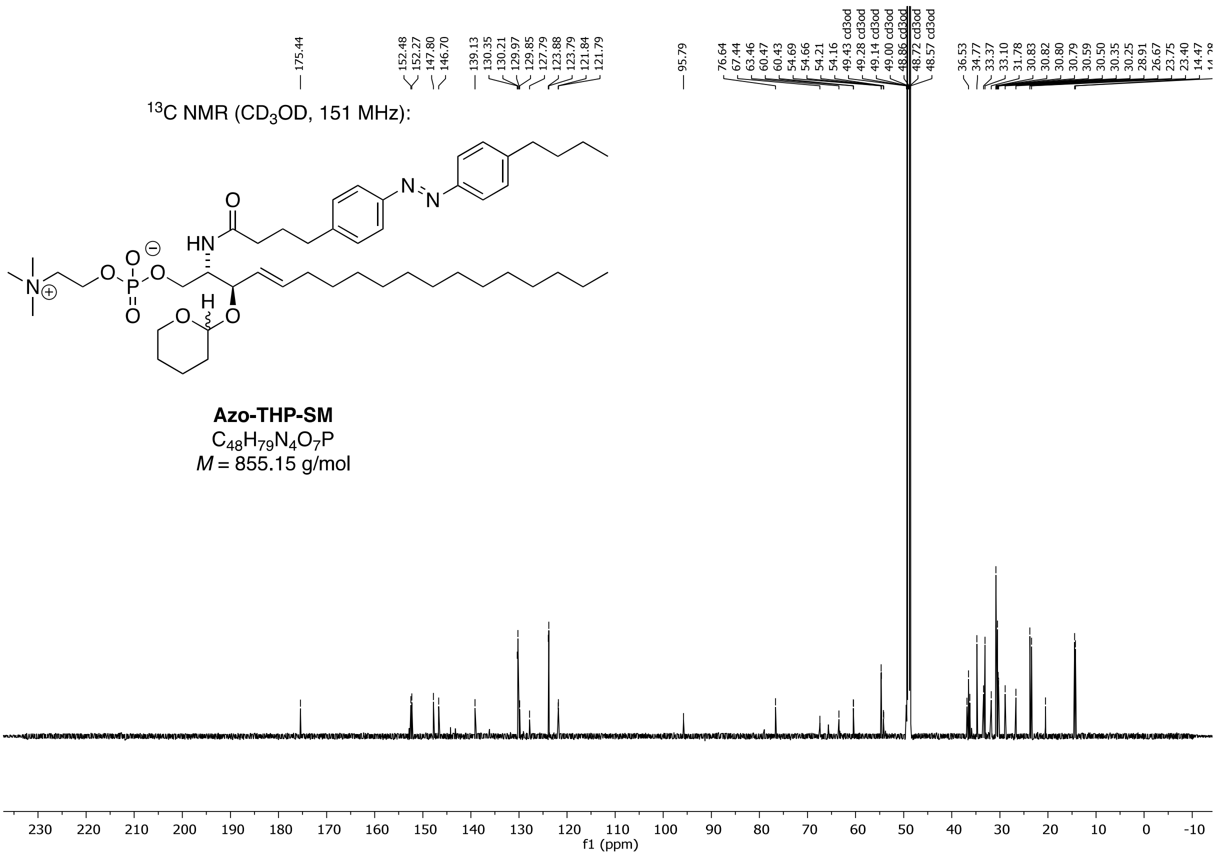

$^{13}$C NMR (CD$_3$OD, 151 MHz):
Azo-THP-SM
$C_{48}H_{79}N_4O_7P$
$M$ = 855.15 g/mol
f1 (ppm)

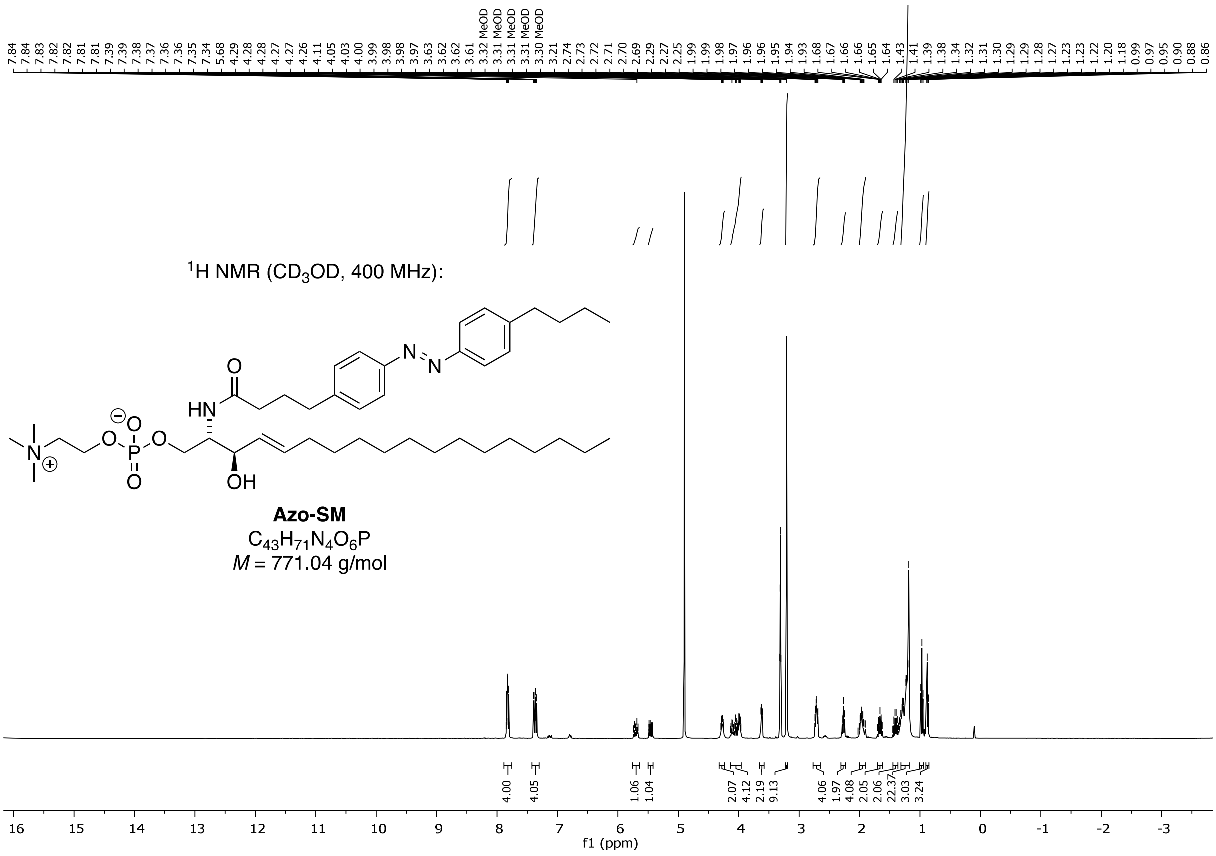
$^1H$ NMR ($CD_3OD$, 400 MHz):
Azo-SM
$C_{43}H_{71}N_4O_6P$
$M$ = 771.04 g/mol
f1 (ppm)

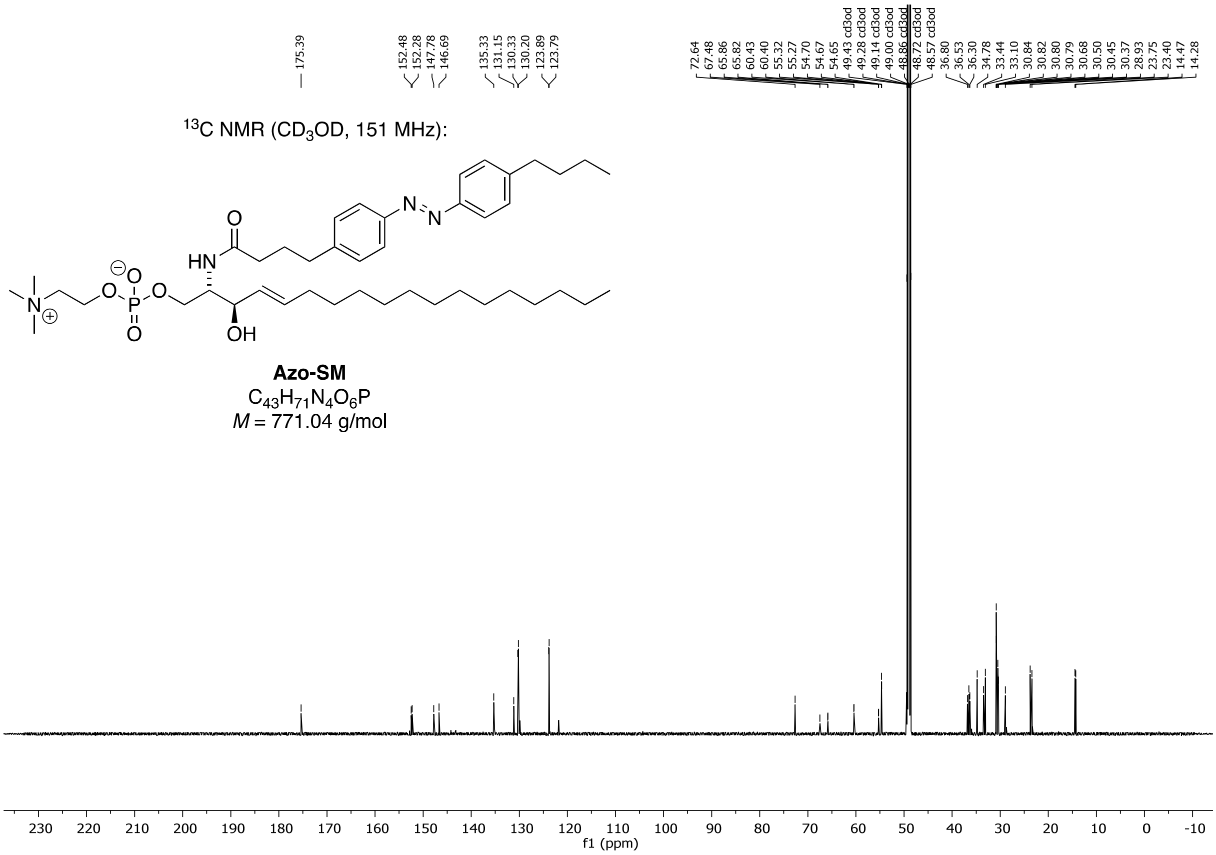
$^{13}C$ NMR ($CD_3OD$, 151 MHz):
Azo-SM
$C_{43}H_{71}N_4O_6P$
$M$ = 771.04 g/mol
f1 (ppm)

[1]H NMR ($CDCl_3$, 400 MHz):

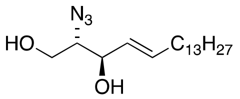

**SI3**
$C_{18}H_{35}N_3O_2$
*M* = 325.50 g/mol

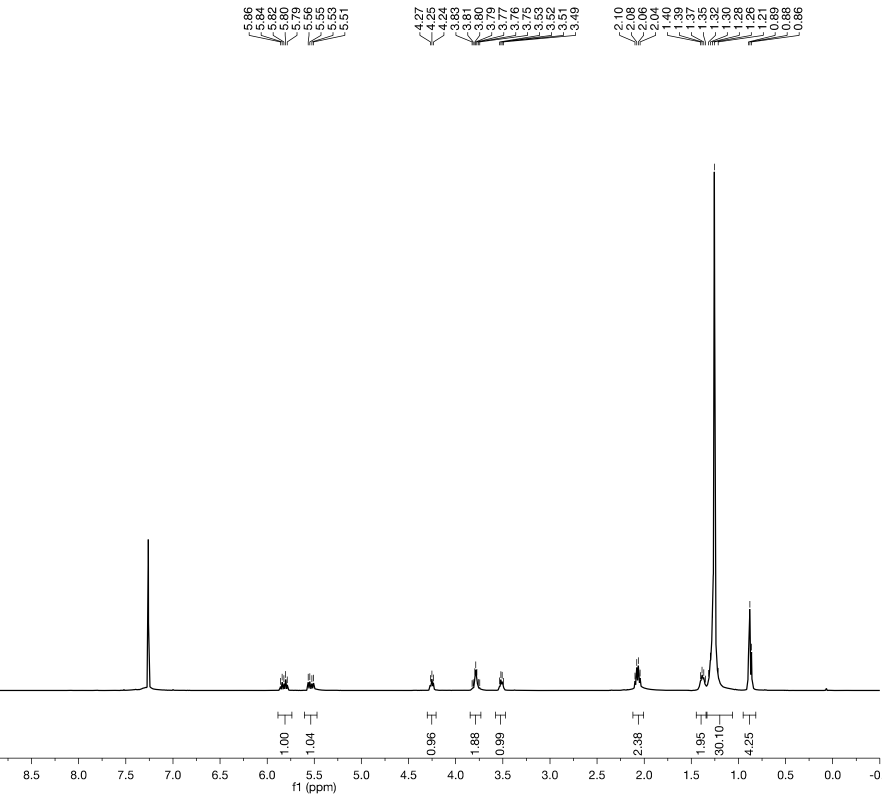

[13]C NMR ($CDCl_3$, 101 MHz):

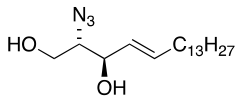

**SI3**
$C_{18}H_{35}N_3O_2$
*M* = 325.50 g/mol

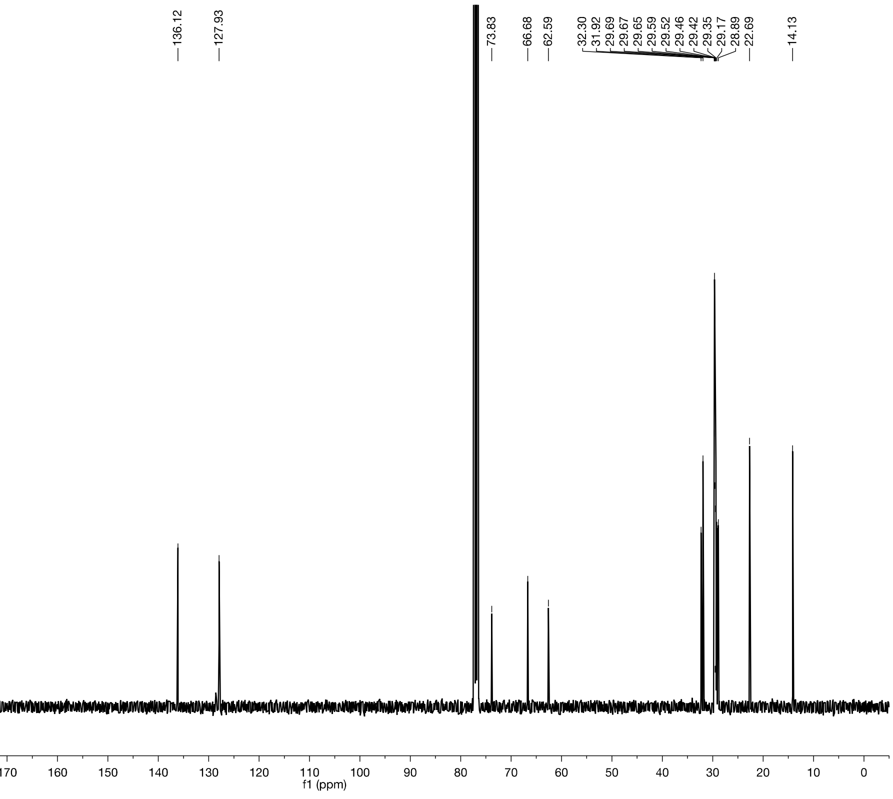

[1]H NMR ($CD_3CN$, 400 MHz):

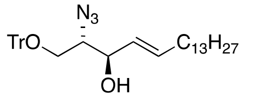

**SI4**

$C_{37}H_{49}N_3O_2$

*M* = 567.82 g/mol

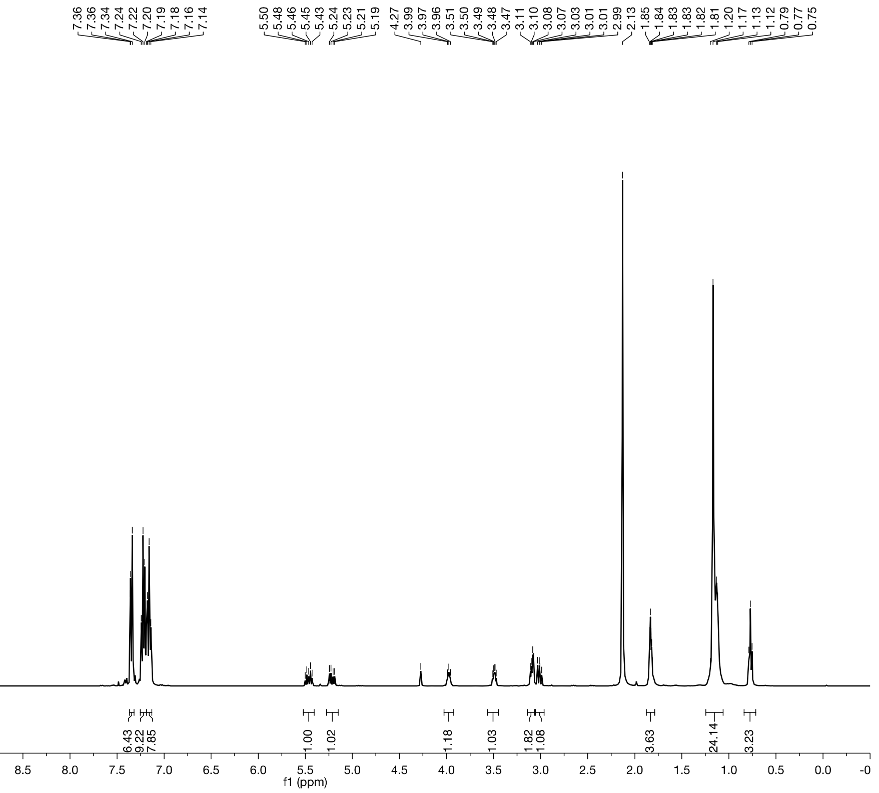

[13]C NMR ($CD_3CN$, 101 MHz):

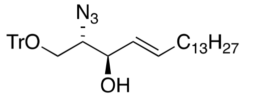

**SI4**

$C_{37}H_{49}N_3O_2$

*M* = 567.82 g/mol

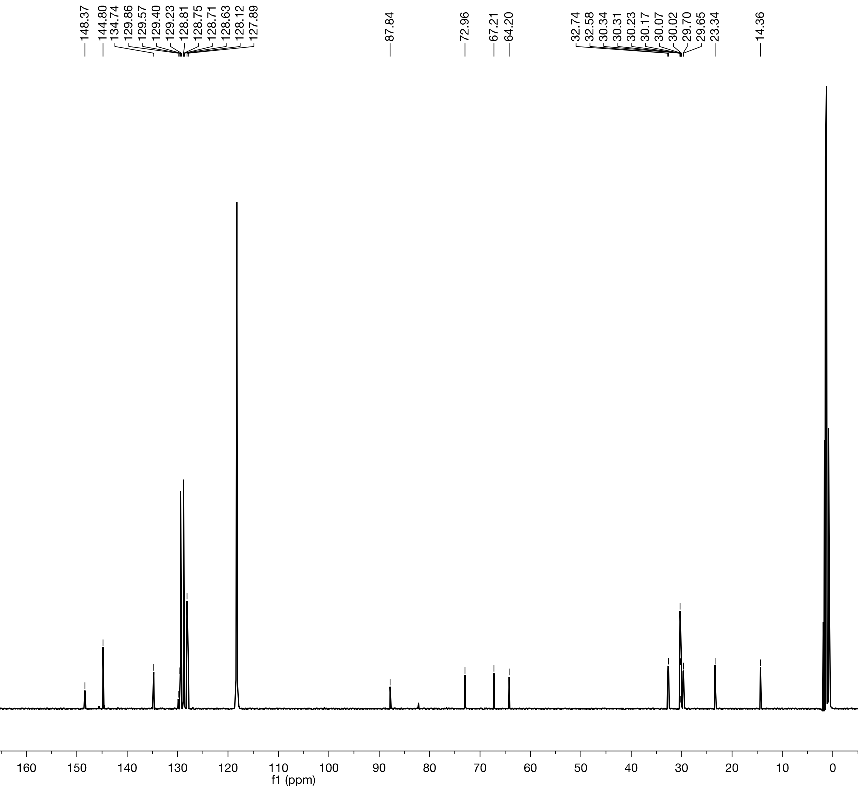

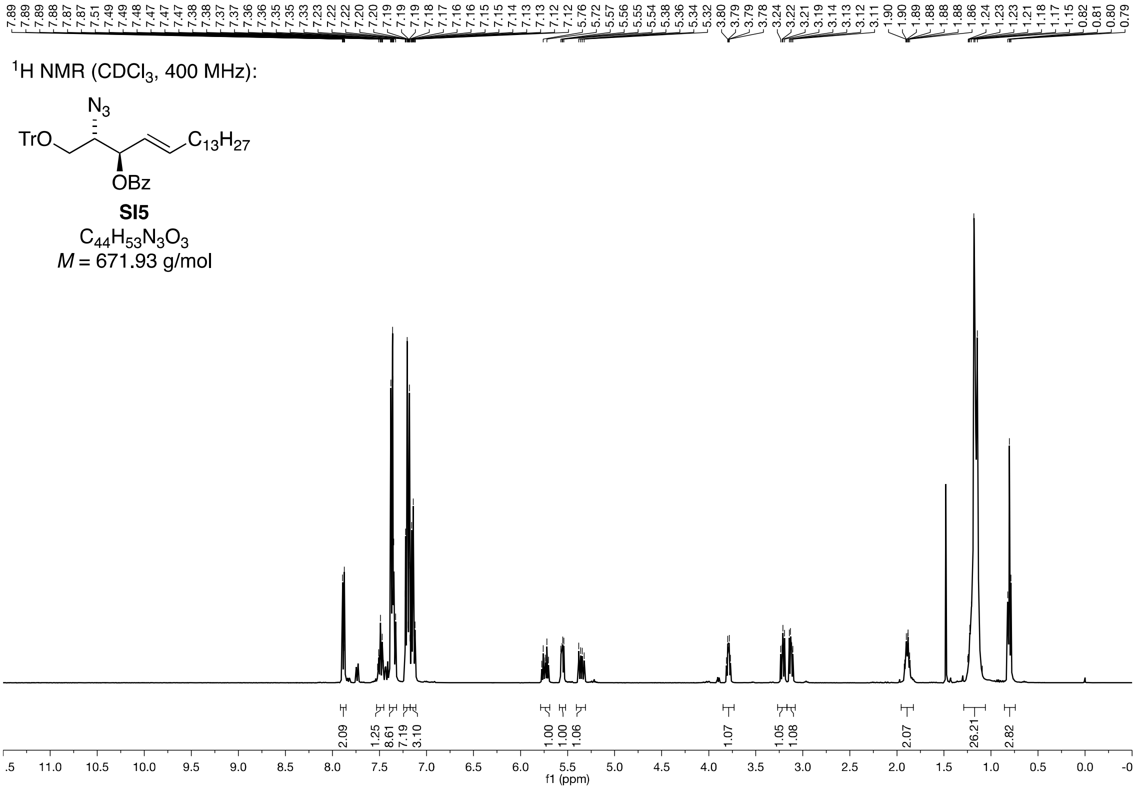
1H NMR (CDCl3, 400 MHz):
N3
TrO
C13H27
OBz
SI5
C44H53N3O3
M = 671.93 g/mol
f1 (ppm)

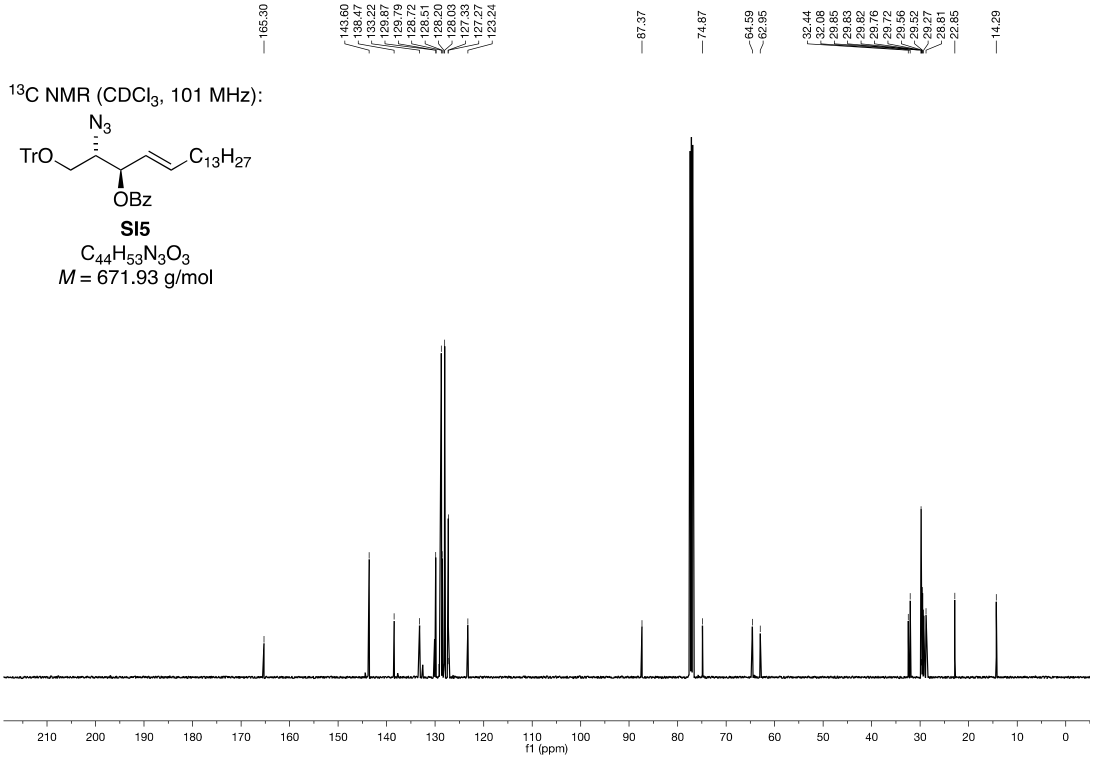
13C NMR (CDCl3, 101 MHz):
N3
TrO
C13H27
OBz
SI5
C44H53N3O3
M = 671.93 g/mol
f1 (ppm)

$^{1}H$ NMR ($CDCl_3$, 400 MHz):

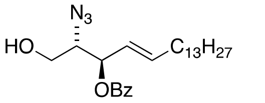

**SI6**

$C_{25}H_{39}N_3O_3$

*M* = 429.61 g/mol

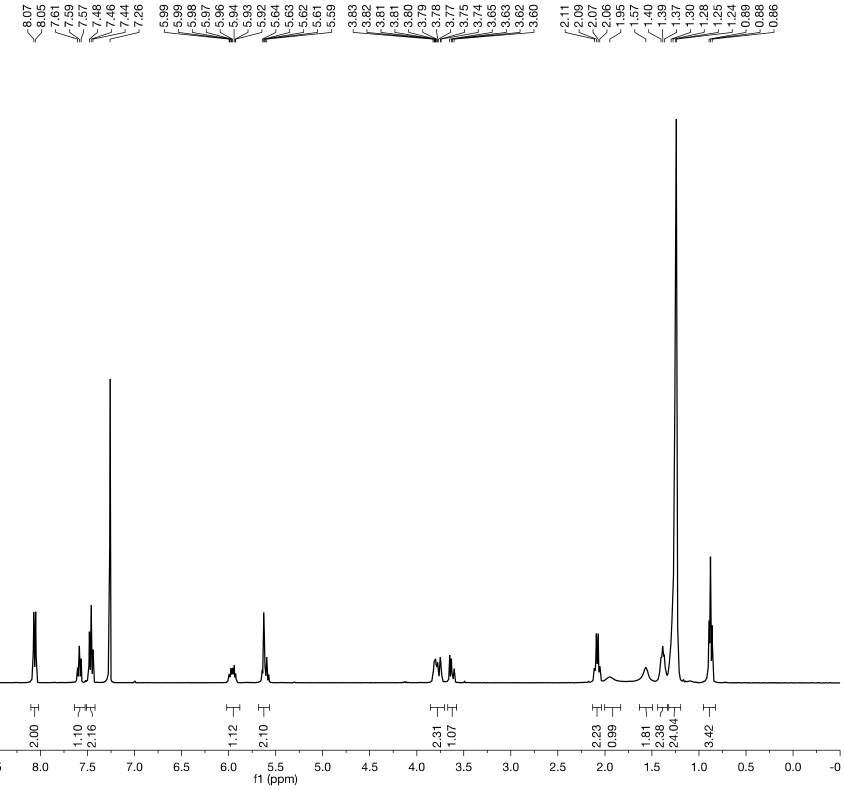

$^{13}C$ NMR ($CDCl_3$, 101 MHz):

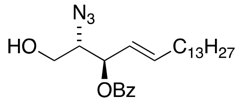

**SI6**

$C_{25}H_{39}N_3O_3$

*M* = 429.61 g/mol

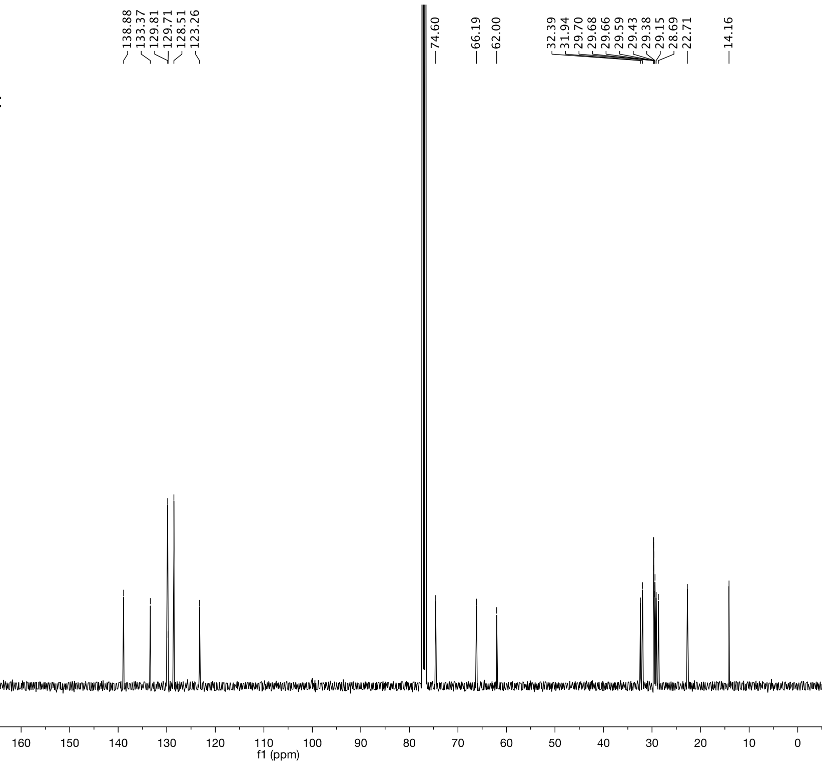

$^{1}H$ NMR ($CDCl_3$, 600 MHz):

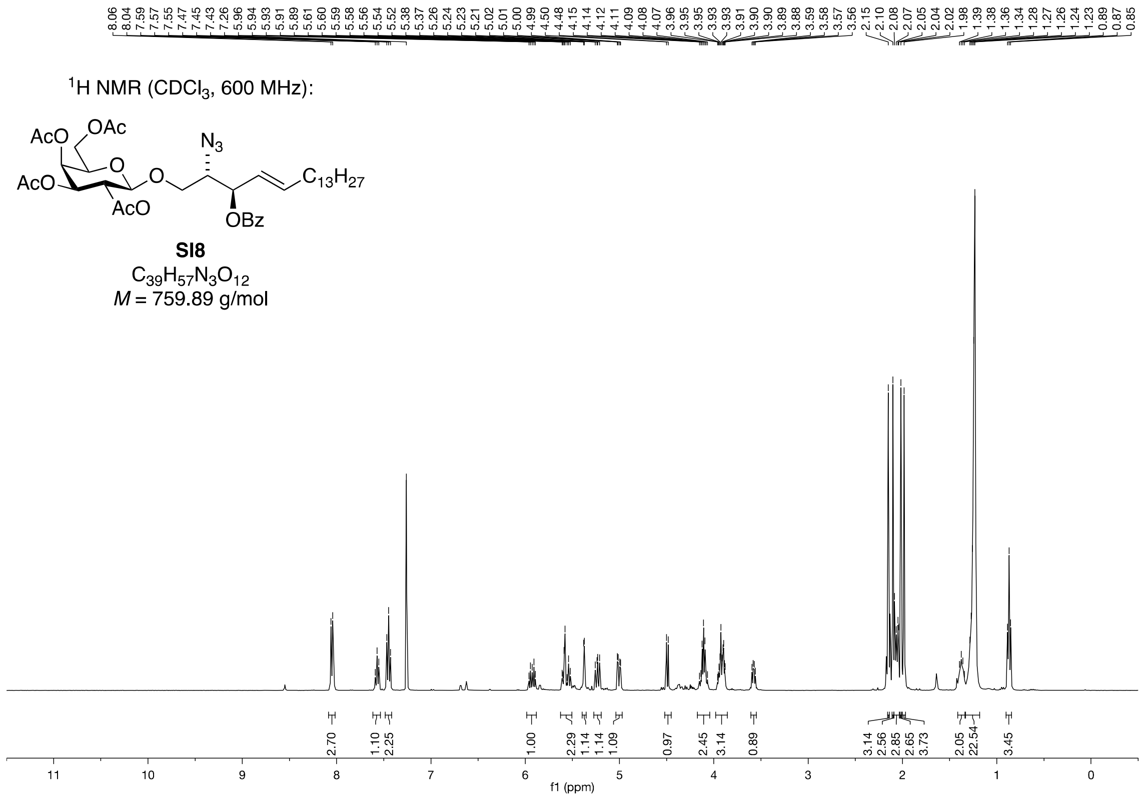

**SI8**

$C_{39}H_{57}N_3O_{12}$

*M* = 759.89 g/mol

$^{13}C$ NMR ($CDCl_3$, 151 MHz):

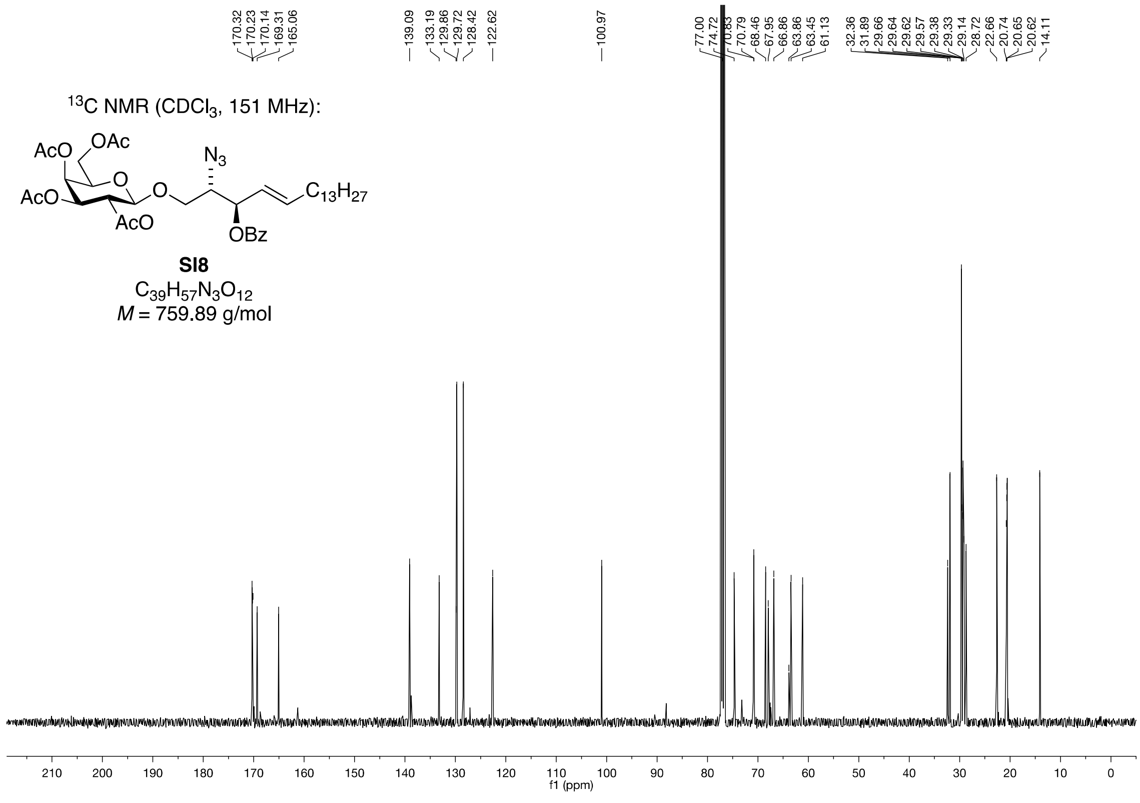

**SI8**

$C_{39}H_{57}N_3O_{12}$

*M* = 759.89 g/mol

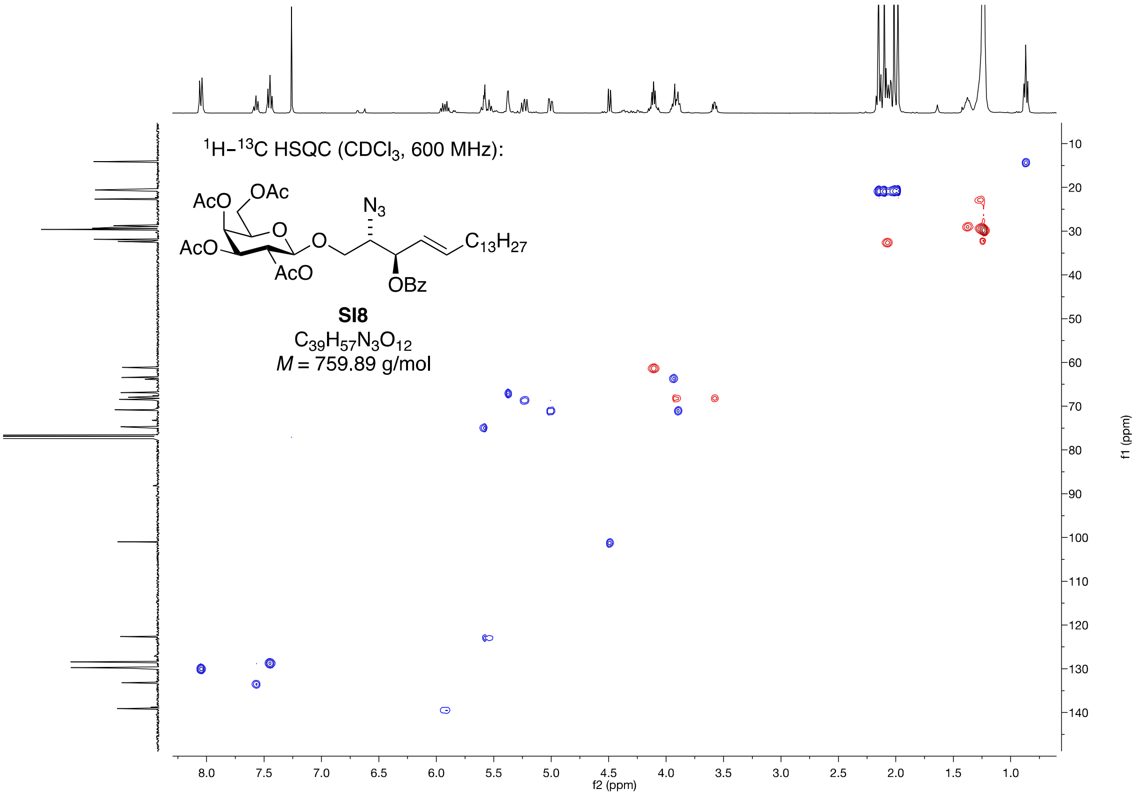

$^1H$–$^{13}C$ HSQC ($CDCl_3$, 600 MHz):
AcO
OAc
AcO
AcO
O
O
$N_3$
OBz
$C_{13}H_{27}$
**SI8**
$C_{39}H_{57}N_3O_{12}$
$M$ = 759.89 g/mol
f2 (ppm)
f1 (ppm)

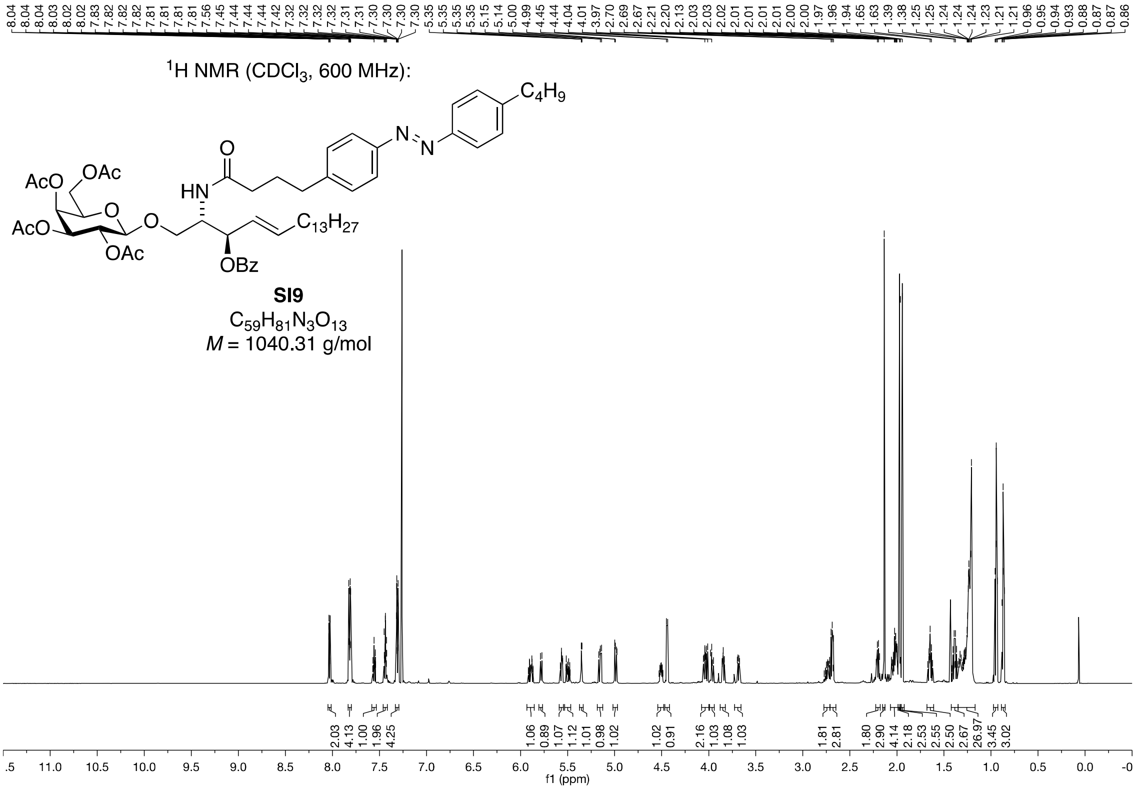
8.04 8.04 8.04 8.03 8.02 8.02 7.83 7.82 7.82 7.82 7.81 7.81 7.81 7.81 7.56 7.45 7.44 7.44 7.42 7.32 7.32 7.32 7.32 7.31 7.31 7.30 7.30 7.30 7.30 5.35 5.35 5.35 5.35 5.15 5.14 5.00 4.99 4.45 4.44 4.04 4.01 3.97 2.70 2.69 2.67 2.21 2.20 2.13 2.03 2.03 2.02 2.01 2.01 2.01 2.01 2.00 1.97 1.96 1.94 1.65 1.63 1.39 1.38 1.25 1.25 1.24 1.24 1.23 1.21 1.21 0.96 0.95 0.94 0.93 0.88 0.87 0.86
$^1$H NMR (CDCl$_3$, 600 MHz):
C$_4$H$_9$
N
N
O
AcO
OAc
HN
O
O
AcO
OAc
C$_{13}$H$_{27}$
OBz
**SI9**
C$_{59}$H$_{81}$N$_3$O$_{13}$
*M* = 1040.31 g/mol
2.03 4.13 1.00 1.96 4.25 1.06 0.89 1.07 1.12 1.01 0.98 1.02 1.02 0.91 2.16 1.03 1.08 1.03 1.81 2.81 1.80 2.90 4.14 2.18 2.53 2.55 2.50 2.67 26.97 3.45 3.02
.5 11.0 10.5 10.0 9.5 9.0 8.5 8.0 7.5 7.0 6.5 6.0 5.5 5.0 4.5 4.0 3.5 3.0 2.5 2.0 1.5 1.0 0.5 0.0 -0
f1 (ppm)

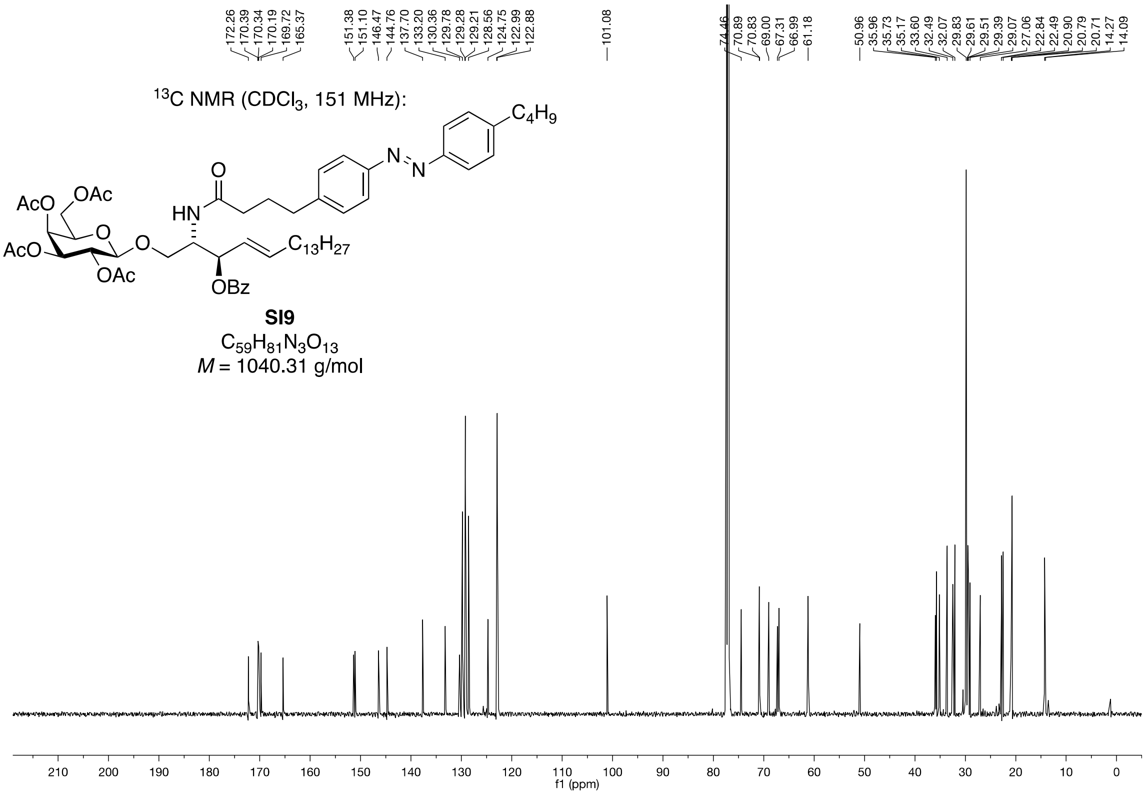
172.26 170.39 170.34 170.19 169.72 165.37 151.38 151.10 146.47 144.76 137.70 133.20 130.36 129.78 129.28 129.21 128.56 124.75 122.99 122.88 101.08 74.46 70.89 70.83 69.00 67.31 66.99 61.18 50.96 35.96 35.73 35.17 33.60 32.49 32.07 29.83 29.61 29.51 29.39 29.07 27.06 22.84 22.49 20.90 20.79 20.71 14.27 14.09
$^{13}$C NMR (CDCl$_3$, 151 MHz):
C$_4$H$_9$
N
N
O
AcO
OAc
HN
O
O
AcO
OAc
C$_{13}$H$_{27}$
OBz
**SI9**
C$_{59}$H$_{81}$N$_3$O$_{13}$
*M* = 1040.31 g/mol
210 200 190 180 170 160 150 140 130 120 110 100 90 80 70 60 50 40 30 20 10 0
f1 (ppm)

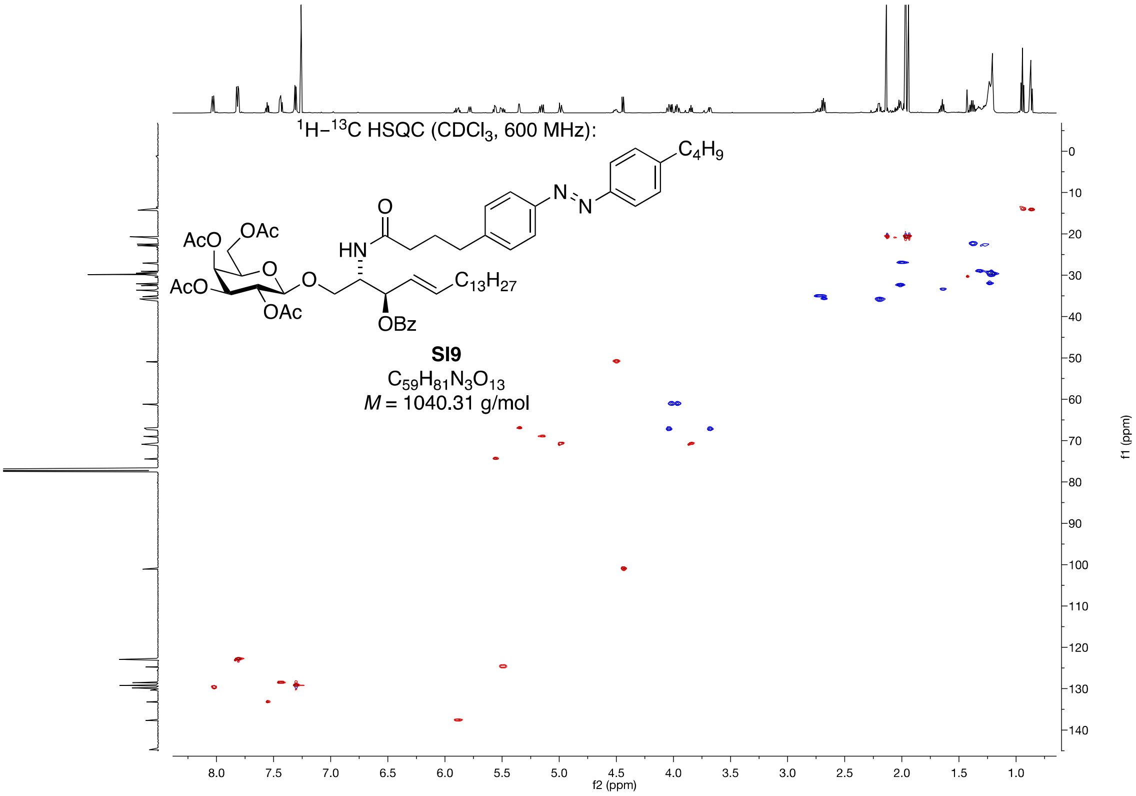

1H–13C HSQC (CDCl3, 600 MHz):
SI9
C59H81N3O13
M = 1040.31 g/mol
f2 (ppm)
f1 (ppm)

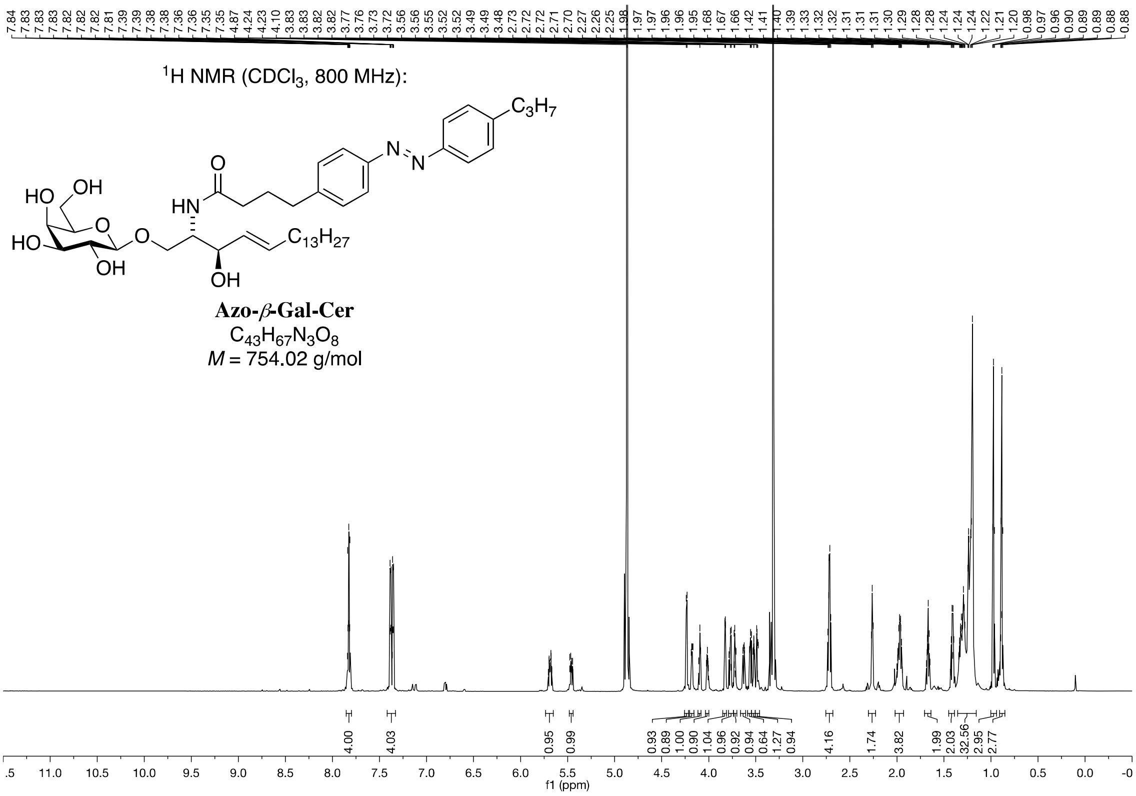
7.84 7.83 7.83 7.83 7.82 7.82 7.81 7.39 7.39 7.38 7.38 7.36 7.36 7.35 4.87 4.24 4.23 4.10 3.83 3.83 3.82 3.82 3.77 3.76 3.73 3.72 3.56 3.56 3.55 3.52 3.52 3.49 3.49 3.48 2.73 2.72 2.72 2.71 2.70 2.27 2.26 2.25 1.98 1.97 1.96 1.96 1.95 1.68 1.67 1.66 1.42 1.41 1.40 1.39 1.33 1.32 1.32 1.31 1.31 1.31 1.30 1.29 1.28 1.28 1.24 1.24 1.22 1.21 1.20 0.98 0.97 0.96 0.90 0.89 0.88 0.88
$^1$H NMR (CDCl$_3$, 800 MHz):
HO
OH
O
HO
OH
O
HN
O
N
N
C$_3$H$_7$
C$_{13}$H$_{27}$
OH
Azo-β-Gal-Cer
C$_{43}$H$_{67}$N$_3$O$_8$
M = 754.02 g/mol
4.00 4.03 0.95 0.99 0.93 0.89 1.00 0.90 1.04 0.96 0.92 0.94 0.64 1.27 0.94 4.16 1.74 3.82 1.99 2.03 32.56 2.95 2.77
.5 11.0 10.5 10.0 9.5 9.0 8.5 8.0 7.5 7.0 6.5 6.0 5.5 5.0 4.5 4.0 3.5 3.0 2.5 2.0 1.5 1.0 0.5 0.0 -0
f1 (ppm)

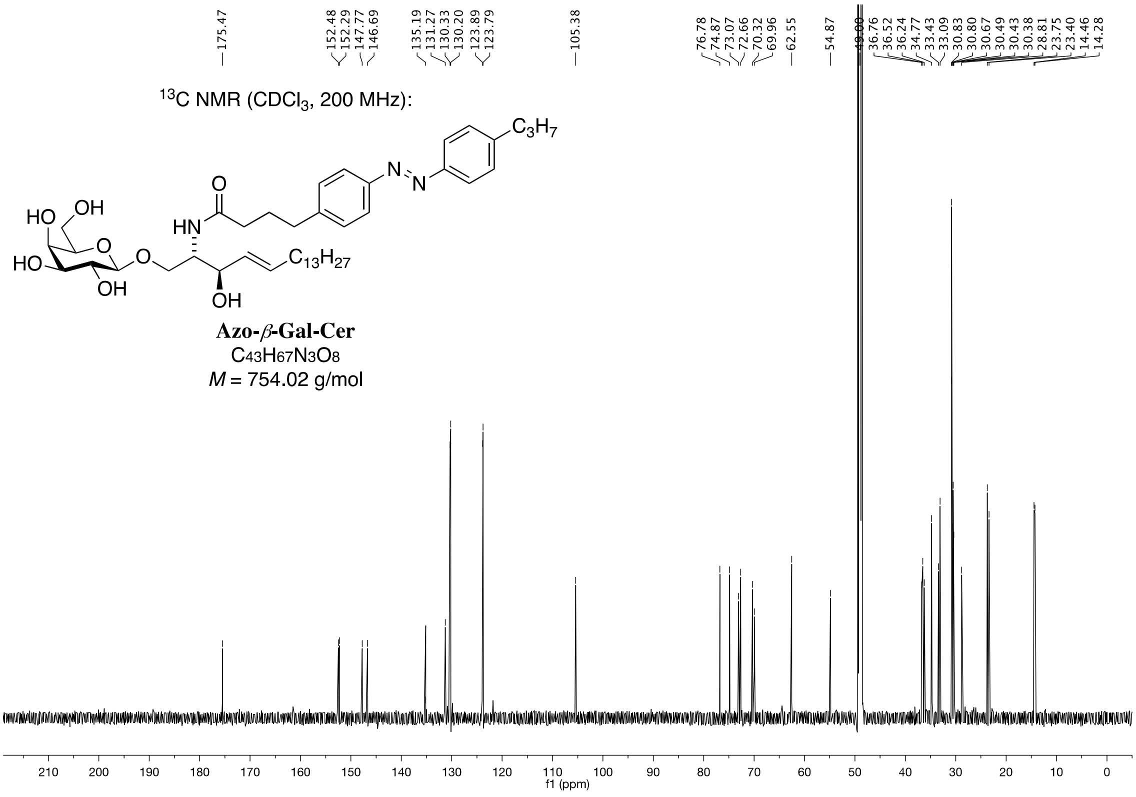
175.47 152.48 152.29 147.77 146.69 135.19 131.27 130.33 130.20 123.89 123.79 105.38 76.78 74.87 73.07 72.66 70.32 69.96 62.55 54.87 49.00 36.76 36.52 36.24 34.77 33.43 33.09 30.83 30.80 30.67 30.49 30.43 30.38 28.81 23.75 23.40 14.46 14.28
$^{13}$C NMR (CDCl$_3$, 200 MHz):
HO
OH
O
HO
OH
O
HN
O
N
N
C$_3$H$_7$
C$_{13}$H$_{27}$
OH
Azo-β-Gal-Cer
C$_{43}$H$_{67}$N$_3$O$_8$
M = 754.02 g/mol
210 200 190 180 170 160 150 140 130 120 110 100 90 80 70 60 50 40 30 20 10 0
f1 (ppm)

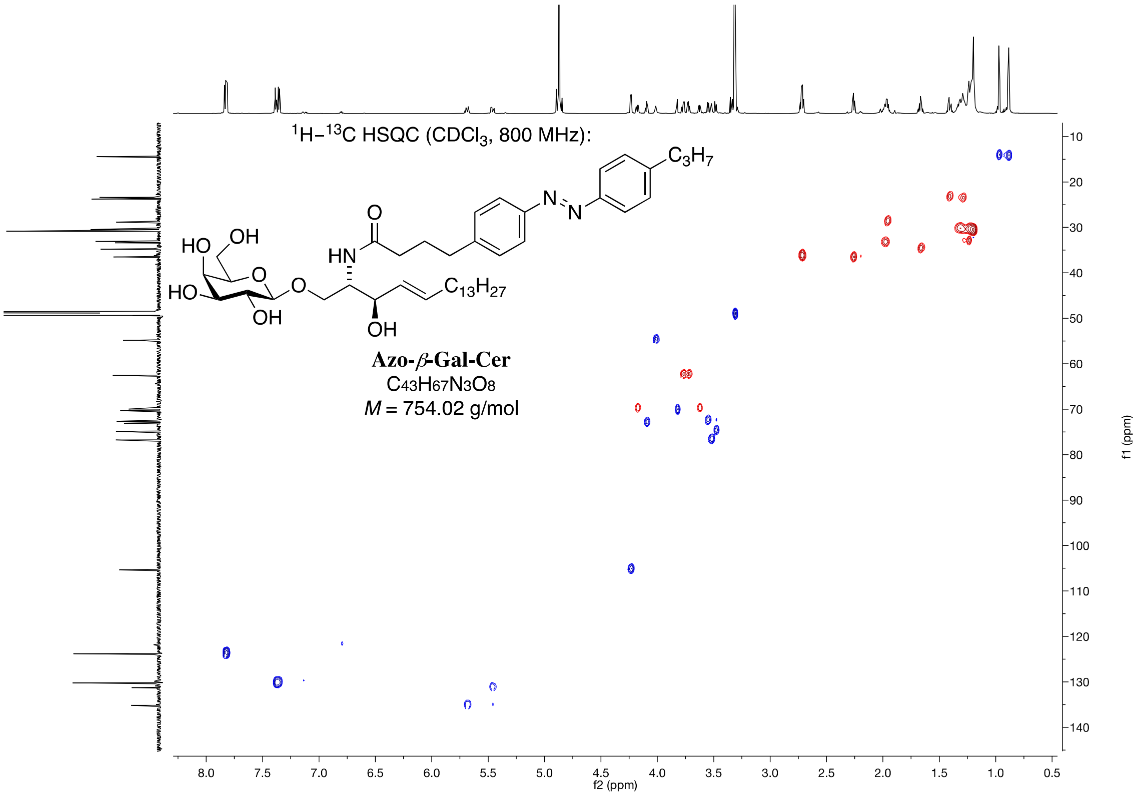

1H–13C HSQC (CDCl3, 800 MHz):
Azo-β-Gal-Cer
C43H67N3O8
M = 754.02 g/mol
f1 (ppm)
f2 (ppm)
